# Multispecies Mixtures: An Individual-Centered Quantitative Genetic Framework for Complex Plant Neighborhoods

**DOI:** 10.64898/2026.05.27.728303

**Authors:** Nicolas Salas, Germain Montazeaud, Peter M. Bourke, Bernadette Julier, Alain Baranger, Jacques David

## Abstract

Mixing crop species or varieties in the same field can raise yield and stability, but performance varies widely among combinations that breeders cannot yet predict. What makes a good neighbor is partly heritable, and quantitative genetics models this as an indirect genetic effect describing a genotype’s contribution to its neighbors’ phenotypes. Here, we extend existing models with a complementary approach based on continuous neighborhoods, in which diverse genotypes of two species are mixed.

In a continuous-neighborhood design, several genotypes of each species are grown in a single trial, with every plant georeferenced. Each plant’s phenotype is decomposed into direct genetic effects, indirect genetic effects from neighbors of both species, and environmental effects. Breeding values are then assembled for each genotype by combining its three genetic effects.

We derived analytical expressions for the variances and covariance of these effects, validated the model, and evaluated its statistical properties through simulations spanning a wide range of parameter settings. For the same number of plants, the continuous-neighborhood design estimated indirect effects far more accurately than a pairwise design and, in addition, allowed indirect environmental variance to be estimated. Applied to a wheat–alfalfa experiment comprising 3,840 plants, from 181 wheat pure lines and 106 alfalfa families, the model attributed 8.6% and 8.2% of the variance in wheat grain number and alfalfa biomass to direct genetic effects respectively, and 4.0% and 4.6% to indirect genetic effects from alfalfa. The framework therefore offers an individual-centered basis for analyzing multispecies neighborhoods and their breeding potential.

**Article Summary:** Growing several crop species or varieties together in the same field can boost yield and stability, but the outcome is unpredictable and the genetic causes remain unclear. We developed a theoretical and statistical framework that links each plant’s performance to its own genes and to those of its neighbors, both from the same and from a different species. Computer simulations of a two-species field showed that these direct and neighbor-driven genetic effects can be reliably separated when enough plants are measured per variety. The framework opens the way to breeding crop mixtures that perform well specifically when grown alongside another species.

## Introduction

Current global intensive agricultural systems have been shown to be unsustainable (Tilman *et al*. 2002), as productivity no longer increases for the most widely cultivated crops (Ray *et al*. 2012), while climate change increases yield uncertainty across time and space (Lesk *et al*. 2016). Agroecology offers a response to these challenges by reintroducing the positive processes observed in natural ecosystems, which are already well adapted to highly variable environments (Malézieux 2012). It relies on increasing diversity in time and space, at both the genotypic and species levels, to favor positive interactions that improve productivity and stability (Yang *et al*. 2019; Tamburini *et al*. 2020; Renard and Tilman 2019). Positive effects of within-field diversity have been documented in variety mixtures (Borg *et al*. 2018; Beillouin *et al*. 2021; Reiss and Drinkwater 2018) and in species mixtures (Brooker *et al*. 2024; Cougnon *et al*. 2022; Mamine and Farès 2020; Brooker *et al*. 2015; Li *et al*. 2023; Stomph *et al*. 2020). However, these effects remain highly variable (Borg *et al*. 2018; Beillouin *et al*. 2021), and defining successful assembly rules for genotypes and crops remains a challenge.

For example, recent advances in precision agriculture now make it possible to grow wheat under a living mulch of alfalfa (**?**), with alfalfa providing multiple ecosystem services to the wheat crop, including nitrogen supply, improved soil structure, and weed suppression (Maamouri *et al*. 2015; Barilli *et al*. 2017; van Bruggen and Finckh 2016). This innovation raises the question of how to improve the performance of this association, first by identifying the best wheat–alfalfa combinations and subsequently by breeding them (El Ghazzal *et al*. 2025). The existence of genetic variation for interaction effects, both within and between species, would have important consequences for the choice of parents used to generate the next generation.

In natural ecosystems, plants interact both intraspecifically, between genotypes of the same species, and interspecifically. These interactions are affected by environmental conditions (Brooker 2006), but the environment is not their only driver, as genetics is also involved (Whitham *et al*. 2003, 2006): in crop communities they are heritable and can evolve through natural or artificial selection (De Lisle *et al*. 2022). Two frameworks have been developed to model them. The trait-based framework (Moore *et al*. 1997) operates at the phenotypic level, describing how the phenotype of an individual is shaped by the phenotypes of its neighbors. It therefore elucidates the ecological mechanisms at work, but only accounts for the part of the indirect effect caused by traits that have been measured (De Lisle *et al*. 2022; Bailey and Desjonquères 2022). The variance-based framework, first proposed by Griffing (1967) and subsequently developed by Bijma et al. (2007) (Griffing 1967, 1968, 1976, 1981; Bijma *et al*. 2007b,a), instead relates the genotype of a neighboring plant to the phenotype of a focal plant through a single statistical term, without specifying the phenotypic pathway through which this effect occurs. Biologically, this effect is indirect, since the neighbor’s genotype acts first on its own phenotype, which in turn affects the focal plant. The framework estimates the total indirect contribution of an individual, whatever the traits that carry it, so that only the response trait needs to be recorded, together with the identity of the neighboring plants (Bijma 2010b; Cappa and Cantet 2008). This suits breeding applications, where ranking and selecting candidates requires knowledge of their total contribution. It is such a framework that we build on here.

In this framework, the phenotype of an individual is decomposed into contributions from its own genotype (direct genetic effect, DGE) and environment (direct environmental effect, DEE), and from the heritable (indirect genetic effect, IGE) and non-heritable (indirect environmental effect, IEE) effects that its neighbours exert on it. We use the simplest possible notation, with a single index identifying each individual and therefore standing at once for its genotype and environment. The estimation models presented below are then adapted to the structure of the data, by declaring genetic effects that describe the relationships between individuals, whether genotypes are repeated identically (pure lines, F1 hybrids or clones) or share a family structure of any kind (see Simulation design and Phenotype construction). For a single trait, the phenotype of an individual *i* interacting with its *J*_*i*_ neighbours of the same species can then be written as:

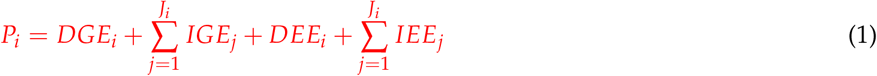

where *P*_*i*_ is the phenotypic observation measured on individual *i* and *J*_*i*_ its number of neighbors. The *DGE*_*i*_ corresponds to the genetic potential of individual *i* to express a given trait, independent of its social environment, and is conceptually equivalent to the classical genetic value in quantitative genetics. The *IGE*_*j*_, referred to as the indirect breeding value or social genetic effect by (Bijma *et al*. 2007b), represents the heritable contribution of a neighbor’s genotype to the focal individual’s phenotype, through any trait that mediates the interaction, such as resource uptake or competition. On the environmental side, *DEE*_*i*_ captures all non-genetic factors that directly affect individual *i*, such as local soil heterogeneity and microclimatic conditions. Finally, *IEE*_*j*_, termed *E*_*social*_ by (Bijma *et al*. 2007b), is the non-genetic contribution of the neighbor *j* to the phenotype of individual *i*. (Bijma *et al*. 2007b) describe the social effect as an environmental effect provided by associates to the focal individual, its heritable part being the *IGE* and its non-heritable part the *IEE*; both are indexed by the neighbor that exerts them, and the model places no restriction on how they arise. In plants, a neighbor acts on the focal individual mainly through its own size and architecture, and it is therefore reasonable to read the *IEE* as reaching *i* through the phenotype of *j*: what a neighbor does to the focal plant depends on how tall, how leafy or how deeply rooted it is, and the *IEE* then carries the part of that phenotype which the genotype of *j* does not explain, for instance when a neighbor is larger for non-genetic reasons and shades *i* more than its genotype alone would predict. Every term is indexed by the individual, which keeps the notation light, but the genetic terms are properties of the genotype: *DGE*_*i*_ is the direct genetic effect of the genotype carried by individual *i*, and all the plants carrying that genotype share its value, while *DEE*_*i*_ is what remains once the direct genetic effects have been accounted for. If the replication of the genotypic identity is not complete between individuals (i.e not clones), the structure between them can be accounted by the kinship or by estimating “family-level” genotypic effects using known pedigree information. The same convention is used for indirect genetic effects in animal breeding (Bijma *et al*. 2007b), where each individual carries its own genotype and the distinction does not arise. In the statistical model the mapping is explicit, the assignment matrix *Zg* linking each plant to its genotype (Equation 11). All genetic effects can co-vary with one another, and their joint distribution is described by a variance-covariance matrix; the same applies to the environmental effects. More generally, the contribution of each neighbor can be weighted by an interaction factor *f*_*ij*_ that decreases with the distance between *i* and *j*, so that the indirect terms become ∑_*j*_ *f*_*ij*_ *IGE*_*j*_ and ∑_*j*_ *f*_*ij*_ *IEE*_*j*_, the neighborhood being extended beyond first-rank neighbors and rescaled to account for a dilution effect (Bijma 2010b); this general structure is detailed in Supplementary Section (General neighborhood structure). For clarity, we present, in the equations and in the simulation, the simplest case: a first-rank neighborhood with distance-independent interaction strength and no dilution.

The variance-based framework has been applied to a wide range of taxa and traits. In animal breeding, it has been used to reduce intraspecific competition without losing productivity gains (Muir 2005; Ellen *et al*. 2014, 2008; Bergsma *et al*. 2008; Camerlink *et al*. 2015; Santostefano *et al*. 2025). In plants, DGE and IGE variances and DGE-IGE covariances have been estimated mainly in tree species (Cappa and Cantet 2008; Costa E Silva and Kerr 2013; Costa e Silva *et al*. 2017; Brotherstone *et al*. 2011), and in annual species such as *Arabidopsis thaliana*, where they enabled advances in QTL discovery and genomic prediction (Montazeaud *et al*. 2023; Sato *et al*. 2024). The magnitude of IGEs relative to DGEs varies considerably across systems and can substantially alter the predicted response to selection (Bijma 2014). These applications, however, are almost exclusively intraspecific. The main variance-based formulation that has been extended to interspecific interactions is the general mixing ability-specific mixing ability (GMA-SMA) approach (Wright 1985; Forst *et al*. 2019; Haug *et al*. 2021; Sampoux *et al*. 2020), which uses the aggregated performance of the mixture rather than individual-level data. Haug et al. (2023) applied it to a pea-barley intercrop and, because the two grains can be separated at harvest, were able to decompose the mixing abilities into direct and indirect genetic effects within the same plot-level experiment (Haug *et al*. 2023; Salomon *et al*. 2026). Such a decomposition therefore requires each component to be measured separately. When components cannot be told apart, only the aggregated performance of the mixture is available through the estimation of the sum of the direct and indirect genetic effects (Wuest and Niklaus 2018), and the aggregate itself requires the same trait to be measured in comparable units for all components. These limitations restrict the ability to identify the genetic architecture of interactions or to select genotypes with contrasting DGE and IGE profiles.

IGE studies in plants also remain few because their accurate estimation is experimentally demanding. The main bottleneck is the combinatorial explosion of genotype and species combinations. A full factorial combining 30 candidate genotypes of each of two species with three replicates per pair already requires 2,700 plots, and a third species brings this number to 81,000, beyond the reach of most breeding programs (Firmat and Litrico 2022). Incomplete factorial designs reduce this cost by testing each genotype with only a subset of the possible partners (Forst *et al*. 2019; Haug *et al*. 2021), and the choice of design and of the number of replicates per genotype has been studied in this context (Forst *et al*. 2019; Forst 2018; Haug *et al*. 2021; Freville *et al*. 2022; Salomon *et al*. 2026). The experimental unit of these designs remains, almost exclusively, a plot holding one pair of genotypes, from the same species or from two species. Addressing intraspecific and interspecific interactions at the same time then requires a separate trial for each interaction type.

In this study we propose an alternative framework in which two species are grown side by side in a single trial, and every plant is measured individually. Each plant has neighbors of both species, so it carries information on both types of interaction at once. The model then estimates in one step what a genotype does for itself, what it does to its conspecific neighbors and what it does to its companion-species neighbors, together with the covariances between all effects. Leveraging these covariances increases the statistical power and the accuracy of the estimation of all genetic values (Henderson and Quaas 1976). It also gives breeders access to the correlated responses that selection generates: because these effects covary, selecting individuals on their DGE alone shifts the mean intraspecific and interspecific IGE of the next generation. Estimating the full covariance structure among DGE, intraspecific IGE and interspecific IGE is therefore necessary to predict, and exploit, these correlated responses when designing selection indices (Hazel 1943; Bourke *et al*. 2021; PESEK and BAKER 1970).

This study aims to (i) propose an individual-level, variance-based framework that jointly models the direct and indirect genetic and environmental effects in plant mixtures, both intra- and interspecific, together with their full covariance structure, (ii) evaluate on simulated data under which conditions of heritability and replication these effects can be accurately estimated, (iii) compare the accuracy reached by a continuous neighborhood design versus a pairwise design at an equal number of plants, and (iv) apply the framework to a semi-controlled experiment in which 181 durum wheat lines and 106 alfalfa half-sib families, representing 3840 plants, were grown together and individually phenotyped.

## Materials and Methods

### Background theory and extension

The notation used throughout is collected in Table 1. Superscripts denote the species in which an effect is expressed, and subscripts the species that carries it, so that 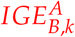 is the indirect genetic effect of an individual *k* of species *B* on the phenotype of a neighbor of species *A*.

**Table 1.**
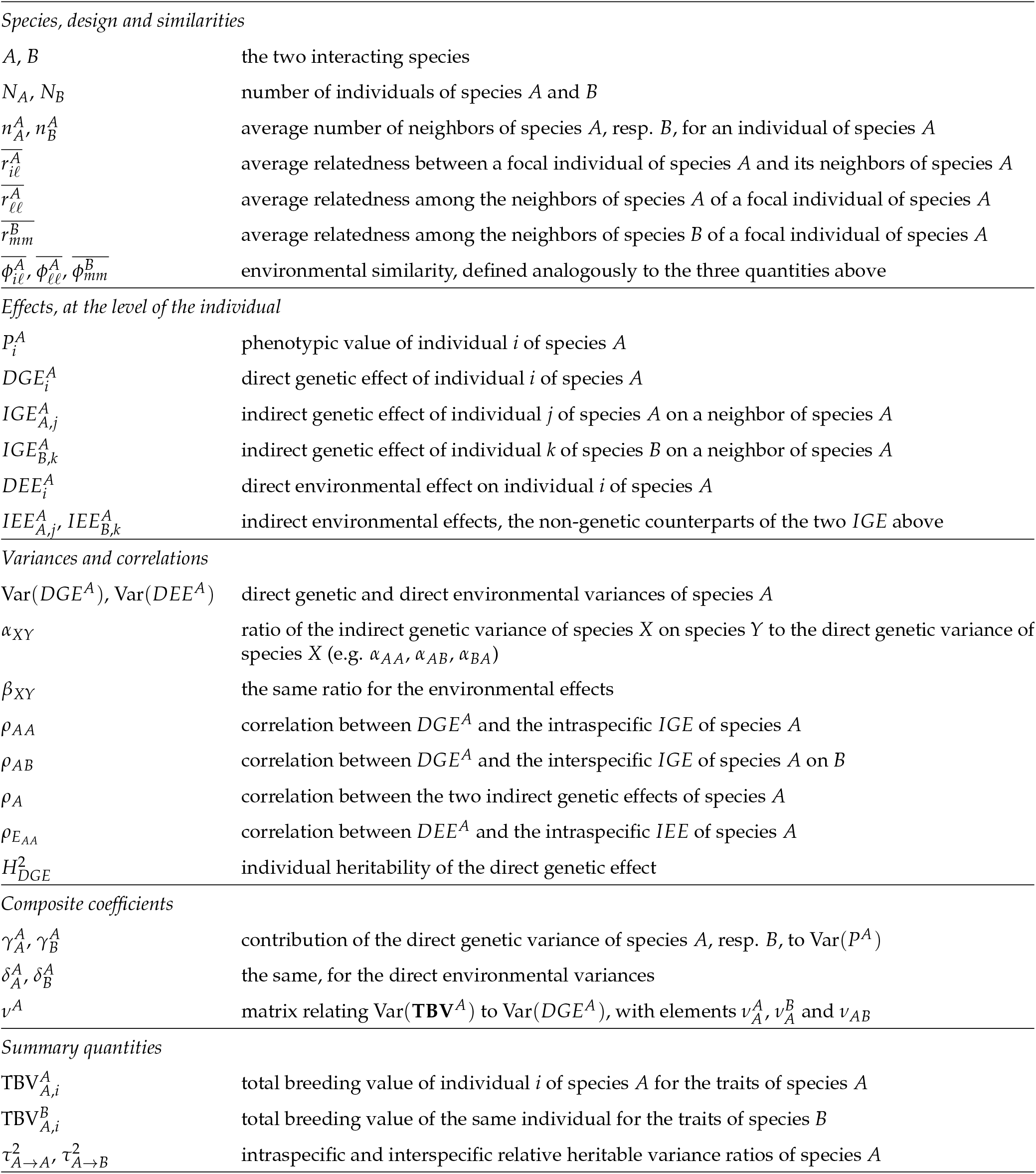
Notation used in the theoretical framework. Equations are given for species *A*; those for species *B* follow by exchanging *A* and *B*. The incidence matrices and the random-effect vectors of the statistical model are defined in Section Statistical model.

#### Phenotypic value

We build on the variance-based framework established by (Bijma *et al*. 2007b,a) for direct and social genetic effects in a single species (Eq. 1), and further developed by (Cappa and Cantet 2008; Costa E Silva and Kerr 2013; Silva *et al*. 2013) for complex neighborhoods with spatially structured interactions, varying distances between plants and varying numbers of neighbors. We extend it to two species, *A* and *B*, with *N*_*A*_ and *N*_*B*_ individuals respectively, so that the phenotypic value of an individual *i* of species *A* depends on its own genetic and environmental effects and on those of its neighbors of both species:

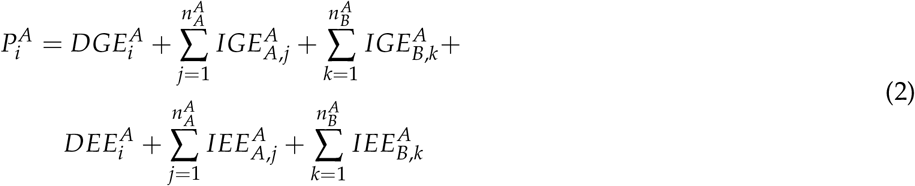

For clarity, we assume that all individuals have the same number of neighbors and that the strength of interaction does not depend on the distance between plants. The interaction factors *f*_*ij*_ can then be omitted: every individual receives the same number of unit-weighted contributions, the indirect terms are on the same scale for all plants, and the global variances can be estimated without confounding. The general case is treated in Supplementary Section General neighborhood structure.

#### Phenotypic variance

Using this model, we can decompose the phenotypic variance of species *A* if we make the following assumptions:

- All individual plants within each species share a common population variance, and each individual has a constant average number of neighbors of the same species as well as a constant average number of neighbors of the other species.
- Genetic relatedness between individuals is modeled by coefficients 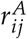, defined as twice the coefficient of coancestry (Lynch *et al*. 1998), and environmental similarity by coefficients 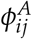.
- All covariances between genetic and environmental effects are zero.

The phenotypic variance of an individual *i* of species *A* then decomposes into the direct genetic and direct environmental variances of both species (derivation in Supplementary Equations S1a, S1b, S1c):

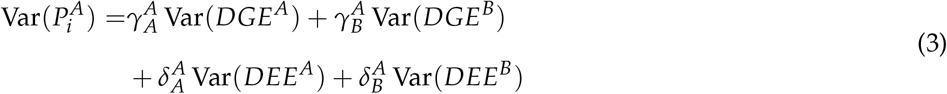

with:

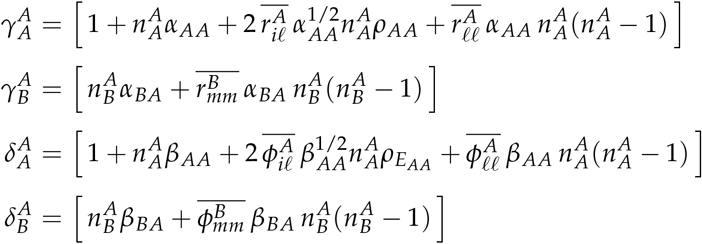

Equation 3 gives the population-level phenotypic variance. In a finite experiment the empirical variance also depends on the covariances between the phenotypes actually observed; the corresponding expected empirical variance is derived in Supplementary Equation S2.

### Total breeding value (TBV)

The total breeding value (TBV) was first defined by (Bijma *et al*. 2007b) as the sum of the heritable effects that an individual transmits to the next generation. In a monospecific design, with a constant average number of neighbors *n*, it is written

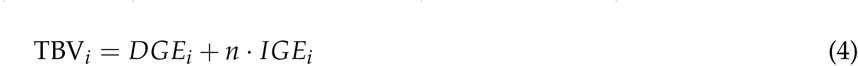

With two species, the traits are measured on different scales, so an interspecific IGE cannot be added to a DGE or to an intraspecific IGE. Following (Bijma *et al*. 2007a), who define the TBV as the total genetic contribution of an individual to the phenotypic variation of the population, we therefore split it into two components: 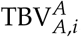, the contribution of an individual *i* of species *A* to the traits of its own species, which is the equivalent of (Bijma *et al*. 2007a), and 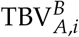, its contribution to the traits of species *B*.

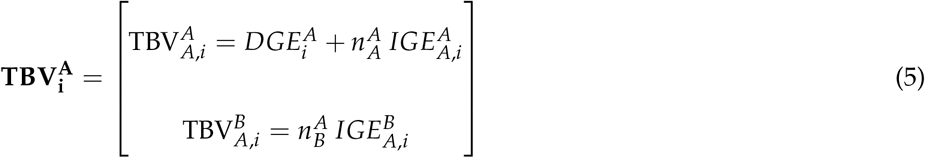

#### Total breeding value variance

The variance of the TBV measures the genetic variation available for selection. The two components can also covary, which constrains the progress achievable on both species at once.

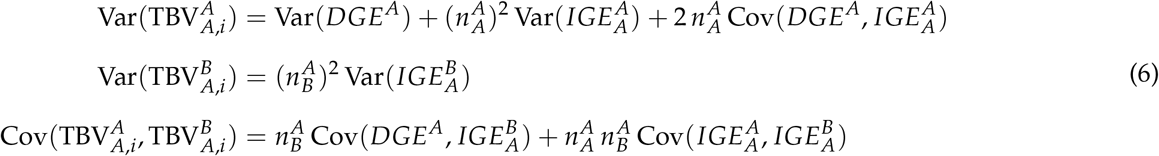

These three quantities are the entries of the variance-covariance matrix of 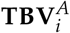. Under the same assumptions as for the phenotypic variance, this matrix can be expressed as a function of the direct genetic variance of the species:

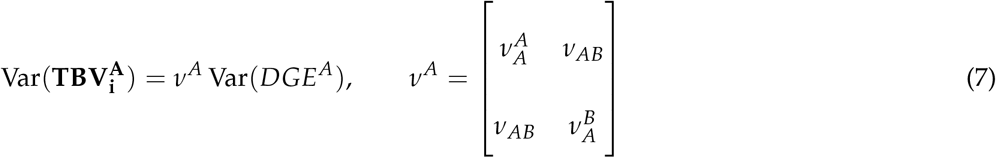

with:

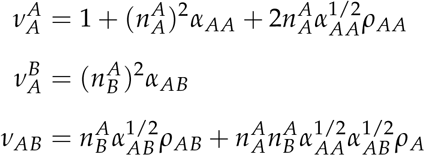

As for the phenotypic variance, the expected empirical value of the TBV variance is derived in Supplementary Section Empirical estimation of total breeding value variance.

#### Relative heritable variance

*τ*^2^ By analogy with classical heritability, we define the relative heritable variance ratio of a species as the ratio between the variance of its TBV and the phenotypic variance. In the single-species case, this ratio is the *τ*^2^ factor (Bergsma *et al*. 2008; Ellen *et al*. 2014; Bijma 2010a). We extend it to two species by defining an intraspecific and an interspecific ratio:

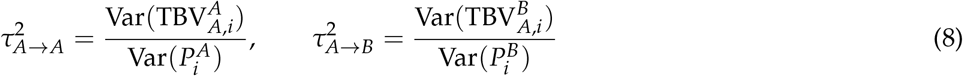

We refer to 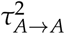 as the intraspecific *τ*^2^ and to 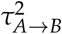 as the interspecific *τ*^2^, and likewise for species *B*. The intraspecific *τ*^2^ measures the share of the phenotypic variation of species *A* that is caused by the genetic effects of species *A* individuals, through their direct effects on their own phenotype and their indirect effects on conspecific neighbors; it is the equivalent of (Bergsma *et al*. 2008), the phenotypic variance now also containing the interspecific IGE received from species *B*. The interspecific *τ*^2^ measures the share of the phenotypic variation of species *B* that is caused by the indirect genetic effects of species *A*.

The two intraspecific ratios are linked by construction. Substituting Equation 8 into Supplementary Equations S17 and S12 yields:

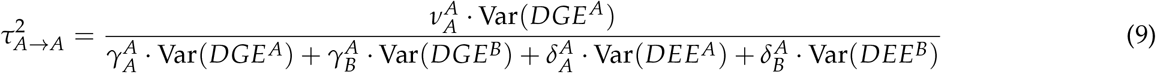

and symmetrically for 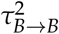. Both ratios include variance components from the two species, so the *τ*^2^ of one species cannot be estimated independently of the other.

### Two species simulation study

To assess the accuracy of our equations and our ability to accurately estimate the different genetic effects, we performed simulations of two interacting species. The simulations were run using the R programming language (R Core Team 2025) and the asreml-R package (The VSNi Team 2023) to fit the linear mixed models.

#### Simulation design

We simulated an alternating-row layout, as classically used in intercropping systems such as cereal– legume mixtures (Figure 1) (Ofori and Stern 1987). Each row holds a single species, inter-row and within-row spacings are constant and equal, and only first-rank neighbors interact, with a strength independent of the distance between plants. The field is large enough for the numbers of same-species and other-species neighbors to be constant, so that edge effects can be neglected.

**Figure 1.**
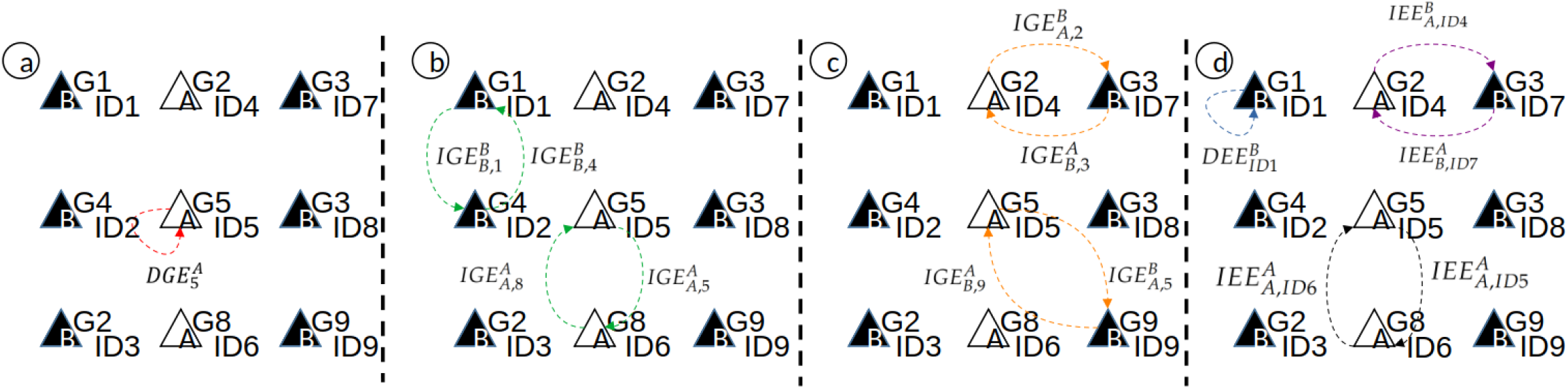
Schematic view of the experimental design used in the simulation. Every triangle represents one individual plant from species A or B. The genotypic identity of the plant is represented in the top right corner of each plant, and the unique identifier of the plant is indicated in the bottom right corner. Panel a shows an example of a direct genetic effect from a plant of genotype 5 (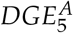; dashed line with an arrow). Panel b illustrates the two types of intraspecific indirect genetic interactions, with 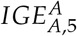 indicating the effect of genotype 5 of species A on a plant of species A and 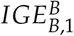 indicating the effect of genotype 1 of species B on its neighbors of species B. Panel c illustrates the two types of interspecific indirect genetic interactions, either within-row or along the diagonal, with 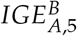 indicating the effect of genotype 5 of species A on its neighbors of species B and 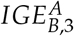 indicating the effect of genotype 3 of species B on its neighbors of species A. Panel d illustrates the environmental effects. In contrast with the genetic effects, which depend on the genotype, the environmental effects are attached to the individual identifier of the plant, reflecting their non-heritable, plant-specific nature. It shows the direct environmental effect of a plant on its own phenotype 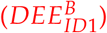, the two intraspecific indirect environmental effects, with 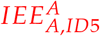 indicating the effect of individual ID5 of species A on a plant of species A, and the two interspecific indirect environmental effects, with 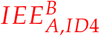 indicating the effect of individual ID4 of species A on its neighbors of species B.

In this design, each individual plant has up to eight neighbors: two from the same species and six from the other species. Within each species, genotypes were assigned to positions at random: each genotype was replicated a balanced number of times and then placed randomly across the available positions of the field. This ensures that direct and indirect effects remain separable: a systematic association between a plant’s own genotype and those of its neighbors would correlate the corresponding incidence matrices and confound the two effects.

Each individual belongs to one of two species, A or B, and to one of the *n*_*genotype*_ genotypes within each species. Each genotype was replicated *n*_*rep*_ times within each species. Therefore, the total number of plants per species was *N* = *n*_*genotype*_ *× n*_*rep*_.

Each plant carries two identifiers, its genotype and its own plant identifier, both shown in Figure 1. Genetic effects are attached to the genotype and environmental effects to the plant identifier, which is the only difference between the two families of incidence matrices. Taking the three plants of species *A* in the central column of Figure 1, the neighborhood matrix 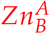 counts, for each of them, the number of neighbors carrying each genotype of species *B*:

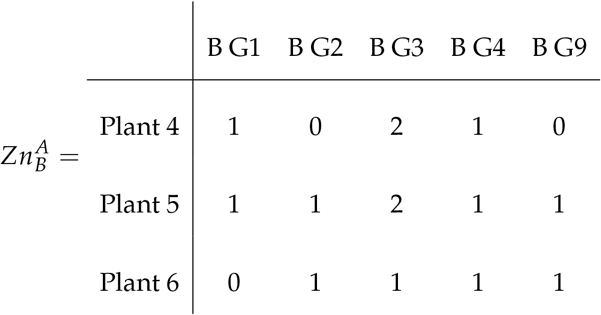

Genotype G3 is carried by two plants, ID7 and ID8, which is why plant ID4 scores 2 in that column. The corresponding environmental matrix 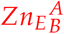 describes the same neighborhood between plant identifiers, so that each neighbor has its own column and all entries are 0 or 1:

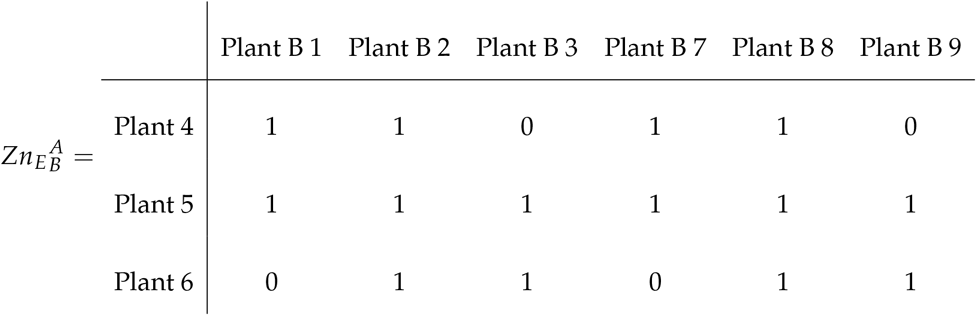

The remaining matrices follow the same construction and are given in Supplementary Section Remaining incidence matrices of the simulated design: 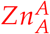 and 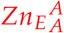 for the conspecific neighbors, the assignment matrix *Zg*^*A*^ linking each plant of species *A* to its genotype, and 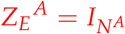, which links each plant to its own direct environmental effect.

#### Genome generation

For each species, genotypes were generated with the coalescent algorithm implemented in the R package *scrm* (Paul R. Staab *et al*. 2015), with 14 diploid chromosomes of 5 *×* 10^5^ base pairs, an effective population size of 10^4^, mutation and recombination rates of 10^−8^ per base pair per generation, a single founder population and at most 10^6^ segregating sites. These values are standard for eukaryotes and produce a non-structured genome representative of a panmictic eukaryotic population, without mimicking any particular species; the chromosome length was kept short to reduce the computational burden.

From the simulated sequences we extracted single nucleotide polymorphisms with a minor allele frequency of 0.20, and computed the VanRaden genomic relationship matrix (VanRaden 2008) for each species with the *kinship* function of the R package *statgenGWAS* (Rossum and Kruijer 2025).

#### Effect construction

For the genetic effects of a species, the variance-covariance matrix **G** between the direct effect, the intraspecific IGE and the interspecific IGE is

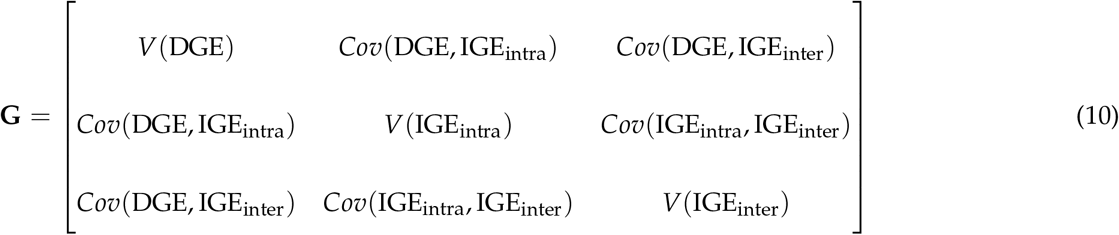

where 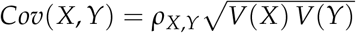, the correlations being those of Table 2. The environmental variance-covariance matrix **E** has the same structure, with the DEE and the two IEE in place of the three genetic effects.

**Table 2.** Parameters of the simulation study. All combinations of the values listed were explored, with the same values for both species.

| Parameter | Values | Rationale |
| --- | --- | --- |
| *Design and replication* |  |  |
| $n_{\text{genotype}}$ | 100 | Fixed. The genetic effects are drawn from a target variance-covariance structure, and enough genotypes are needed for the sample actually drawn to represent it faithfully. |
| $n_{\text{rep}}$ | 2, 5, 10, 15, 20 | Determines the size of the experiment, from 400 to 4,000 plants in total, and therefore the statistical power. |
| $H^2_{DGE}$ | 0.05, 0.1, 0.2, 0.3, 0.5, 0.7 | Leading parameter of the study (Eq. 12). Spans the range from a barely heritable trait to a highly heritable one. |
| $\text{Var}(DGE)$ | 1 | Fixed as the unit of scale. $\text{Var}(DEE)$ is then set so as to reach the target heritability (Supplementary Eq. S18). |
| *Variance ratios* |  |  |
| $V(IGE_{\text{intra}})/V(DGE)$ | 0.05, 0.1 | The few published estimates for growth- and reproduction-related traits in plants and trees are of this order of magnitude (Montazeaud *et al.* 2023; Silva *et al.* 2013; Cappa and Cantet 2008). Small ratios are also the hardest case, since the indirect variance is then the most difficult to detect and to separate from the other effects; larger ratios would only make estimation easier. |
| $V(IGE_{\text{inter}})/V(DGE)$ | 0.05, 0.1 | (Haug *et al.* 2023) report interspecific effects of about the same magnitude as the intraspecific ones, so the same range was used. |
| $V(IEE_{\text{intra}})/V(DEE)$ | 0, 0.1 | No published estimate is available for the indirect environmental effects, previous studies having absorbed them into a spatial or residual structure instead of explicitly modelling them (Costa e Silva *et al.* 2017; Silva *et al.* 2013). The IGE range was reused, with 0 to cover the case where neighbors carry no environmental effect. |
| $V(IEE_{\text{inter}})/V(DEE)$ | 0, 0.1 | As above. |
| *Correlations* |  |  |
| $\rho_{DGE, IGE_{\text{intra}}}$ | -0.6, 0 | Published estimates are null to slightly negative (Montazeaud *et al.* 2023; Cappa and Cantet 2008; Costa E Silva and Kerr 2013). |
| $\rho_{DGE, IGE_{\text{inter}}}$ | -0.6, 0 | No published estimate. The intraspecific range was reused. |
| $\rho_{IGE_{\text{intra}}, IGE_{\text{inter}}}$ | 0, 0.6 | No published estimate. Null to moderately positive, on the hypothesis that competitive ability rests partly on the same genes in both interaction types. |
| $\rho_{DEE, IEE_{\text{intra}}}$ | -0.3 | Fixed. Represents a moderate non-heritable competition between neighbors. |
| $\rho_{DEE, IEE_{\text{inter}}}$ | -0.3 | Fixed, as above. |
| $\rho_{IEE_{\text{intra}}, IEE_{\text{inter}}}$ | 0.3 | Fixed. A plant that strongly affects one species tends to affect the other in the same way. |
| *Simulation* |  |  |
| Replicate simulations | 50 per parameter set | Measures the variability between runs of the same parameter set, which comes from the initial population, the spatial arrangement of the plants and model convergence. A small share of these fits does not return a complete genetic covariance matrix, as detailed in Section Model fitting and Estimation accuracies. |

Additive allelic effects were drawn for each SNP from **G** with the *mvrnorm* function of the R package *MASS* (Venables and Ripley 2002), and the three genetic effects of a genotype were obtained by summing the effects of all its SNPs. Environmental effect values were drawn from **E** directly, once per plant. The genetic effects are therefore shared by all the plants of the same genotype, whereas the environmental effects are attached to a single plant, each plant receiving its own DEE and its own pair of IEE.

#### Phenotype construction

Combining these effects with the incidence matrices, we write the phenotype of the plants of species *A* as

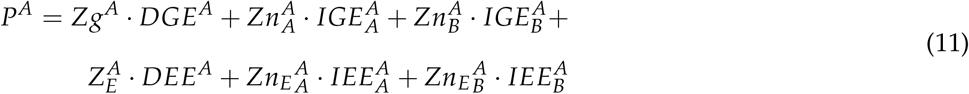

which is the model of Section Phenotypic value applied to the simulated design.

#### Simulation parameters

We selected the individual heritability of the DGE, defined as the ratio between the DGE variance and the phenotypic variance of the individuals, assuming no IGE variance, as the leading “variance” parameter.

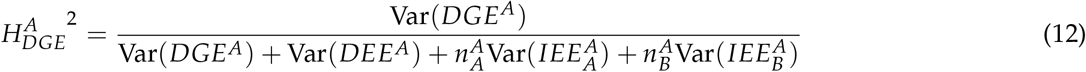

This choice was motivated by the fact that DGE individual heritability closely parallels the individual heritability in classical animal models, making it a familiar and useful indicator for breeders and geneticists.

The DGE variance was fixed to 1, and the DEE variance was then computed so as to reach the target heritability, under the assumption that there were no environmental similarities between plants and, as in the definition of 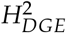, no indirect genetic variance. The expression, obtained by rearranging Equation 12, is given in Supplementary Equation S18.

We explored all combinations of the parameters given in Table 2, which also presents their range and their justification. The same values were used for both species.

To limit edge effects the field was made as square as possible, so the realized number of replicates per genotype could differ slightly from the target. For some parameter combinations the variance-covariance matrices were not positive definite; minor adjustments were then applied with the *make*.*positive*.*definite* function of the R package *corpcor* (Schafer *et al*. 2021).

#### Statistical model

To estimate the different effects underlying the phenotypes, we used linear mixed modeling and the restricted maximum likelihood (REML) approach with the *asreml* function from the R package *asreml-R* (The VSNi Team 2023). As genetic effects from both species are involved in the phenotypes of each species, we fitted a joint model for both species with two unstructured variance-covariance structures that account for the genetics of both species. This model takes the form of a “long-format” (stacked phenotype vectors) bivariate model:

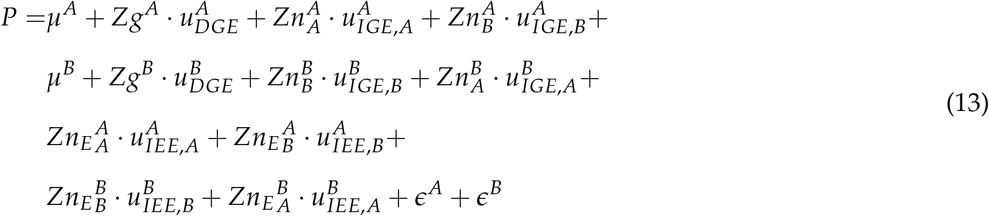

where *P* stacks the phenotypes of both species, *µ*^*A*^ and *µ*^*B*^ are the species’ intercepts, the *u* vectors are the random effects labeled as in Table 1, and *ϵ*^*A*^ and *ϵ*^*B*^ are the residuals, which carry the direct environmental effects.

The genetic effects were given an unstructured variance-covariance matrix per species. Each structure groups the three effects carried by one species, that is, its direct effect and its two indirect effects, rather than the effects it receives:

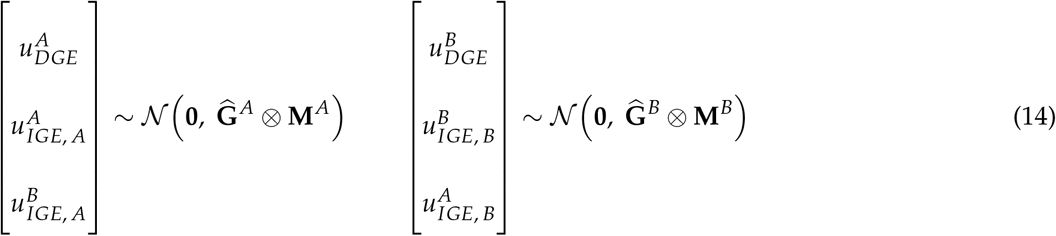

where 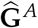 and 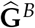 are the estimated variance-covariance matrices between the genetic effects of each species, **M** and **M**^*B*^ the covariance structures among the genotypes of each species, and ⊗ the Kronecker product. Each **M** is either the kinship matrix of the species concerned, which targets the additive genetic variance, or the identity matrix of the same order, which targets the total genetic variance among the genotypes evaluated: **M**^*A*^ ∈ *{***K**^*A*^, **I**^*A*^*}* and **M**^*B*^ ∈ *{***K**^*B*^, **I**^*B*^*}*.

The two structures answer different questions. With the kinship matrix, genotypes are related and each one is informed by the records of its relatives as well as its own, so the variance estimated is the additive genetic variance of the population they are drawn from. With the identity, genotypes are treated as unrelated and each is informed only by its own records, so the variance estimated is the total genetic variance among the genotypes evaluated, additive and non-additive alike.

As the direct environmental effect is included in the residual variance, only the variance-covariance matrices of the indirect environmental effects can be estimated:

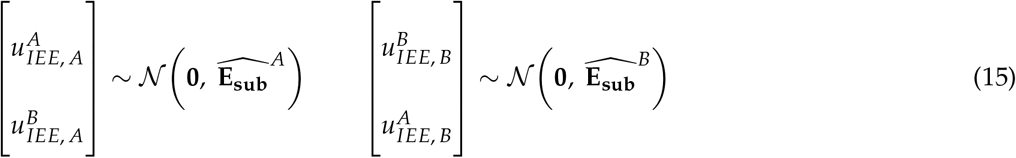

where 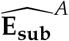 and 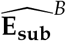 are the sub-matrices of the environmental variance-covariance structure **E** that exclude the DEE terms. As a consequence, although the DEE-IEE correlation was simulated (at −0.3), it is not estimated by the model and is effectively forced to zero during the inference: the DEE is defined at the plant level with a single observation per plant, so it is confounded with the residual, and estimating its covariance with the IEE would require a covariance between the residual and a random effect, which is not identifiable here.

The residuals of the two species were assumed independent, with species-specific variances 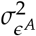 and 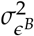.

To assess the importance of accounting for IEEs in the inference model, we also fitted a so-called “null” model with the same general structure but without the IEEs.

#### Comparison with a pairwise design

(Firmat and Litrico 2022) showed that full factorial designs, in which every genotype of one species is combined with every genotype of the other, require a number of units that is out of reach of a breeding program. Incomplete factorial designs reduce this cost by testing each genotype with only a subset of the possible partners (Forst *et al*. 2019; Haug *et al*. 2021; Salomon *et al*. 2026). Their experimental unit (i.e. pot or plot) accommodates a limited number of genotypes (usually two), severely restricting the number of combinations and hence interactions that can be tested.

We compared the continuous neighborhood design used throughout this work with the smallest and most explicit form of a pairwise design, a pot holding two plants (Figure 2). Both designs were simulated from the same phenotypic model, and the genetic effects were estimated at the individual plant level with Equation 2 in both, so that the comparison reflects the design and not the model used to analyze it. A pairwise unit holds two genotypes and therefore a single interaction type, so a pairwise design consists of three separate trials, *A* with *A, B* with *B* and *A* with *B*, analyzed together. The continuous neighborhood provides both interaction types on the same spatial footprint.

**Figure 2.**
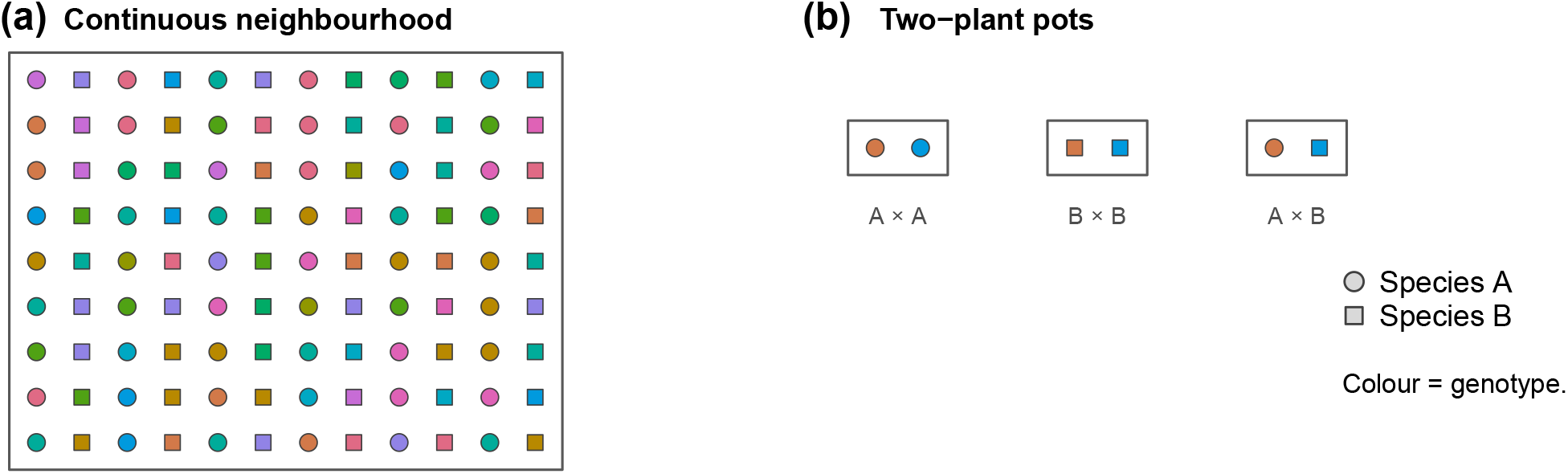
Comparison of two experimental designs for plant neighborhood experiments. Shape denotes species and color denotes genotype. **(a)** The continuous neighborhood used throughout this work, in which every plant may be a different genotype, so that many genotypes are grown per unit area and each plant interacts with several partners of both species at once. **(b)** Two-plant pots, in which one genotype interacts with a single partner; a pot involves a single interaction type, so the three interaction types require three separate pots.

Each design was simulated twice, with or without indirect environmental effects. Where they were simulated, they were fitted in the continuous neighborhood and left out of the pairwise design, because they are not estimable there. In a pot, each plant “emits” its indirect environmental effect towards exactly one receiver and receives the same from exactly one emitter, so that their incidence matrix is a permutation and their contribution to the covariance of the observations is *V*(*IEE*) **I**, which has the same form as the residual: the two cannot be disentangled, regardless of the number of pots. In a continuous neighborhood design a plant receives from eight emitters and each plant emits towards eight receivers, so the same effect appears in several observations and can be separated from the residual. The genetic effects were estimated in the presence of indirect environmental effects in both designs, so that the comparison demonstrates what omitting them costs the pairwise design.

The experimental effort was counted as the number of plants grown, which is also the number of phenotypic measurements taken, every plant being measured in both designs. Each design was simulated at several sizes, from 5 to 500 replicates per genotype for the continuous neighborhood and from 2 to 250 pots per genotype in the pairwise set-up. Each parameter combination was replicated 40 times. The comparison was run at two values of the dilution factor. At *d* = 0 each neighbor contributes its full indirect effect, so a plant of the continuous grid receives the sum of eight contributions where a potted plant receives one, and the designs differ both in the number of distinct partners a genotype grow in the vicinity of and in the amount of neighborhood signal each plant receives. At *d* = 1/2 the weights are rescaled so that every plant is exposed to the same total indirect variance whatever its number of neighbors (Cappa and Cantet 2008; Costa E Silva and Kerr 2013), leaving only the number of distinct partners to explain the differences in accuracy between the pot and continuous designs (Supplementary Section Design comparison at normalized exposure).

### Model fitting and Estimation accuracies

All the fits reported here used **M** = **I**, the genotypes being simulated as an unstructured sample in which the kinship matrix departs little from the identity. Fitting the same simulated datasets with the kinship matrix was consistently a little better than fitting them with the identity, but the difference was small (Supplementary Section Genotype covariance structure: kinship or identity). Working without it therefore costs little here, and gives a lower bound on what the same experiment would achieve.

We used two indices to quantify how accurately the model recovered the simulated genetic parameters: i) the relative error index (*V*_*error*_), calculated as the 10-logarithm of the ratio between the estimated and true variances for each effect, and ii) the Pearson correlation between the true simulated genotypic values and their estimated values (BLUPs). Both are computed against the simulated truth, and not against the data the models were fitted to, so a model gains nothing here from having more parameters. It is important to note that those correlations can be upper-bounded in a finite experiment. Their upper bounds are set by the size of the effect relative to the remaining variation and by the number of phenotypes in which the effect appears. That bound is derived for each genetic effect in Supplementary Section Upper bound on the accuracy of the estimated genetic effects.

A small share of the fits did not converge, or returned an incomplete genetic covariance matrix: 98.7 % of the fits were complete overall. These runs are not shown in the figures. They concentrate at the highest heritabilities and replication levels, where the genetic effects become difficult to tell apart from one another (Section What the framework estimates, and at what experimental cost).

The analytical expressions were also checked against the simulated data. For each of 128 parameter combinations, theoretical predictions were obtained with the analytical formulas (Supplementary Equations S2, S13 and S14, and Equation 8), and compared with the values computed from the simulated phenotypes and per-individual TBV using the *var* function of R (R Core Team 2025). The grid deliberately included scenarios in which *τ*^2^ exceeds 1, notably when Cov(*DGE*, 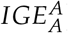) is strongly positive, a regime specific to systems with social interactions, to verify that the equations remain accurate beyond the classical [0, 1] interval of heritability-like ratios.

To measure the computational cost of the model, we fitted three models to the same simulated experiment, differing only in their structure: a conventional single-species model with a direct effect and an intraspecific IGE, giving 2*q* genetic equations for *q* genotypes per species; a two-species model without the interspecific term, giving 4*q* equations; and the full model of Equation 13, giving 6*q* equations. The covariance among the correlated effects was unstructured in every case. Each of the three models was fitted twice, once with the kinship matrix as the genotype covariance structure (**M** = **K**) and once with the identity (**M** = **I**), since the inverse of a kinship matrix is dense whereas the identity leaves the genetic block separable by genotype, and the two are therefore not expected to cost the same. We varied the number of genotypes with the number of plants held at 8,000, and separately the number of plants with the number of genotypes held at 200 per species. All fits were single-threaded, with three replicates per point and the median reported, and no indirect environmental effect was fitted, so that the genetic block was the only thing that varied.

### Application to a wheat-alfalfa semi-controlled experiment

The framework was then applied to a two-species semi-controlled experiment carried out at Montpellier, France, from December 2024 to June 2025. Durum wheat (*Triticum turgidum* ssp. *durum*), represented by 181 pure lines with a broad genetic diversity (David *et al*. 2014), and alfalfa (*Medicago sativa* ssp. sativa), represented by 106 half-sib families were co-cultivated outside together in twelve containers of one cubic meter filled with soil. Each alfalfa half-sib family was obtained from a single mother plant from the synthetic polycross population “Syn1” (El Ghazzal *et al*. 2025).

Within each container, plants of both species were sown in alternating columns in a grid of 20 columns by 16 rows with a 5 cm step, which gives 160 plants of each species per container and 1920 plants of each species in the experiment. Each wheat genotype was replicated 10.6 times on average and each alfalfa genotype (mother plant) 18.1 times. Every plant was positioned, labelled and phenotyped individually, so that the experiment closely corresponds to the the design presented in Figure 1. Although multiple traits were recorded, we report here the number of grains per plant for wheat and the top 6 cm dry biomass of the fourth cut for alfalfa just after the wheat harvest. Both traits were centered and scaled to unit variance within species.

The neighborhood matrices were built as described in Supplementary Section General neighborhood structure, each neighbor being weighted by 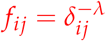 with *λ* = 1 and *δ*_*ij*_ the distance between the two plants, without row normalization, which corresponds to the dilution factor *d* = 0 used in the simulations. The rank of the neighborhood matrix kept for the final analysis was chosen by AIC comparison (Supplementary Section Neighborhood order in the semi controlled experiment).

Equation 13 was then fitted to the two traits jointly, with **M** = **I** for both species. Sowing date, seed weight and container were fitted as fixed effects. Border plants differ from the others in two ways that the model would otherwise attribute to the neighborhood. They have fewer neighbors (accounted for within the Zn matrices) and they stand at the container edge, where light interception, temperature and water availability differ from the middle of the container. To account for this, we added a fixed border effect to the model for these individuals. Variances were estimated using the same model as used for the simulations (The VSNi Team 2023), apart from the fixed effects listed above. We fitted the model twice, once with the indirect environmental effects and once without them, and kept the version with the lowest AIC, which was the one without (2313.6 against 2316.9).

Each component is reported with *z*, the ratio of its estimate to its standard error, the percentage given alongside being its share of the phenotypic variance. We read |*z*| *>* 2 as support for a non-zero component, which remains an approximation, a variance being bounded below by zero.

## Results

### Analytical results

The analytical expressions for Var(*P*^*A*^ ), Var(**TBV**^*A*^), 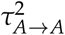 and 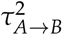 were validated against simulated data over 128 parameter combinations, including the regime where *τ*^2^ exceeds 1. They agreed closely with the values computed directly from the simulated phenotypes (Pearson *r >* 0.99 throughout), with no systematic bias across the heritability range explored (Supplementary Figures S2, S3 and S4). All results below were obtained with 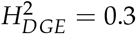 for both species and four neighbors of each species per plant, all *β* coefficients being zero; *α*_*inter*_ and *ρ*_*inter*_ denote the interspecific variance ratio and correlation, that is *α*_*AB*_ and *ρ*_*AB*_ for species *A*.

#### Relative heritable variance ratios

Figure 3a shows how 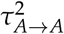 varies as a function of the magnitude of intraspecific IGEs, *α*_*AA*_, and of the correlation between DGE and intraspecific IGE, *ρ*_*AA*_. Three regions were identified. When *ρ*_*AA*_ is negative, 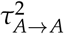 can fall below *H*^2^, even for moderate values of *α*_*AA*_. In this case, intraspecific social interactions constrain heritable variance below the baseline expected in the absence of social effects. When *ρ*_*AA*_ is positive and *α*_*AA*_ is moderate, 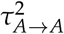 exceeds *H*^2^ but remains below 1, indicating that intraspecific social interactions amplify the heritable variance available for selection without exceeding the total phenotypic variance available for selection. Finally, when both *ρ*_*AA*_ and *α*_*AA*_ are large, 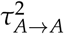 can exceed 1 and can reach values above 2. In this configuration, the total heritable variance exceeds the phenotypic variance of a population of isolated individuals, a result that is specific to systems with social interactions and reflects the cumulative genetic effects that each individual exerts on all its neighbors. The sign and magnitude of *ρ*_*AA*_ therefore play a critical role in determining whether intraspecific interactions constrain or amplify genetic progress.

**Figure 3.**
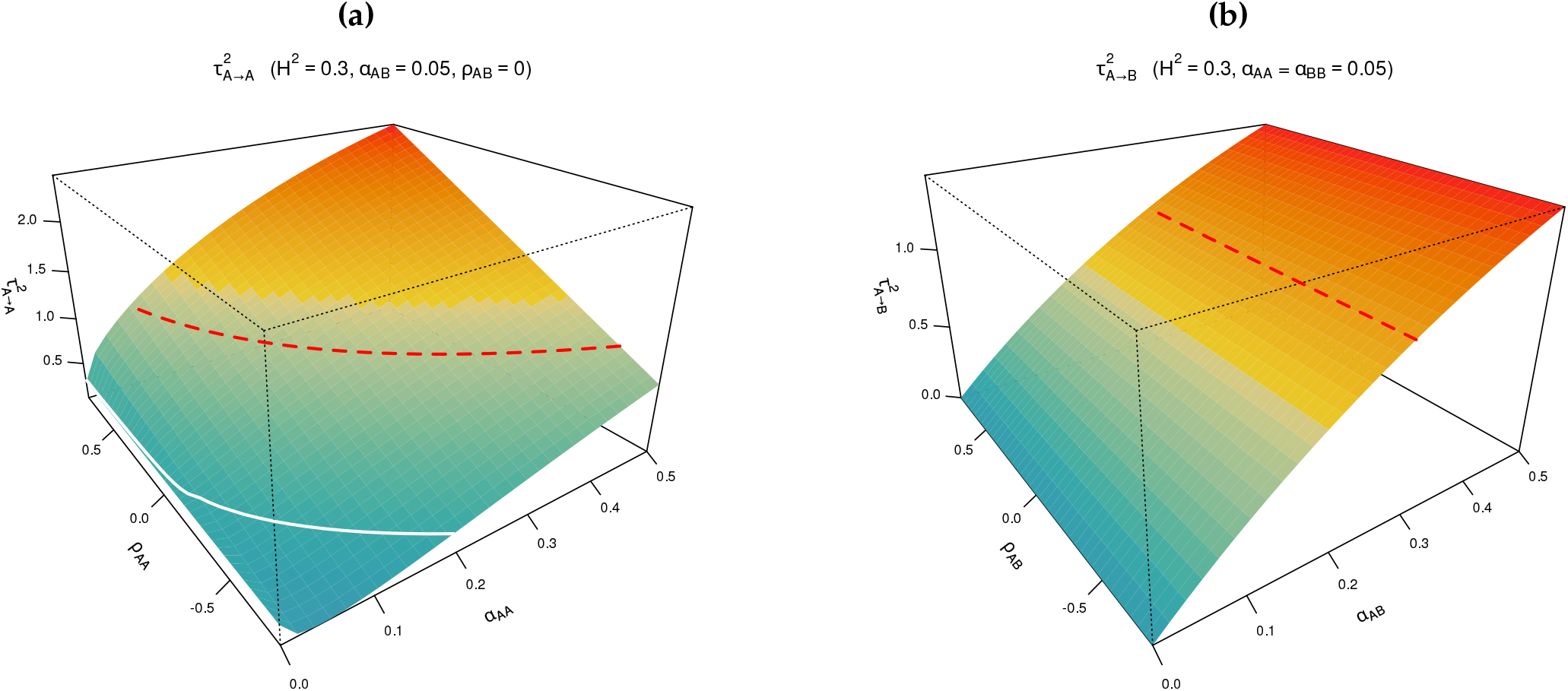
Relative heritable variance ratios of species *A*, with 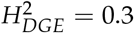 for both species. **(a)** Intraspecific ratio 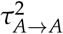, the proportion of phenotypic variance in species *A* that is heritable through both the direct genetic effects of each individual on its own phenotype and its indirect genetic effects on conspecific neighbors, as a function of *α*_*AA*_ and of *ρ*_*AA*_, with interspecific parameters fixed at small values (*α*_*inter*_ = 0.05, *ρ*_*inter*_ = 0) to isolate the intraspecific dynamics. The white line marks 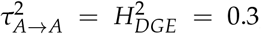, the heritable variance expected in the absence of social interactions. **(b)** Interspecific ratio 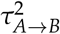, the proportion of phenotypic variance in species *B* attributable to the indirect genetic effects of species *A* individuals on their heterospecific neighbors, as a function of *α*_*AB*_ and of *ρ*_*AB*_, with intraspecific parameters fixed at *α*_*AA*_ = *α*_*BB*_ = 0.05. In both panels the red dashed line indicates a ratio of 1, above which the total heritable variance exceeds the phenotypic variance of a population of isolated individuals.

The interspecific ratio behaves differently (Figure 3b). Its surface is entirely determined by *α*_*AB*_: increasing the magnitude of interspecific IGEs leads to a near-linear increase in 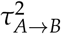, regardless of the value of *ρ*_*AB*_. Even moderate values of *α*_*AB*_, of the order of 0.2 to 0.3, are sufficient to make 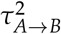 exceed 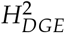, suggesting that selecting species *A* could substantially improve the mean phenotype of species *B* in the mixture, without any direct selection on species *B* itself, provided that the co-cultivation pattern is maintained. For large values of *α*_*AB*_, 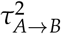 can exceed 1, the same cumulative effect as in the intraspecific case, each individual of species *A* affecting all its heterospecific neighbors at once. The independence of 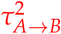 from *ρ*_*AB*_ has a simple explanation. The interspecific IGE of species *A* is expressed in the phenotype of species *B*, while the direct genetic effect of species *A* acts on its own phenotype, so the two never enter the same phenotypic variance. The correlation *ρ*_*AB*_ between them therefore appears neither in the phenotypic variance of species *B* nor in the variance of the interspecific total breeding value, and 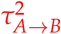 does not depend on it.

#### Covariance between 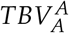 and 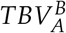

Figure 4 shows how 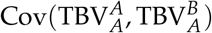 varies as a function of *α*_*AA*_ and *α*_*AB*_, with fixed correlation parameters (*ρ*_*AB*_ = −0.6 and *ρ*_*A*_ = 0.6). A positive covariance indicates that individuals genetically beneficial for species *A* also tend to benefit species *B*, allowing the simultaneous improvement of both species through selection. A negative covariance indicates a trade-off between the two situations. As shown in Equation 6, this covariance is decomposed into two terms with opposite signs. The first term, 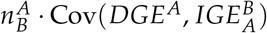, scales with *α*_*AB*_ and is driven by *ρ*_*AB*_ = −0.6; it is always negative in this scenario. The second term, 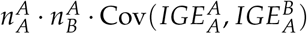, scales with both *α*_*AA*_ and *α*_*AB*_ and is driven by *ρ*_*A*_ = 0.6; it is positive. Therefore, the sign of the total covariance results from the competition between these two factors. When *α*_*AA*_ is small, the first term dominates and the covariance is negative. As *α*_*AA*_ increases, the second term grows and eventually dominates, causing the covariance to be positive. This sign reversal shows that the existence of a trade-off between improving a species for itself and improving the associated species is not a fixed property of the system; it depends on the relative magnitude of intraspecific and interspecific IGEs. Unlike 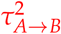, this covariance does depend on *ρ*_*AB*_: its first term is proportional to *ρ*_*AB*_, so the correlation between the direct and the interspecific indirect genetic effects of species *A* links the two total breeding values of species *A*, even though this correlation is absent from the phenotypic variance of either species. Supplementary Figure S5 shows this dependence, with the covariance varying linearly with *ρ*_*AB*_ and *ρ*_*A*_ over the positive-definite region and independently of heritability.

**Figure 4.**
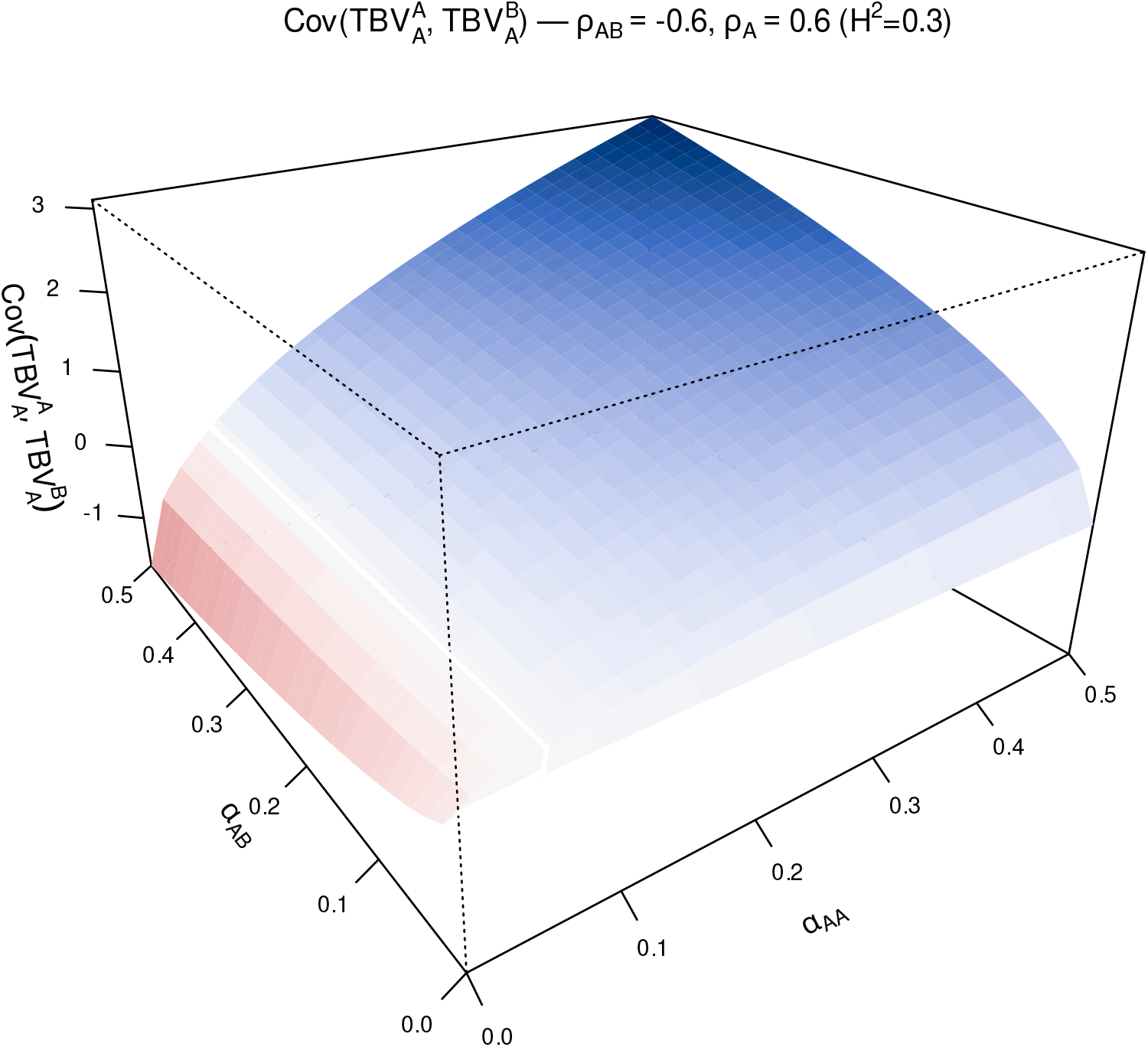
Covariance between the intraspecific total breeding value 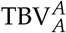 and the interspecific total breeding value 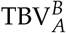 of species *A* , as a function of the proportion of intraspecific IGE variance compared to the DGE variance, *α*_*AA*_, and the proportion of interspecific IGE variance compared to the DGE variance, *α*_*AB*_. The other parameter values were 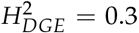, the correlation between DGE and interspecific IGE of *A* was *ρ*_*AB*_ = −0.6 while the correlation between interspecific IGE from *A* on *B* and intraspecific IGE within species *A* was *ρ*_*A*_ = 0.6. The graph was generated using equations 6. The white line marks the zero-covariance boundary, separating a region of negative covariance (in red) from a region of positive covariance (in blue).

### Estimation accuracies

In the following sections, all parameters were held constant except for the specific model parameters under investigation.

#### Impact of varying social genetic variance ratios

We first quantified the accuracy of genetic effect estimation across a range of heritabilities and DGE:IGE ratios, keeping the following parameters fixed for both species: [ineq.

In this first set of simulations, no indirect environmental effect was simulated (*V*(*IEE*) = 0): the environmental effects reduce to the direct term, which is contained in the residual. Estimation accuracies for DGE were already satisfactory for 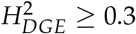 even with only 2 replicates per genotype (*n*_*rep*_ = 2), as expected for the accuracy of a genotype mean, which increases with both 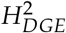 and *n*_*rep*_ (Figure 5; Supplementary Section Upper bound on the accuracy of the estimated genetic effects). With more replicates, the accuracy improved for all estimated genetic effects (DGE, intraspecific IGE, and interspecific IGE), even for low heritabilities, and *V*_*error*_ converged to zero. Notably, accuracy also increased with the variance ratio *V*(IGE)/*V*(DGE), although this effect was modest, and *V*_*error*_ converged only slightly faster at higher ratios. Intraspecific IGE accuracies were consistently lower than interspecific IGE accuracies, likely because within-species interactions involved fewer neighbors 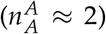 than between-species interactions 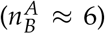 in our simulation; this difference disappeared at high heritability and replication.

**Figure 5.**
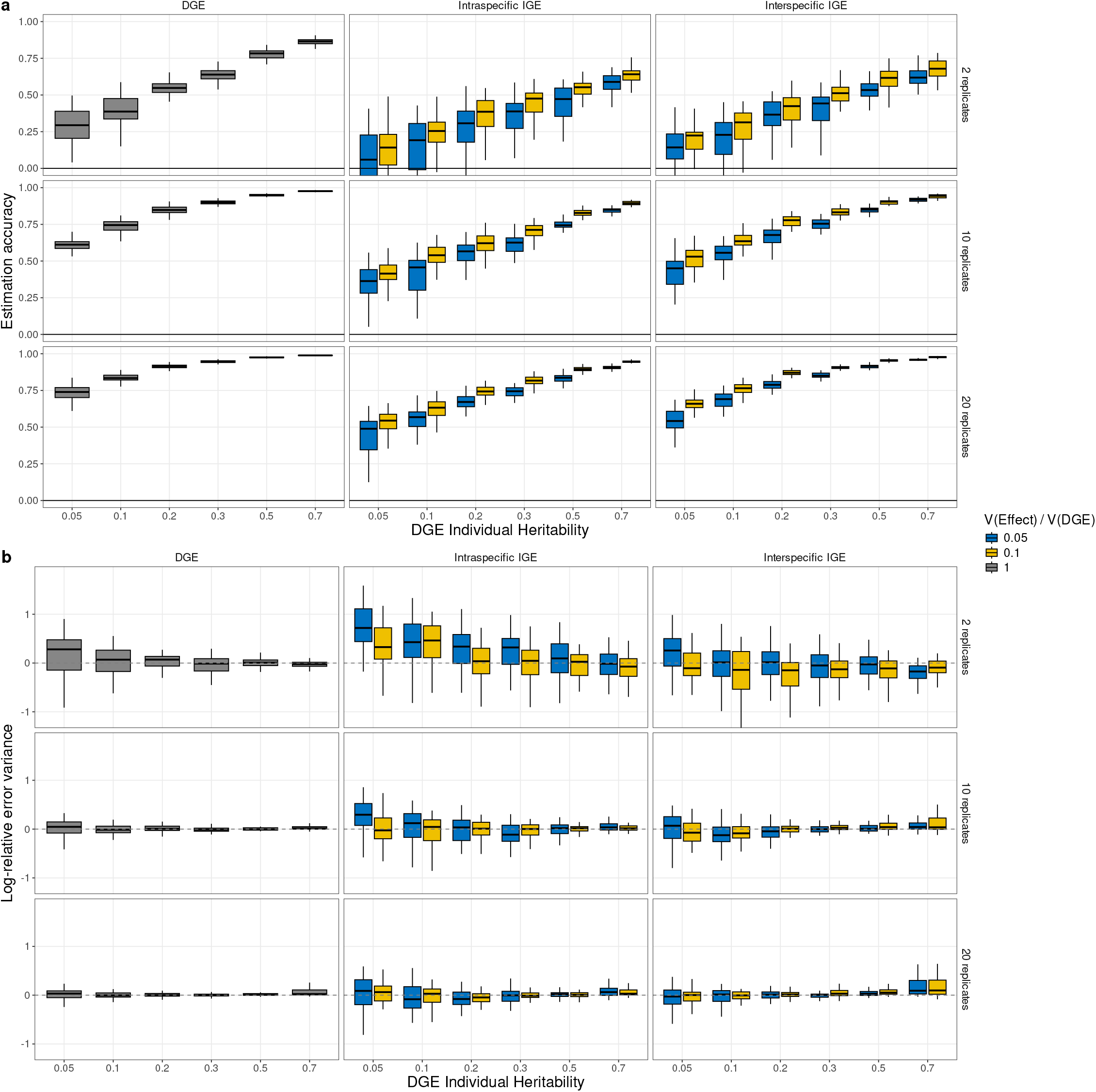
Estimation accuracies (a) and errors (b) obtained by simulating two interacting species in the case of varying levels of plant-plant intraspecific interactions. No indirect environmental effect is simulated in this figure (*V*(*IEE*) = 0), so the environmental effects reduce to the direct term (contained in the residual) and no DEE-IEE covariance is present in these data. DGE, direct genetic effect; IGE, indirect genetic effect; IEE: Indirect environmental effect. DGE heritability is defined as V(DGE)/V(Pheno). The simulated data follow the model given in Equation. 11 (see text). **(a)** correlations between estimated and true genetic values of the different genetic effects DGE, IGE intra and IGE Inter according to the heritability and the number of replicates per genotype, **(b)** the logarithm of the ratio of the estimated over true variance of each genetic effect. Colors indicate the relative ratio of V(IGE)/V(DGE). In column DGE, both V(IGEinter)/V(DGE) and V(IGEintra)/V(DGE) are fixed at 0.05. In column IGE inter, V(IGEintra)/V(DGE) is fixed to 0.05. Reciprocally, in column IGE, V(IGEinter)/V(DGE) is fixed to 0.05. Correlations between all effects were fixed: r(DGE,IGEintra)=-0.6, r(DGE,IGEinter)=-0.6, r(IGEintra,IGEinter)=0.6. Each boxplot represents up to 50 independent simulations with 100 genotypes per species, runs that did not return a complete genetic covariance matrix being excluded (Section Model fitting and Estimation accuracies). For clarity, the outlier points were removed from the graph.

Regarding variance estimation, the log-ratio *V*_*error*_ = log_10_(*V*_estimated_/*V*_true_) was positive when information was scarce. At *n*_*rep*_ = 2, *V*_*error*_ could reach values close to 1, corresponding to a tenfold overestimation, and it affected the intraspecific IGE most, which is the smallest component and the one carried by the fewest neighbors. This bias fell steeply with replication and with heritability, and around 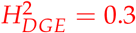 the variance components were recovered without appreciable bias from ten replicates onwards, the dispersion between simulations narrowing over the same range: the standard deviation of the estimated direct genetic variance fell from 0.34 at two replicates to 0.07 at twenty. At twenty replicates the median *V*_*error*_ was 0.006 for the DGE, 0.025 for the intraspecific IGE and 0.022 for the interspecific IGE, an overestimation of the variance components of 1 to 6 percent, reaching about 0.07, some 15 percent, for the interspecific IGE at the highest heritability.

At the upper end of the heritability range the estimates behaved differently. At 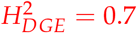 the error on the variance components grew with replication instead of shrinking, and a few fits did not converge. The three genetic components were then no longer estimated independently of one another: from one simulation to the next, the whole genetic block was overestimated or underestimated together, while the variance of the total breeding value stayed close to its true value. Convergence failures and inaccurate variance components of this kind are the signature of a weakly identified model, as (Cantet and Cappa 2008) showed for animal models with competition effects. Separating the variance components is therefore difficult at very high heritability, and adding replicates does not remove the difficulty.

This difficulty does not carry over to the prediction of the genetic effects themselves. The accuracy of the predicted values (BLUPs) keeps improving when 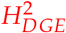 increases (Figure 5): ranking genotypes and splitting the phenotypic variance into its components are two different problems, and only the second one is affected. Overall, these results confirm that the framework performs well even when little IGE variance is expressed relative to DGE, provided that a sufficient number of replicates per genotype is available.

The variance ratio of one indirect effect left the accuracy and *V*_*error*_ of the other effects essentially unchanged. This suggests that the model properly estimates the variances with no or little influence of the magnitude of the other effects (Supplementary Figures S8 and S9).

The correlations between genetic effects had a marked effect on the accuracy of the IGEs. A strong DGE-IGE correlation improved the estimation of the IGE, and the gain was larger for the intraspecific IGE, which is the hardest effect to estimate: the joint model transfers information from the well-estimated DGE towards the indirect effect it covaries with (Henderson and Quaas 1976). This transfer was specific, the correlation between the DGE and one IGE leaving the accuracy of the other IGE essentially unchanged. The correlation between the two IGEs, in contrast, had little effect on their estimation, neither of them being estimated well enough to inform the other. None of these correlations changed *V*_*error*_ (Supplementary Section Effect of the genetic correlations on estimation).

#### Effect of modeling indirect environmental effects

Most plant IGE studies do not explicitly include IEE in the statistical models they use to estimate genetic effects (Sato *et al*. 2024; Costa e Silva *et al*. 2017). To assess what this omission costs, we compared two models: one including the IEE as random effects (hereafter the *IEE model*) and one omitting them (hereafter the *null model*). We fixed the following parameters: 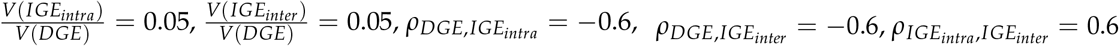.

Failing to account for IEEs leads to confounding between environmental and genetic neighbor effects, leading to an overestimation of the IGE variances, while the DGE estimates remain unbiased. When IEEs are present (IEE variance ratio *>* 0), the IEE model yields higher BLUPs estimation accuracy and lower *V*_*error*_ for both intraspecific and interspecific IGEs compared to the null model, particularly when the number of replicates is sufficient (*n*_*rep*_ *>* 2; Figure 6). No meaningful differences were observed between the two models for DGE estimates, indicating that the direct genetic effects were robust to the omission of IEEs. When IEEs are absent (IEE variance ratio = 0), the IEE model does not show any detectable loss of accuracy or increase in *V*_*error*_ for any genetic effect relative to the null model (Figure S10). The added term improves the estimates only when a non-negligible share of the environmental variance is actually carried by the neighbors. Taken together, these results suggest that systematically including IEEs in the model is a conservative and robust strategy that improves IGE estimates when IEEs are real, without compromising the accuracy of genetic effect estimates.

**Figure 6.**
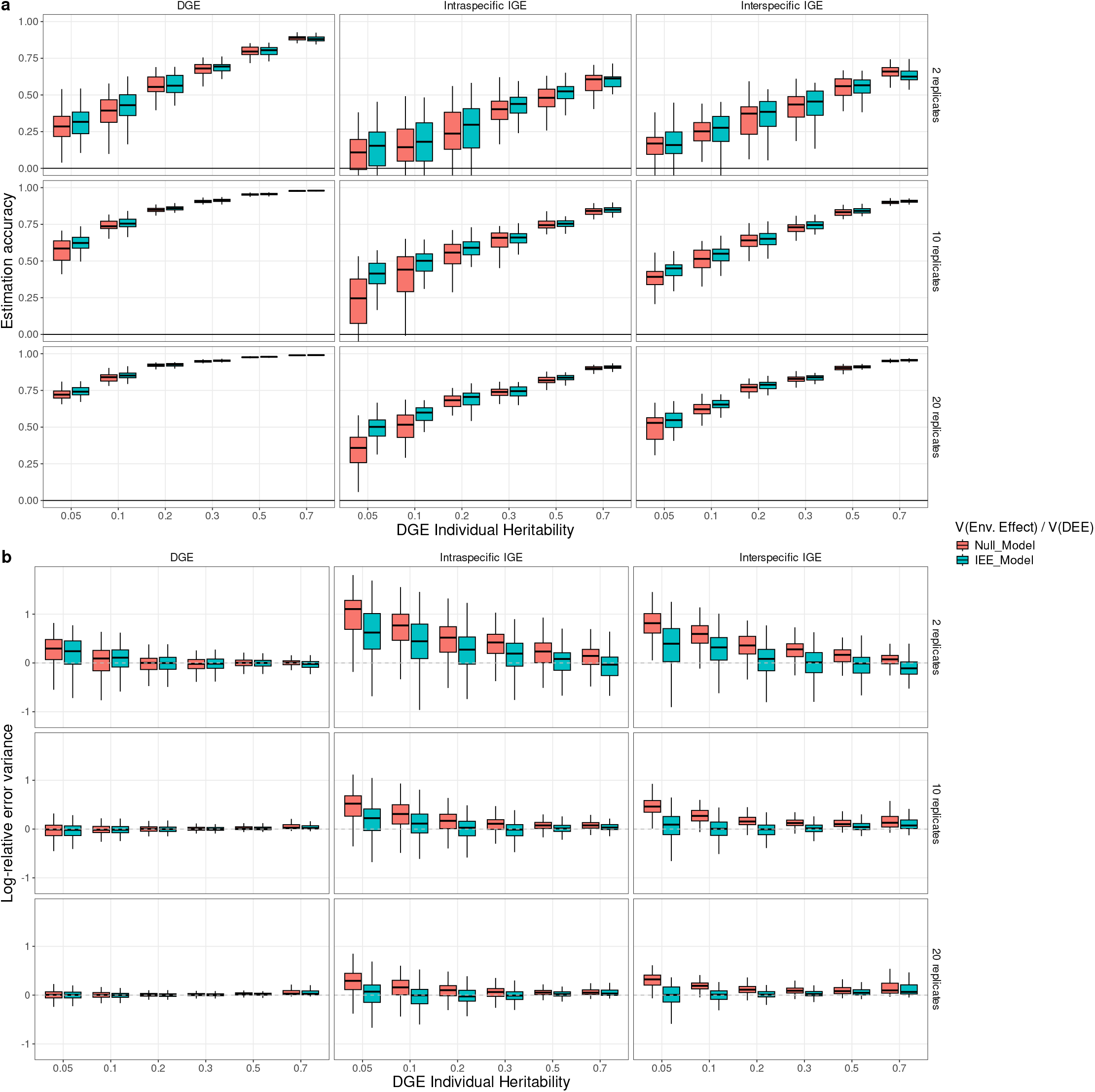
Estimation accuracies (a) and errors (b) obtained by two different models accounting explicitly or not for IEE on simulation of two interacting species. DGE, direct genetic effect; IGE, indirect genetic effect; DGE heritability is V(DGE)/V(Pheno); IEE: Indirect environmental effect. The simulated data follow the model given in Eq. 11 (see text). **(a)** correlations between estimated and true genetic values of the different genetic effects DGE, IGE intra and IGE Inter according to the heritability and the number of replicates per genotype, **(b)** the logarithm of the ratio of the estimated over true variance of each genetic effect. The colors indicate the model used for inference. The IEE model corresponds to the model presented in 13 and the null model corresponds to the same model without the IEE-related terms. All pairwise correlations between IGE and DGE are fixed at -0.6. The correlation between the two IGE is fixed at 0.6. All the IGE over DGE variances ratios are fixed to 0.05. Both indirect environmental effects were simulated with a fixed variance ratio of 0.1 compared to the variance of the direct environmental effect. Each boxplot represents up to 50 independent simulations with 100 genotypes per species, runs that did not return a complete genetic covariance matrix being excluded (Section Model fitting and Estimation accuracies). For clarity, outlier points were removed from the graph. The effect of the choice of the estimation model on the accuracy and the log-relative error of the variance in absence of IEEs is represented in S10

### Computational time

The cost of the model is set by the block of correlated genetic effects: a conventional IGE model carries two of them, a direct effect and an intraspecific IGE, whereas the model proposed here carries three, the second species adding an interspecific IGE. The size of that block is set by the number of genotypes.

The time per fit of the full model grows close to the cube of the number of genotypes, with an empirical exponent of 2.9 over the last doubling, against about 1.1 for the conventional model (Figure 7). The interspecific effect is not a fixed overhead: comparing the two-species models, which differ by that effect alone, it multiplies the time per fit by 1.5 at 100 genotypes per species, by 4.4 at 200, by 11 at 400 and by 22 at 800. The absolute values nonetheless remain moderate, 30 seconds at 400 genotypes per species and under four minutes at 800 on a single processor core and less than 8Gb of memory.

**Figure 7.**
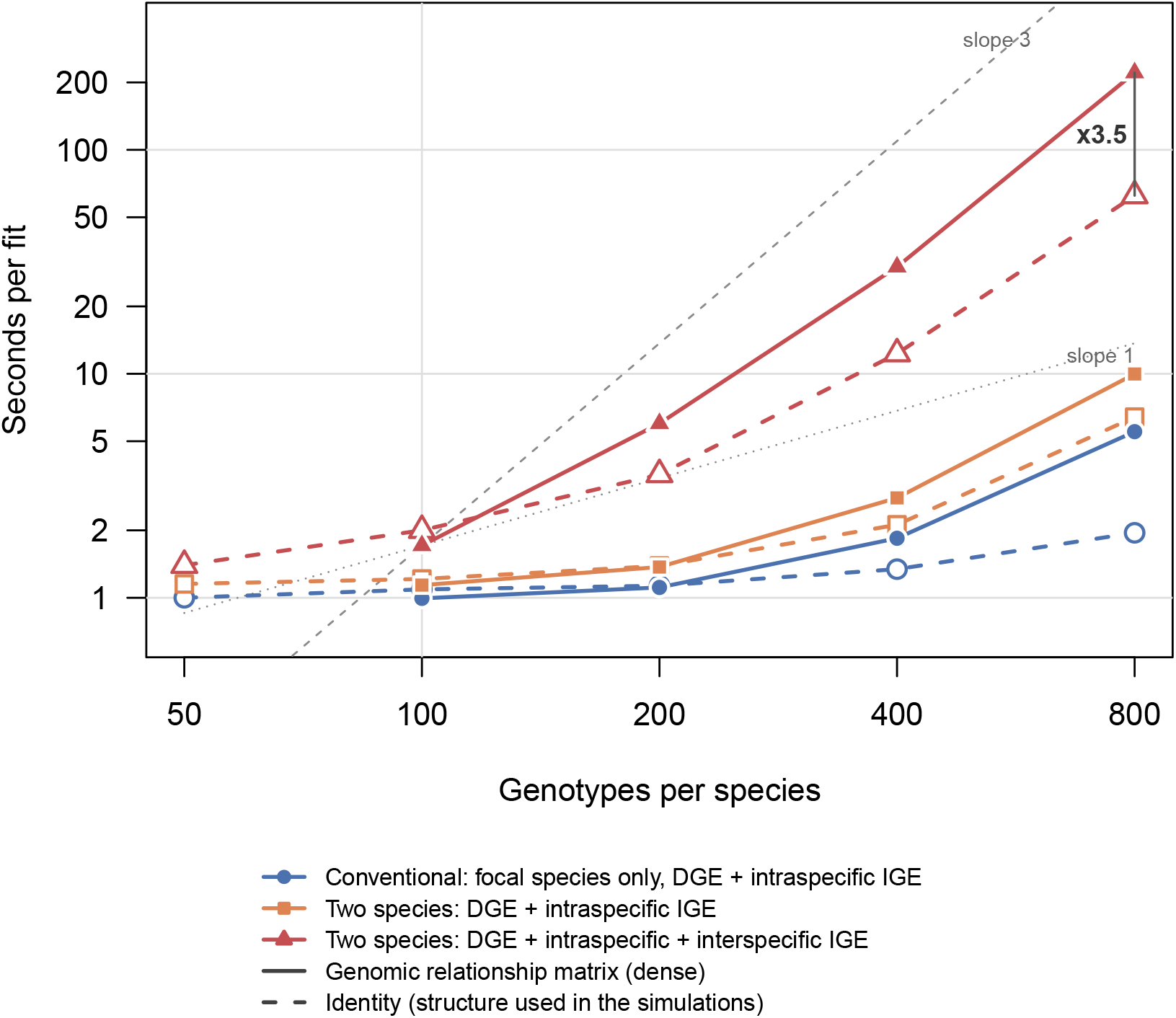
Time per fit against the number of genotypes per species, on logarithmic axes, so that a power law appears as a straight line and its exponent as a slope. Each model is fitted with both genotype covariance structures: a dense genomic relationship matrix (solid lines, filled symbols) and the identity matrix (dashed lines, open symbols). Reference lines of slope one and three are drawn from the first point of the full model fitted with the relationship matrix. The number of plants is held fixed at 8000.

### Comparison with a pairwise design

The direct genetic effect is predicted equally well by the two designs, the two curves being indistinguishable over the whole range (Figure 8). The designs qualitatively diverge as regards indirect effect estimation, which are informed by the neighbors with which a genotype interacts. A pot exposes one genotype to only a single partner so the continuous neighborhood predicts indirect effects markedly better.

**Figure 8.**
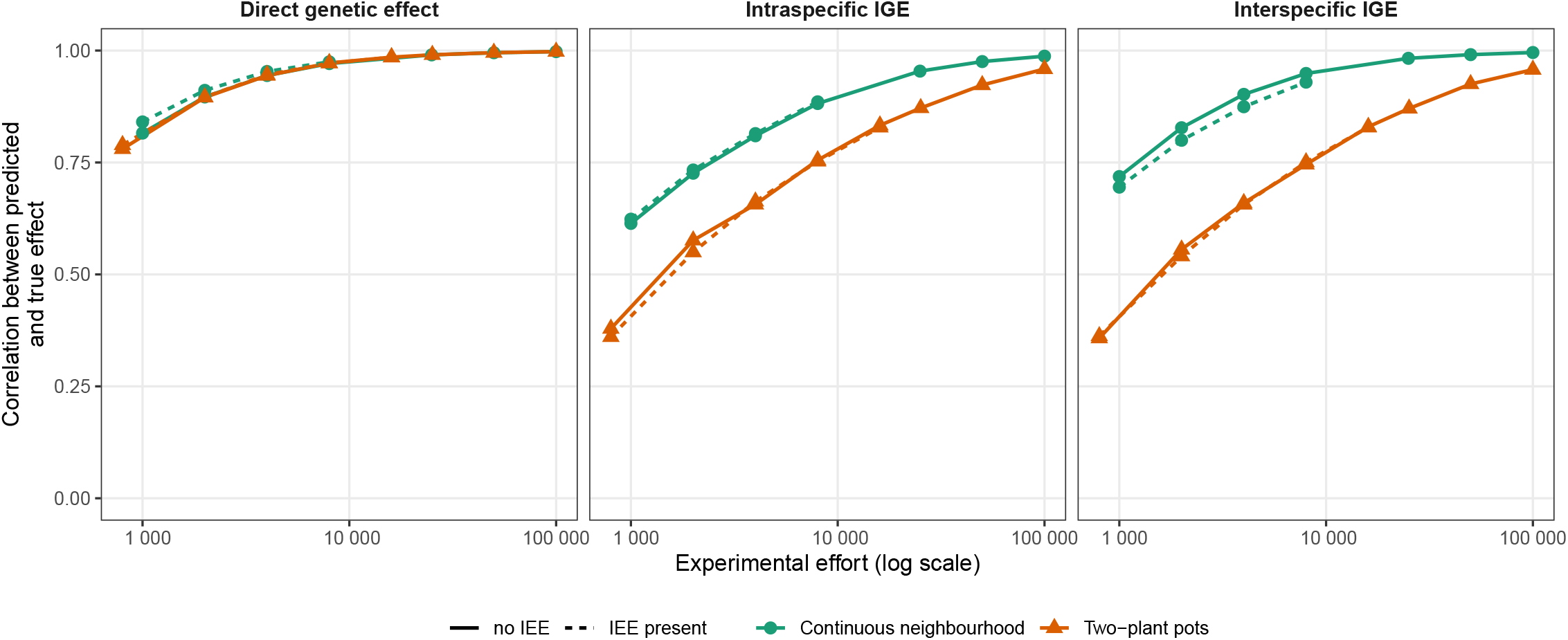
Accuracy of the predicted genetic effects for the two designs, against the number of plants grown, for the direct genetic effect and for the two indirect genetic effects. Every plant is measured in both designs, so this count is also the number of phenotypic measurements. Solid lines: no indirect environmental effect. Dashed lines: indirect environmental effects present in the data, and fitted in the continuous neighborhood, the only design in which they are estimable. Simulation parameters, identical for both species: 100 genotypes per species, 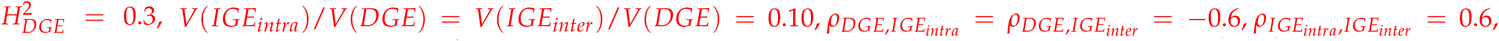, and, where indirect environmental effects are present, *V*(*IEE*_*intra*_)/*V*(*DEE*) = *V*(*IEE*_*inter*_)/*V*(*DEE*) = 0.1. The dilution factor is zero and no genotype-by-genotype interaction is simulated. The genetic effects are fitted with an unstructured covariance matrix. Each point is the median of 40 independent simulations.

Two mechanisms contribute to this gap in Figure 8. A genotype of the continuous grid grow in the vicinity of several partners at once, so the same number of plants encompasses many more genotype combinations, which is what enables the estimation of an indirect effect. Each plant is also exposed to eight neighbors instead of one, so the indirect variance it carries is larger relative to the residual. Rescaling the weights so that every plant receives the same total indirect variance removes the second mechanism (Figure S11): the direct effect is unchanged and the continuous neighborhood design loses most of its advantage on the two indirect effects. The ranking nonetheless holds at every size, and since accuracy rises slowly with the number of plants, that small vertical distance costs a great many plants: the pairwise design reaches at 16,000 plants the accuracy the continuous neighborhood reaches at 8,000, and at 50,000 the accuracy it reaches at 25,000.

The pairwise design shows no such cost, and this follows from its structure rather than from any robustness. In a two-plant pot each plant has a single neighbor, so the indirect environmental effect it receives is one deviation per plant: it is indistinguishable from the residual and merely adds to it, which is also why it cannot be estimated there. Ignoring it therefore costs almost nothing, beyond a slightly larger residual. In a continuous neighborhood the same effect is shared between plants whose neighborhoods overlap, so it is no longer residual-like: it can be estimated.

### Estimates from the semi-controlled experiment

A total of 3,573 phenotypes entered the analysis, out of 3,840 positions: 1,755 for wheat grain number and 1,818 for alfalfa biomass. Plants without a phenotypic value were kept in the neighborhood matrices: one that failed or died still occupied its position and still acted on the plants around it, so it contributed no observation but remained a neighbor.

The neighborhood the data selected (i.e. lowest AIC) was not the adjacent one. At first rank no IGE could be distinguished from zero. The effect of alfalfa on its conspecific neighbors emerged at rank 2 and its effect on wheat at rank 3, both remaining supported up to rank 10, while the AIC kept falling until rank 6 (Supplementary Section Neighborhood order in the semi controlled experiment).

The estimates that follow, and Figure 9, need two conventions to be read. A direct effect is what a plant’s own genotype contributes to its own phenotype. An indirect effect can be read in two ways: as what a plant receives from its whole neighborhood, which is the scale of the variance shares reported here and in Figure 9, or as what a single plant emits, which is the scale of the ratios *α*_XY_ given further down and is much smaller. Every share is expressed on the phenotypic variance of the receiving trait.

**Figure 9.**
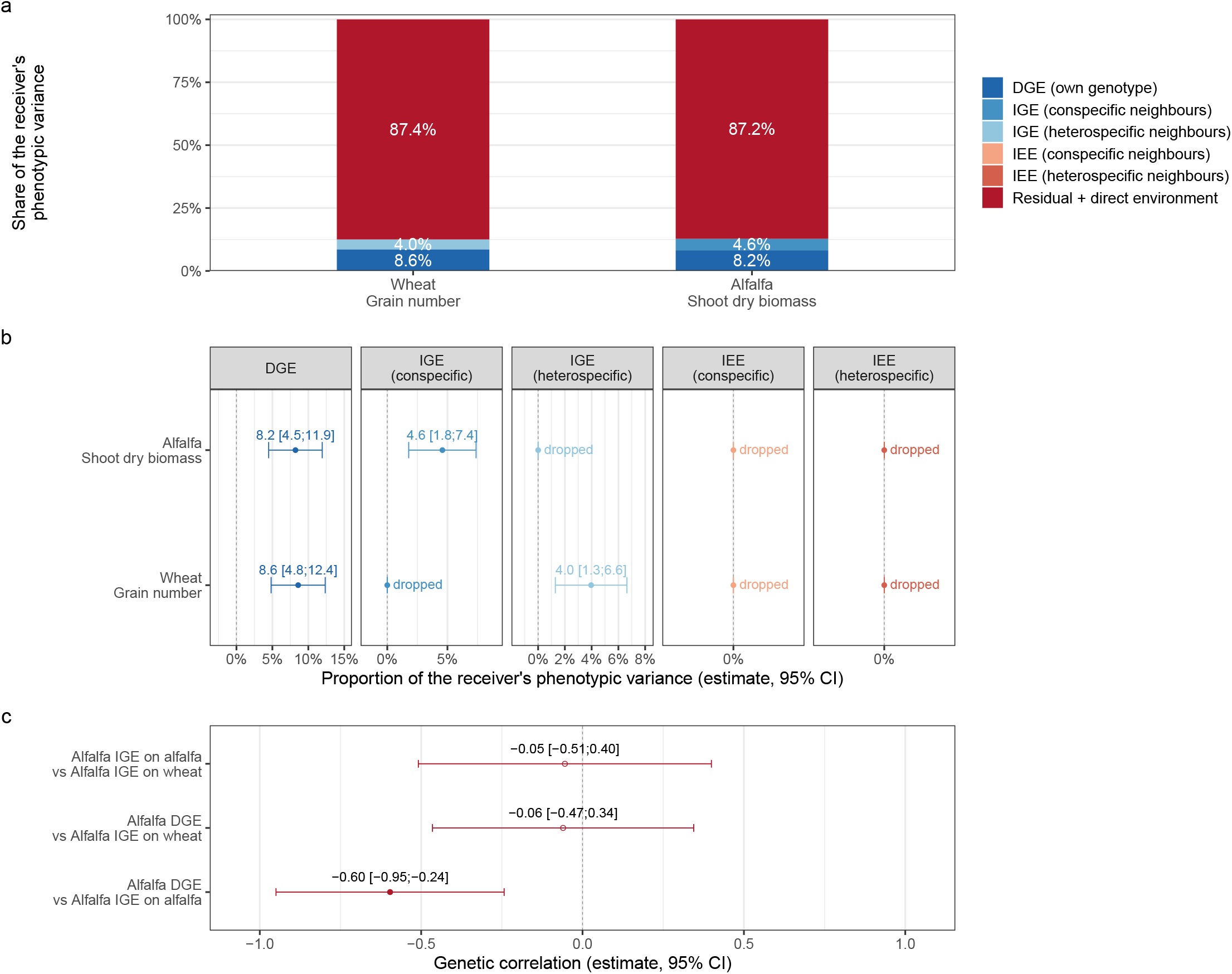
Variance components estimated on the wheat-alfalfa semi-controlled experiment, at neighborhood rank 6. DGE, direct genetic effect; IGE, indirect genetic effect; IEE, indirect environmental effect. Conspecific neighbors carry the intraspecific effects and heterospecific neighbors the interspecific ones. Every share is expressed on the phenotypic variance of the receiving trait, e.g., the interspecific effect shown on wheat is the one emitted by alfalfa genotypes and read on wheat grain number. **(a)** full partition of the phenotypic variance of each receiving trait, summing to 100 per cent **(b)** the same components as point estimates with intervals of *±*1.96 standard error. Effects estimated at zero are denoted as “dropped” (c) correlations between the genetic effects retained in the model, with intervals of *±*1.96 standard error; filled points depart from zero.

The direct genetic effect accounted for 8.6 % of the total phenotypic variance of wheat grain number (*z* = 4.47) and 8.2 % for alfalfa biomass (*z* = 4.32). Two indirect effects were supported by the data, both emitted by alfalfa, one on each species: alfalfa acting on alfalfa accounted for 4.6 % of the variance of alfalfa biomass (*z* = 3.21), and alfalfa acting on wheat for 4.0 % of the variance of grain number (*z* = 2.90). Each species therefore carries one indirect effect in its phenotypic variance, which is how Figure 9 arranges them, although both are emitted by alfalfa. The two effects emitted by wheat were estimated at zero and dropped from the model, as were the indirect environmental effects of both species.

Per emitting individual, in adjacent-contact units (Supplementary Section General neighborhood structure), these shares correspond to ratios of *α*_intra_ = 0.135 and *α*_inter_ = 0.121 for alfalfa. A small effect per emitting plant is therefore enough to reach a few per cent of the phenotypic variance of the receiving species once it accumulates over a whole neighborhood. The experiment lies close to *α*_*XY*_ = 0.1, the value at which the accuracies of Figure 5 were obtained. The genetic correlation between the direct and intraspecific indirect effects of alfalfa was negative, −0.60 (95 % CI −0.95 to −0.24). None of the other genetic correlations departed from zero.

## Discussion

This study proposes the first variance-based quantitative genetic framework that simultaneously models intra- and interspecific indirect genetic effects in plant communities. Since (Bijma *et al*. 2007b), variance-based IGE models have quantified the heritable component of social interactions and predicted responses to selection, but almost exclusively in intraspecific contexts (Cappa and Cantet 2008; Costa E Silva and Kerr 2013; Montazeaud *et al*. 2023; Sato *et al*. 2024), their extension to interspecific interactions having remained limited to plot-level approaches without intraspecific diversity within the mixture (Haug *et al*. 2023). Here the phenotype of a focal individual depends on its own direct effects and on the indirect genetic and environmental effects of the neighbors of both species (Eq. 2), and two quantities follow that have no equivalent in previous models: a total breeding value decomposed into what an individual contributes to its own species and to the associated one, and an interspecific relative heritable variance ratio. Its parameters can be estimated from precisely georeferenced individual observations of two co-cultivated species.

### What the semi-controlled experiment shows

The framework was built for individually georeferenced plants of two species grown together, and the empirical experiment shows that such data can be produced and analysed. On a wheat-alfalfa mixture in which every plant was measured individually, the model separated the direct genetic effects of both species from the effects each exerts on the other, and it attributed shares of the same order to the direct and to the indirect terms. On these traits, indirect genetic effects are not a marginal component.

On the two traits related here, both indirect effects were emitted by alfalfa, on its own species and on wheat, whereas none of the indirect effects emitted by wheat could be distinguished from zero. The two species do not play symmetrical roles in this mixture, but many more traits have been recorded in this experiment and will form the basis of a planned future publication. The objective here was to demonstrate the feasibility of such a design: 1920 plants of two different species were georeferenced and regularly phenotyped. These preliminary results should therefore not be taken as a definitive study of this system.

The two species also differed in the nature of their genetic material. Wheat genotypes were pure lines, so the genotypic effect fitted for wheat gathered the whole G component of each line, additive and non-additive alike. Alfalfa genotypes, in contrast, were half-sib families, and the family effect fitted for alfalfa therefore captured only one quarter of the additive variance (Gallais 2003). Marker information on all alfalfa individuals would have given the genomic kinship between them, and the whole additive variance could then have been estimated. For wheat, adding a genomic relationship matrix, and an epistatic one derived from it, would allow the additive part of G to be separated from the additive-by-additive component. This experiment therefore shows that the framework can be applied to two species with different genetic structures at the same time.

Expressed per emitting individual, these ratios fall within the range explored in the simulations, and within the range of the values already reported in the literature (Montazeaud *et al*. 2023; Sato *et al*. 2024; Haug *et al*. 2023; Santostefano *et al*. 2025). The accuracies obtained in that part of the range are therefore realistic expectations for an experiment of this kind: at 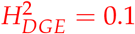, twenty replicates per genotype, *α*_*XY*_ = 0.1 and a correlation of −0.6 between direct and indirect effects, the simulations returned accuracies of 0.85 for the direct effect, and 0.70 and 0.79 for the intra- and interspecific indirect effects.

Finally, these results show that genetic diversity for plant-plant interactions does exist both among and within species in wheat and alfalfa, in conjunction with more in-depth studies carried out on plants. Our contribution here is to provide a new formal framework for the analysis of continuous neighborhoods, which proves effective at detecting these indirect effects.

### What the framework estimates, and at what experimental cost

Within the parameter space explored in the simulations, we confirmed that the framework and its associated statistical modelling separates the direct and indirect genetic effects of two interacting species. Validating this through simulation is a necessary step for any new quantitative genetic model, since analytical derivations guarantee mathematical consistency but not statistical estimability under realistic experimental conditions (Muir 2005; Cappa and Cantet 2008). Our results suggest that ten replicates per genotype, i.e., about 2,000 plants for 100 genotypes per species, is a baseline for reaching moderate accuracy on low-heritability interaction traits, and that fewer are needed as heritability increases. This order of magnitude matches what has been reported for intraspecific IGE models in plants: Montazeaud *et al*. (2023) achieved comparable accuracies in *Arabidopsis thaliana* with a similar number of replicates, and Cappa and Cantet (2008) noted that IGE estimation in forest trees requires large experiments precisely because indirect effects contribute little variance relative to direct ones. The spatial layout is not neutral either: since the number of neighbors of each species dictates how much information each indirect effect receives, breeders should tailor the neighborhood design of their experiments to the effects they wish to prioritise.

A key feature of the model is that all effects are treated as random and drawn from multivariate normal distributions, so that a complete variance-covariance matrix is estimated across all traits simultaneously, within and between species. By leveraging genetic correlations, the model uses the precisely estimated DGE, which benefits from the strong signal of the individual’s own genotype, to partially recover the less precisely estimated IGEs (Henderson and Quaas 1976). In our simulations the DGE variances were substantially larger than the IGE variances, as reported in the empirical literature for both plants and animals (Montazeaud *et al*. 2023; Ellen *et al*. 2014; Muir 2005); but non-zero DGE-IGE covariances consistently improved IGE prediction accuracy, and the improvement grew with replication number. This is where joint multivariate estimation matters most: in systems where indirect genetic effects are weak.

The variance components were recovered accurately over the range of heritabilities tested 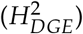 ), and they deteriorate at both ends of that range for opposite reasons. At low heritability and few replications the variance estimates are inflated because the information available on the smallest components is limited; the effect is largest for the intraspecific IGE and it disappears as replication increases, which is the expected behavior of variance component estimation when a component is small relative to the residual (Harville 1977; Visscher 1998). At high heritability the difficulty is of another kind, and the partition rather than the total is what suffers (Section Impact of varying social genetic variance ratios).

This difficulty is documented, and it is not specific to our model. (Cantet and Cappa 2008) encountered it in animal models with competition effects, where the variance components can become impossible to estimate however many animals are recorded, and showed that whether they are estimable at all is a property of the design, of how individuals are distributed and of who interacts with whom, rather than of the amount of data collected. They also note that such situations do not impair the prediction of breeding values, and our results confirm this: the accuracy of the predictions continually improves over the range of high heritabilities. (Bijma 2010a) reaches the same conclusion for indirect genetic effects, their precision being set by the way individuals are grouped rather than by the size of the experiment. From a breeder’s perspective this is the relevant property, since the response to selection depends on a good correlation between predicted and true values rather than on accurate estimations of the variance components themselves.

### Indirect environmental effects and spatial correction

Modelling the non-genetic effect of a neighbor explicitly is unusual in plant IGE studies, where this variance is generally left to the residual or absorbed by a spatial correction (Costa e Silva *et al*. 2017; Silva *et al*. 2013). The two are, however, distinct objects. A spatial structure such as AR1*×*AR1, or a spline, models a smooth autocorrelation surface over field positions: it describes where a plant sits in the field, not which neighbor acts on it. The IEE is a directed effect carried from an identified neighbor to the focal plant through the neighborhood matrix, that is, an emission by one given plant and a reception by another. The covariances they induce differ accordingly: a spatial term makes two plants co-vary as a function of the distance between them, the IEE only to the extent that their neighborhoods overlap. On a regular grid the two are not independent: two plants standing close together necessarily share part of their neighborhood, so the covariance the IEE induces is itself a decreasing function of distance, which is why non-genetic neighbor effects are usually absorbed into a spatial term. Fitting both an IGE and IEE removes from the residual the part of the spatial covariation that the identity of the neighbors explains. Both approaches are therefore complementary: IEE can capture local interactions patterns while autocorrelation effects can take into account more dispersed ones.

### Implications for selection and for evolution

The covariance between the two TBV of a species is, in our view, the most consequential quantity for co-cultivation breeding. It determines whether individuals of species *A* that are genetically beneficial for their own species also benefit species *B*, or whether the two objectives conflict. The sign of these covariances is not a fixed property of the system: it emerges from the relative magnitude of the intraspecific and interspecific IGEs to the DGE and from two genetic correlations that can act in opposite directions. A positive covariance means that selecting for intraspecific performance automatically generates indirect genetic gain in the associated species while a negative one implies a trade-off that no single-species index can resolve (Hazel 1943; PESEK and BAKER 1970; Bourke *et al*. 2021). The correlation between the direct effect of species *A* and its interspecific IGE, *ρ*_*AB*_, links a quantity expressed in the phenotype of *A* to one expressed in the phenotype of *B*, and can be estimated in any design holding the two species together, even though it appears neither in the phenotypic variance of species *A* nor in either *τ*^2^ ratio. The correlation between the two indirect effects of species *A, ρ*_*A*_, requires the intraspecific and the interspecific IGE of the same genotypes to be estimated within a single fit. Pairwise designs can provide it, on the condition that the trials devoted to each interaction type share the same genotypes and are analysed together. The covariance is then estimated between effects expressed in two different trials, so any genotype-by-environment interaction attenuates it.

Because the *τ*^2^ of each species includes variance components from both species, the evolutionary potential of one cannot be assessed independently of the genetic properties of the other. This echoes the concept of community heritability (Shuster *et al*. 2006), according to which heritable variation at the community level emerges from the interactions among members rather than from any single species in isolation. A consequence is that *τ*^2^ can take values outside the classical [0, 1] interval (Bijma 2010a; Bergsma *et al*. 2008). A strongly positive DGE-IGE covariance amplifies the heritable variance in a way that has no equivalent in classical animal models (Ellen *et al*. 2014), as documented in laying hens, where IGEs for survival enabled selection responses that classical models would not have predicted (Muir 2005; Ellen *et al*. 2008). A strongly negative covariance brings *τ*^2^ below *H*^2^ and constrains the variance available for selection. Comparing the two is therefore a useful diagnostic for whether social interactions act as a lever or a brake, and the framework extends across species through 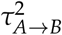. Selection acting on one species may in turn shape the evolutionary trajectory of the other (De Lisle *et al*. 2022).

### Individual measurement and the pairwise design

The contribution of a DGE-IGE model is that it separates what a genotype does for itself from what it does to its neighbors. Using the General Mixing Ability (GMA) framework, IGEs cannot be estimated independently from DGEs, since GMA is the sum of these two effects (Wuest and Niklaus 2018). In a breeding program, the two contributions can be weighted differently through a selection index, e.g., when the two species do not carry the same economic weight. The GMA framework remains an efficient way to evaluate mixture performance and to identify good combinations (Haug *et al*. 2021; Moutier *et al*. 2022; Salomon *et al*. 2026), being analogous to general combining ability in hybrid breeding (Hallauer *et al*. 1988) and the SMA to specific combining ability (Melchinger and Gumber 1998). Estimating the two effects separately presupposes that a phenotype can be attributed to the individual component that produced it.

Plot means suffice when the components can be separated at harvest, as Haug *et al*. (2023) did in pea-barley intercrops by exploiting the difference in grain size; when they cannot, and in particular when genotypes of the same species must be told apart, the aggregate is all that remains. Measuring every plant removes that condition at a price: the phenotype measured on a single plant is less accurate than a mean over many, so more observations are needed for a given accuracy, and every plant has to be precisely located and its neighbors known.

The sessile habit is what makes this possible. Neither the focal plant nor its neighbors move, so distances are fixed for the whole life cycle and each plant belongs to a neighborhood defined before the experiment starts, a stable partition that animal populations cannot provide (Santostefano *et al*. 2025) and which is why that literature works with fixed groups, its precision being set by the way individuals are grouped rather than by the size of the experiment (Bijma 2010a). Forest genetics has exploited this for two decades: Cappa and Cantet (2008) weighted each competitor by an intensity of competition decreasing with distance, and Silva *et al*. (2013) estimated direct and indirect genetic (co)variances for growth and disease damage in a pedigreed *Eucalyptus globulus* population. That literature has also asked which neighbors matter, Costa e Silva *et al*. (2017) identifying the most influential neighbor positions through a factor-analytic structure. To our knowledge, no genetic experiment has combined intraspecific and interspecific interactions, either in animals or in plants. The pairwise unit used in crop mixtures faces the opposite constraint. The number of seeds a combination costs is set by the size of the plot, whereas a continuous neighborhood measures single plants: each of our wheat lines was represented by about ten plants in all, where plots would have called for hundreds of seeds for each combination. Holding the level of measurement fixed and comparing a continuous neighborhood with two-plant pots at an equal number of seeds, the two designs estimate the direct genetic effect equally well and the continuous neighborhood estimates both indirect effects far better (Figure 8). The advantage persists under the rescaling that gives every plant the same total indirect variance whatever its number of neighbors (Cappa and Cantet 2008; Costa E Silva and Kerr 2013), still reaching a given accuracy with about half as many plants. Furthermore, the environmental variation induced by the neighborhood cannot be estimated in a pairwise design since the environmental contribution of a single neighbor of the focal phenotype is indistinguishable from the focal plant’s own residual, whatever the number of pots grown. This could affect the BLUP estimates if these effects are large, through the increase in residual variance. However in our durum-alfalfa case, no IEE were detected on the two reported traits.

The obstacle that Firmat and Litrico (2022) describe is a matter of how combinations in pairwise designs are counted. Having shown that the designs required for community-level genetic prediction lie beyond the reach of most breeding programs, they concluded that breeding strategies should be adapted to the goals perceived and the qualitative nature of the interactions. Continuous neighborhood designs offer an interesting alternative to pairwise designs if a few thousand plants can be measured individually, as demonstrated in our alfalfa-durum wheat case. Estimates of DGE and IGE should be relatively reliable and should enable us to choose high-performing mixtures and the choice of parent lines for the next generation, particularly focusing on alfalfa families that have a favourable IGE on wheat.

If the genetic architecture of plant-plant interactions rested on strong genotype-by-genotype interactions, it would greatly alter the way experimental designs are assembled and their statistical models fitted, compared with the model presented here. Such interaction effects are defined for pairs of specific genotypes, and their estimation requires each pair to be observed several times. While an average indirect effect is informed by any plant neighbouring a genotype, a G x G interaction is informed only by the neighbourhoods that involve a specific genotypic pair. Such pairs are not restricted to different genotypes: when both plants share the same genotype, and because several genetic effects act at once, IGE and DGE x IGE interactions exist and are mostly absorbed into the DGE in only interspecific pairwise designs (Haug *et al*. 2021; Forst *et al*. 2019). Specific designs have been proposed recently, adding supplementary mono-genotypic plots, to disentangle DGE and the sum of IGE and DGE x IGE in an interspecific pairwise design (Salomon *et al*. 2026). In all cases, the number of plants required grows rapidly, which places such designs out of reach of an experiment aimed at screening many genotypes. Here, we assumed these interactions to be small relative to the direct and average indirect effects, which empirical evidence supports in most systems (Haug *et al*. 2023; Sampoux *et al*. 2020), though not everywhere: Mathieu *et al*. (2025) found that the heritable variation in wheat disease resistance was mostly mediated by specific interactions.

### Model assumptions, extensions and perspectives

Our framework rests on a small number of simplifying assumptions, each of which can be relaxed depending on the objective. In the simulations we assumed purely additive genetic effects, which is the appropriate assumption for the material we have in mind, inbred lines in which dominance is not expressed, and for a response that is close to linear. Dominance and epistasis become relevant with heterozygous material such as hybrids or outcrossing species, and traits whose interactions extend over a wider spatial range may also require a non-linear response to density, or to distance from the focal plant. Both can be incorporated in the same formalism (Bijma 2010b; Goldberg *et al*. 1999), as derived in Supplementary Section General neighborhood structure.

More generally, the decomposition of a phenotype into direct and indirect genetic and environmental effects, together with their covariances, is a modelling pattern that recurs at many scales. It appears between neighboring plants, as in our framework, but also between a host and its microbiome, where the host phenotype is decomposed into a host genetic value, a microbial value and a gene-microbe covariance (Week *et al*. 2024, 2025), and even at the intracellular scale, in the mito-nuclear system, where the organellar genome contributes an indirect genetic effect to a trait expressed by the nuclear genome (Joseph *et al*. 2013; Wolf and Wade 2009). The model developed here can therefore be transposed to any system in which the phenotype of one organism is shaped by the genes of an interacting partner of another kind, provided that partners can be identified and that the trait is measured on the focal organism alone.

The main practical obstacle is phenotyping. Applying the framework in the field requires precise spatial identification and individual measurement of every plant, which can make such experiments expensive. In dense multi-species communities the measurement itself is difficult, since neighboring canopies overlap and assigning the harvested material to the right individual introduces a potential source of error. Such errors do not merely add noise: material attributed to a neighbor rather than to the focal plant displaces variation from the direct to the indirect effects. Prediction can help to reduce the number of plants that must be phenotyped: genomic prediction for DGE and IGE, once established on a phenotyped reference population, allows breeding values of new genotypes to be predicted from their genotype alone (Meuwissen *et al*. 2001; Bančič *et al*. 2021). Phenomic prediction also offers a complementary route, using fast proxies such as near-infrared spectra to build prediction models (Rincent *et al*. 2018). High-throughput phenotyping is another avenue to use: UAV imagery with AI-assisted segmentation measures far more plants per hour than a person can, gives access to traits that cannot be scored by hand on every individual, and resolves individuals within a dense canopy, which is the spatial identification the neighborhood matrices require (Duddu *et al*. 2019; Madec *et al*. 2017). Computing time, by contrast, is not limiting at the scales considered here (Section Computational time).

A natural next step is to apply the framework to other experimental data. Cereal-legume mixtures are a suitable test-case, as its agronomic benefits and eco-physiological basis having been documented from the early synthesis of (Ofori and Stern 1987) to more recent reviews on the ecological determinants of intercrop productivity (Bedoussac *et al*. 2015). Several authors have since asked whether dedicated breeding is required for species grown in mixtures (Annicchiarico *et al*. 2019), a question that maps directly onto the effects formalized here: whether genotypes selected for their own performance also perform well as neighbors, and whether the two objectives are genetically correlated.

## Data availability

All the simulation data and R scripts used to perform simulations and construct the graphs in this study are accessible with all the metadata associated at : https://doi.org/10.57745/ZLKTJE.

## Acknowledgments

During the preparation of this manuscript, the authors used Anthropic’s Claude large language models (Claude Opus 4.5 and 4.6, and Claude Sonnet 4.5, accessed via https://claude.ai and the Anthropic API) for four purposes: (i) exploring the scientific literature relevant to the topic of this manuscript, (ii) assistance in drafting and revising the English text of the manuscript through iterative conversational queries and (iii) assistance in writing and debugging R scripts used to generate simulations, publication-quality figures, and (iv) help to validate equation derivations. In all cases, the outputs were critically reviewed, manually edited, and validated by the authors, who took full responsibility for the accuracy, originality, and scientific content of the final manuscript, including the absence of plagiarism. All sources used or suggested by the LLM were independently verified by the authors of this study.

## Supplementary Materials

### General neighborhood structure

The first-rank, distance-independent neighborhood used in the simulations is a special case of the general model. More complex neighborhoods are obtained by changing the neighborhood matrices *Zn*, without any modification of the model itself.

For a focal individual *i*, let *N*_*x*_(*i*) be the set of its neighbors up to rank *x* (or within a given radius), and let *δ*_*ij*_ be the distance between *i* and a neighbor *j*. Each neighbor receives an interaction factor *f*_*ij*_ = *g*(*δ*_*ij*_), where *g* is a decreasing function of distance, for instance *g*(*δ*) = exp(−*λδ*) or *g*(*δ*) = *δ*^−*λ*^. The entries of the neighborhood matrix *Zn* are then the sums of these weights, over the neighbors of each genotype for the genetic effects and over each individual neighbor for the indirect environmental effects, instead of the simple neighbor counts used in our simulations. The first-rank, equally weighted neighborhood used here is the special case in which *N*_*x*_(*i*) contains only the immediately adjacent plants and all *f*_*ij*_ are equal.

The way these weights scale with the number of neighbors follows the dilution model of (Bijma 2010b), in which the indirect effect received from each neighbor is scaled by 1/*n*^*d*^, with *n* the number of neighbors and *d* a dilution factor. Our simulations correspond to rank 1 and *d* = 0, so each neighbor contributes its full indirect effect and the total is the sum over neighbors. Other values recover classical scalings: *d* = 1/2 gives the variance-preserving case, in which the variance of the total indirect effect does not depend on the number of neighbors, equivalent for equal weights to the normalization 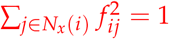 (Cappa and Cantet 2008; Costa E Silva and Kerr 2013; Silva *et al*. 2013). This normalization ensures that the variance of the total indirect effect received by a focal individual, ∑_*j*_ *f*_*ij*_ *IGE*_*j*_, equals the indirect genetic variance *V*(*IGE*) when neighbors are unrelated and non-inbred; without it the estimated indirect genetic variance would depend on the number of neighbors, making comparisons across individuals or designs impossible (Cappa and Cantet 2008). Finally, *d* = 1 corresponds to averaging over neighbors.

Which neighborhood should be retained is an empirical question, and it can be settled on the data: models differing only in *Zn* estimate the same number of parameters and can be compared by likelihood or AIC. The only study we know of that has tested this explicitly reports that the effect of neighbors on herbivory in mixed plantings of *Arabidopsis thaliana* was best captured by the nearest neighbors, the contribution of more distant ranks being small (Sato *et al*. 2024, Supplementary Fig. 3). In our own semi-controlled experiment the answer is the opposite: the indirect effects are undetectable at first order and emerge only from order 2 onwards (Supplementary Section Neighborhood order in the semi controlled experiment). How far a neighborhood reaches is therefore a property of the system rather than of the model, and it is worth fitting a series of ranks and comparing them rather than assuming one.

### Univariate derivation of phenotypic variance

Assuming that the phenotypic value is given by Eq (3) for a single trait per species and that covariances between genetic effects from different species, as well as genetic and environmental effects covariances, are null, the phenotypic variance of individual *i* of species *A* can be expressed as

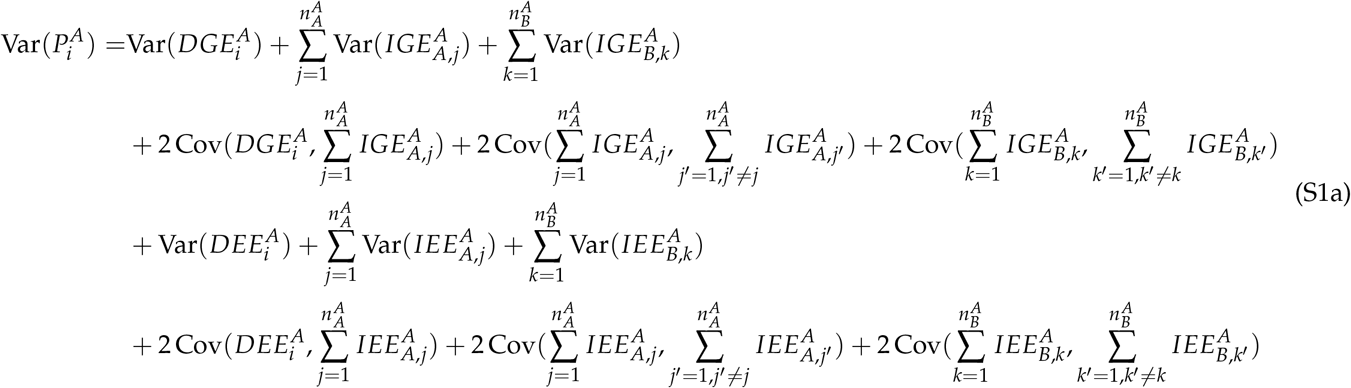

The following assumptions are made:

- All indirect genetic (resp. Environmental) variances are proportional to the direct genetic (resp. Environmental) variance of the species from which they originated.
- The correlations between direct and indirect genetic (resp. environmental) effects are constant for each species in a given generation.
- All plants of each species share common variances and have constant average numbers of neighbors from the same and the other species.
- Genetic relationships between genotypes are modeled by relatedness coefficients 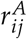 for individuals *i* and *j* of species *A*.
- Environmental relationships between genotypes are modeled by similarity coefficients 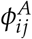 for individuals *i* and *j* of species *A*.
- Genetic effects are defined at the genotype level for each species, while environmental effects are defined at the plant level.
- All the individuals share the same populational genetic (and environmental) variances.

The phenotypic variance for species A can be reformulated as

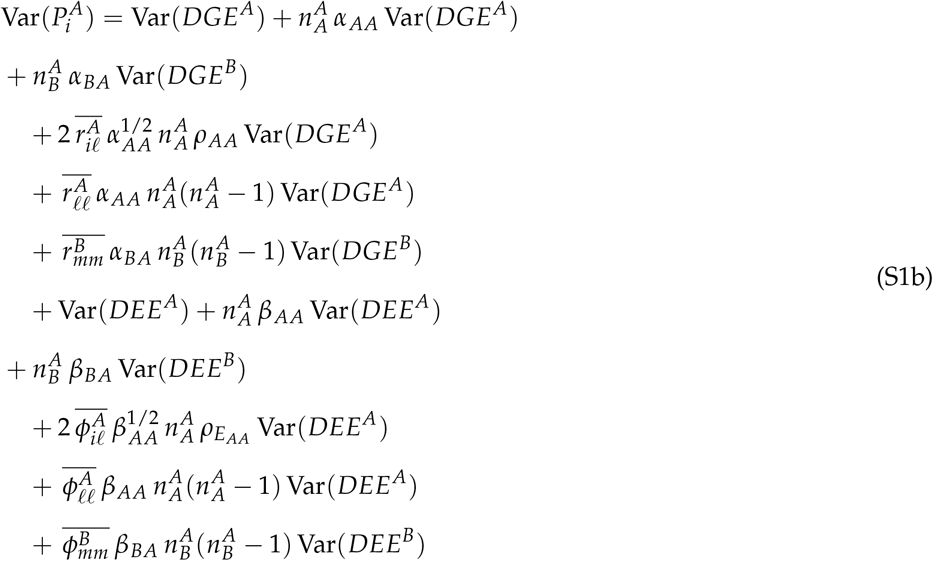

After factorization, this equation becomes

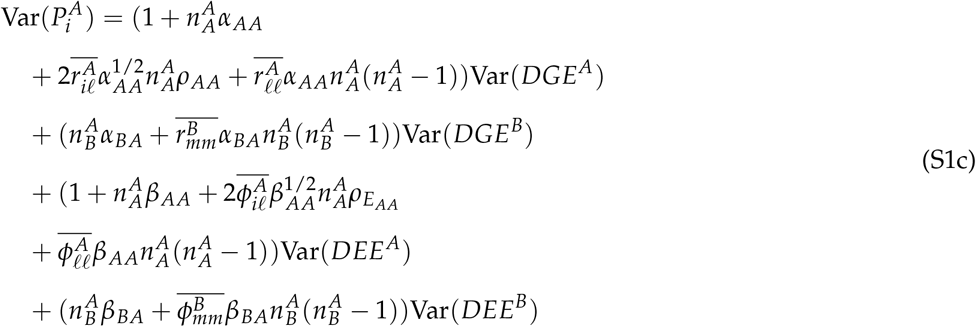

### Empirical variance derivation

The empirical phenotypic variance, denoted as Var(*P*^*A*^ ), can be estimated directly from the observed data for species *A*. Given a sample of *N*_*A*_ individuals from species *A* with measured phenotypic values 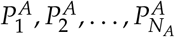.

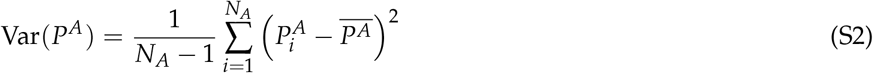

where 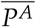 is the sample mean.

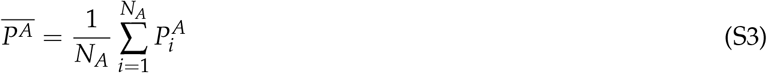

We can first decompose the empirical phenotypic variance as follows.

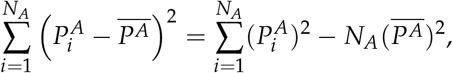

It can therefore be rewritten as

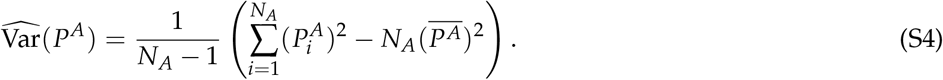

To calculate the expected value of this variance, we can compute the expected value of each term, as the expectation function is additive.

For the first term, we can define for each individual *i* (from the Koenig-Huygens formula) ,

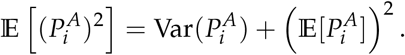

Therefore,

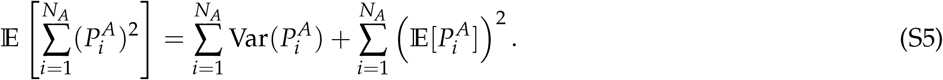

For the second term, we must return to the sample mean formula:

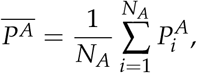

its variance is

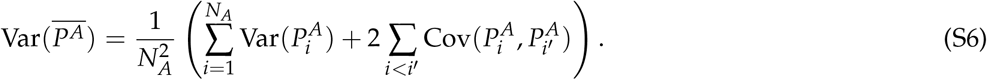

Moreover,

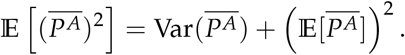

Combining Equations (S4), (S5), and (S6), and noting that

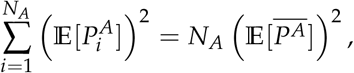

we obtain

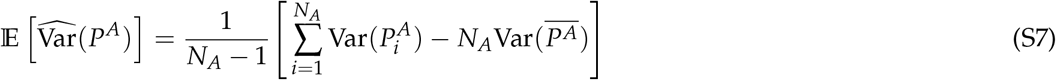

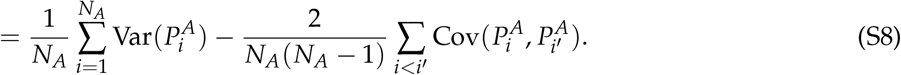

Equation (S8) shows that the expectation of the empirical phenotypic variance equals the average of the individual phenotypic variances corrected by the mean pairwise covariance between distinct individuals. This result is a general result and is not limited to phenotypic variance; it can be applied to any variance estimated from sample data.

### Empirical estimation of phenotypic variance

The empirical phenotypic variance, denoted as Var(*P*^*A*^ ), can be estimated directly from the observed data for species *A*. Given a sample of *N*_*A*_ individuals from species *A* with measured phenotypic values 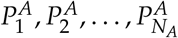 . The expected empirical variance can be expressed as the average of the marginal variances minus the average off-diagonal covariance (derivation in Empirical variance derivation):

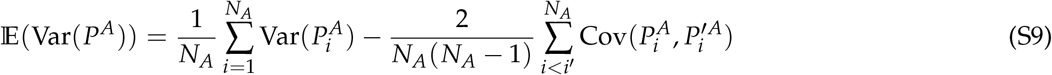

where :

- 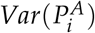 is defined as the phenotypic “population” variance defined in Phenotypic variance

This formulation emphasizes that the empirical variance is determined by the mean of the individual variances corrected for the average covariance between distinct individuals.

#### Phenotypic covariance between individuals

Assuming that all covariances between the genetic and environmental effects are zero, The covariance between the phenotypic values of two individuals *i* and *i*^*′*^ from species *A*, denoted 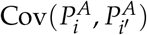, can be written in index notation as

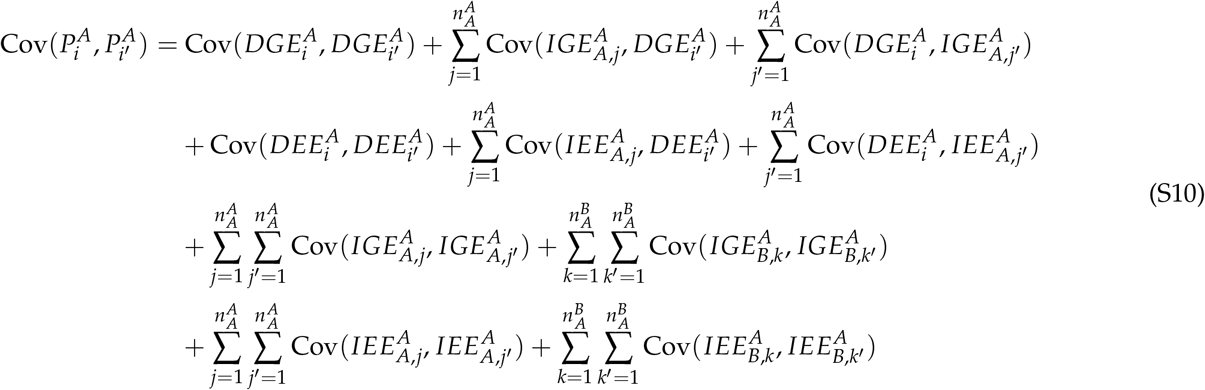

For clarity, we grouped individuals into neighboring sets based on their spatial and social relationships. Specifically, all neighbors *j* of individual *i* (from species *A*) belong to group *ℓ*, while all neighbors *j′* of individual *i′* (from species *A*) belong to group *ℓ*^*′*^. Similarly, all neighbors of individual *i* from species *B* are assigned to group *m*, All neighbors of individual *i′* from species *B* are assigned to group *m′*.

Using this group aggregation, we can simplify the formula as follows.

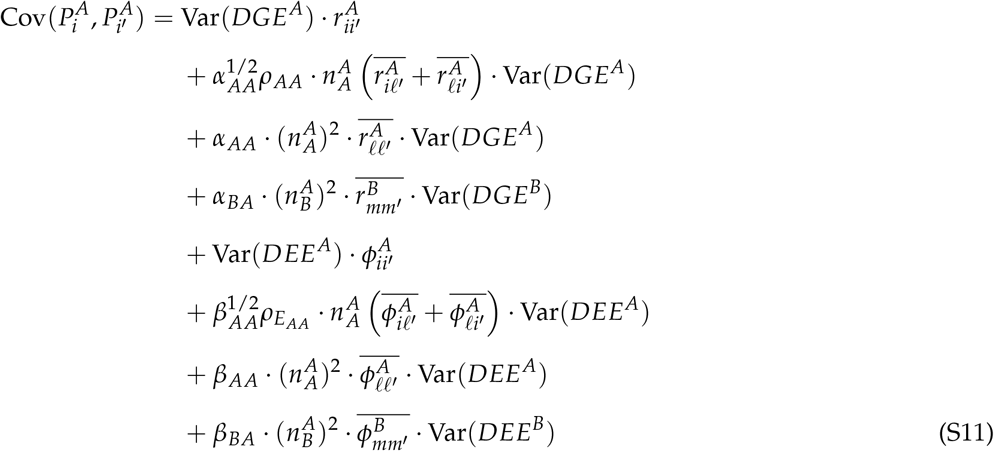

where:

- 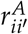: genetic relatedness between individuals *i* and *i′* of species *A*.
- 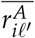: average genetic relatedness between individual *i* and neighbors *ℓ*^*′*^ of *i′* of species *A*.
- 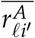: average genetic relatedness between neighbors *ℓ* of *i* and individual *i′* of species *A*.
- 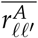: average genetic relatedness between neighbors *ℓ* of *i* and neighbors *ℓ*^*′*^ of *i′* of species *A*.
- 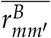: average genetic relatedness between neighbors *k* of *i* and neighbors *k′* of *i′* of species *B*.
- 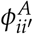: environmental similarity between individuals *i* and *i′* of species *A*.
- 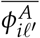: average environmental similarity between individual *i* and neighbors *ℓ*^*′*^ of *i′* of species *A*.
- 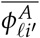: average environmental similarity between neighbors *ℓ* of *i* and individual *i′* of species *A*.
- 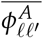: average environmental similarity between neighbors *ℓ* of *i* and neighbors *ℓ*^*′*^ of *i′* of species *A*.
- 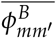: average environmental similarity between neighbors *k* of *i* and neighbors *k′* of *i′* of species *B*.

This expression accounts for all pairwise covariances between the direct and indirect genetic and environmental effects of individuals *i* and *i′* and their neighbors.

#### Expected empirical phenotypic variance

Combining Equations (4), (7), and (9), we can express the empirical phenotypic variance of species A as a function of the variances of the direct genetic and environmental effects from both species, weighted by global coefficients that encapsulate the influence of neighborhood structure and relatedness/similarity patterns.

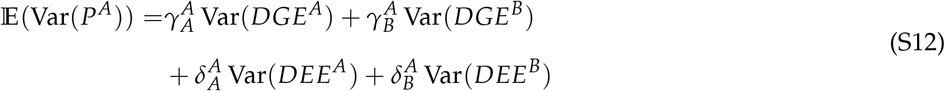

with:

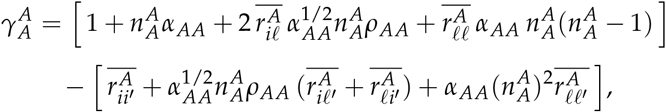

Multispecies IGE Framework for Plant Mixtures

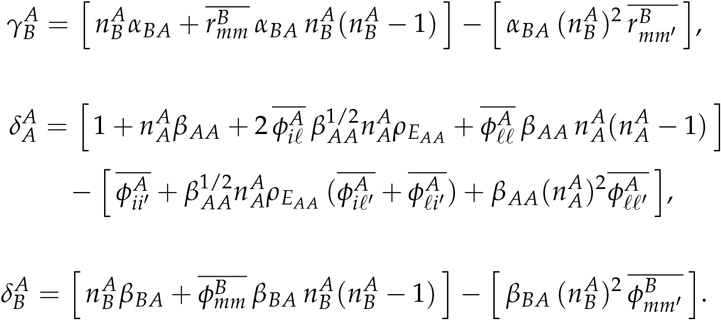

### Empirical estimation of total breeding value variance

As before, we computed empirical variances from the observed TBV samples. For species A, we tracked two TBV types, 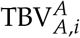 (effects on species A traits) and 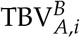 (effects on species B traits). Their sample variances read

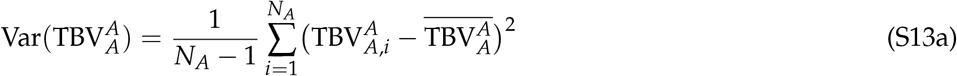

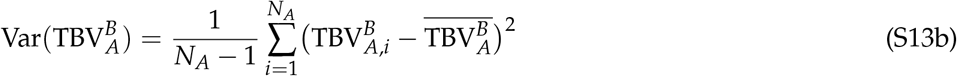

The empirical covariance between the two TBV types is

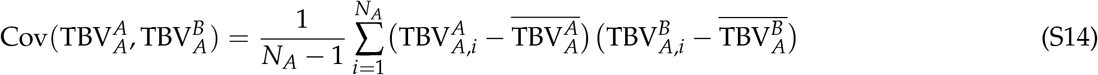

For empirical phenotypic variance, the expected sampling variance is obtained as the average marginal variance minus the average off-diagonal covariance.

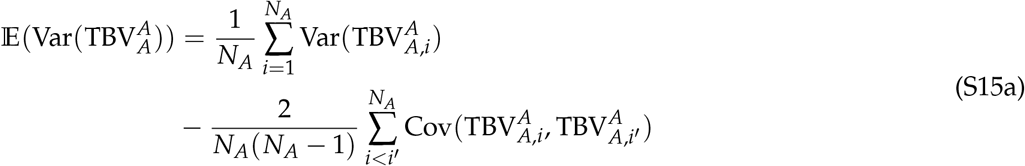

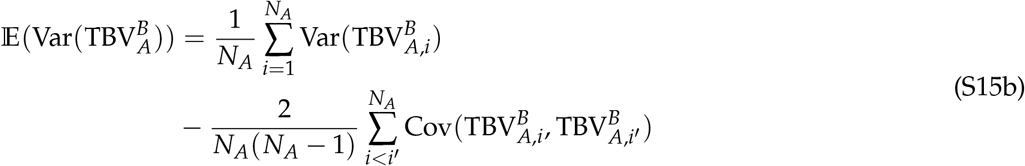

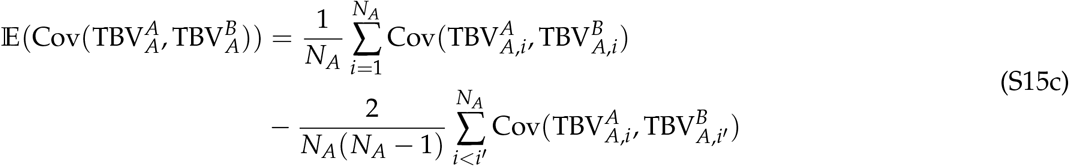

#### Total breeding value covariance between individuals

For one trait per species, we can directly express the parametric equation of the covariances between the TBV for both types as follows:

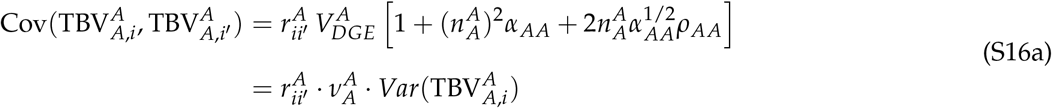

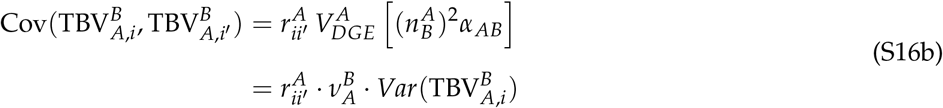

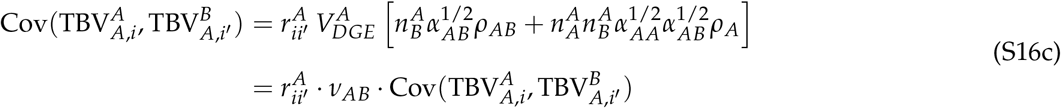

#### Expected total breeding value empirical variance

In summary, we can write the expected empirical variance of the TBV for species *A* as

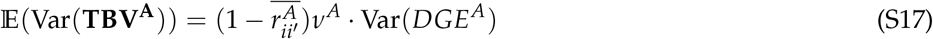

### Remaining incidence matrices of the simulated design

The matrices below complete those given in the main text. They are written for the three plants of species *A* in the central column of Figure 1, whose genotypes are G2, G5 and G8 and whose plant identifiers are ID4, ID5 and ID6.

The neighborhood matrix 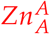 counts, for each focal plant of species *A*, the number of conspecific neighbors carrying each genotype of species *A*, and 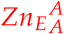 describes the same neighborhood between plant identifiers. In this particular subgrid the two matrices coincide, because the three plants carry three distinct genotypes; they differ as soon as a genotype is replicated within a neighborhood, as it is for genotype G3 of species *B* in the main text.

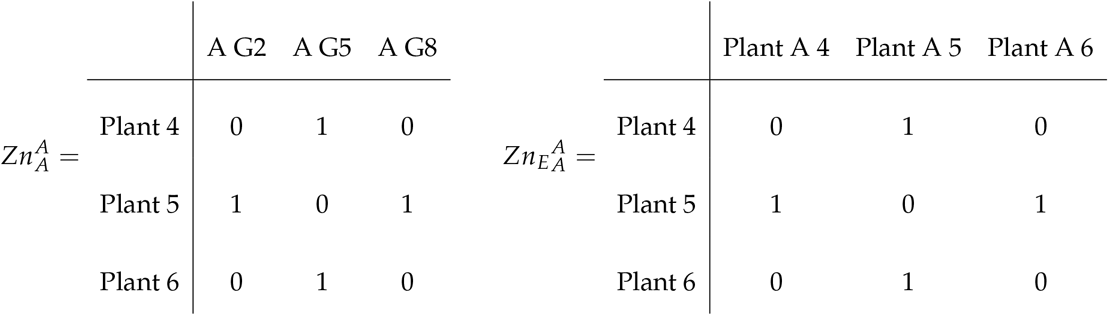

The assignment matrix *Zg*^*A*^ links each plant of species *A* to its genotype, and *Z*_*E*_ ^*A*^ links each plant to its own direct environmental effect, and is therefore the identity matrix of size *N*^*A*^.

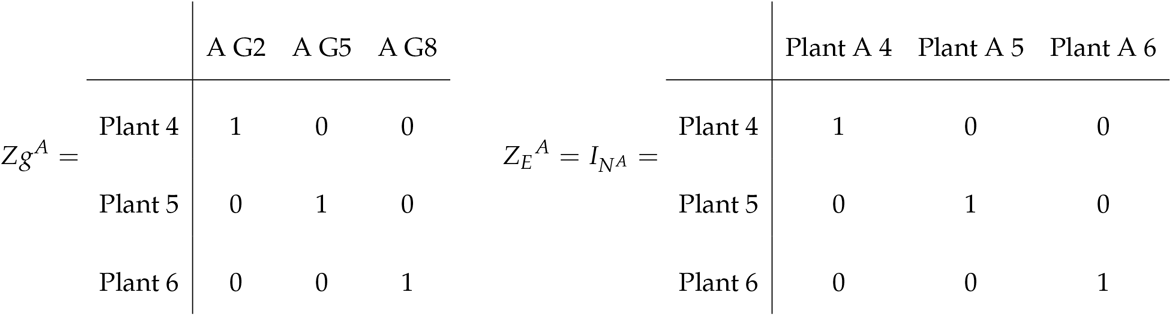

### Direct environmental variance used in the simulations

The variance of the direct environmental effects was obtained by rearranging the definition of the individual heritability of the direct genetic effect (Equation 12), under the assumption that there are no environmental similarities between plants and no indirect genetic variance:

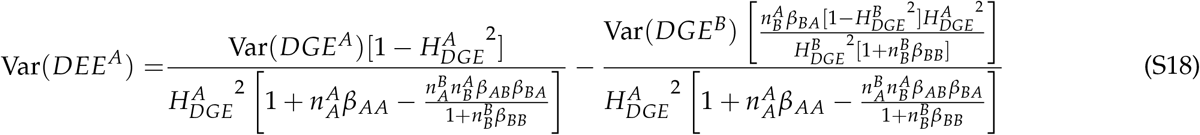

### Genotype covariance structure: kinship or identity

The genotype covariance structure **M** of Equation 13 can be the kinship matrix **K** or the identity **I**. To measure what this choice costs, we ran a dedicated series of simulations: three heritabilities, with the variance ratios fixed at 0.05, ten replicates per genotype and no indirect environmental effect. Each of these datasets was fitted twice, once with each structure, so that the two estimates come from exactly the same data and their difference is not confounded with the variation between simulations. Replicates for which either fit failed to return a complete covariance matrix were discarded from both, which is why the number of simulations per boxplot varies from 48 to 50. The gains reported below are medians of these per-replicate differences, whereas the figure shows the two distributions separately, each of which still carries the variability between simulated datasets. Variance components estimated under **K** were rescaled by 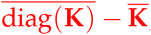, which is the factor relating the additive variance under **K** to the variance of the effects across genotypes, so that both parameterisations are read on the same scale.

The kinship matrix was never the worse of the two, and its advantage shrinks as heritability rises: the median gain in accuracy fell from 0.03 at *H*^2^ = 0.1 to below 0.01 at *H*^2^ = 0.7, where it is no longer distinguishable from zero for the direct effect and the interspecific indirect effect (Figure S1a). This is the expected behaviour, the sharing of information between relatives being most useful when each genotype is poorly informed. On the variance components the ordering reverses with heritability: at *H*^2^ = 0.1 both structures are equally biased, whereas at *H*^2^ = 0.7 the identity inflates the estimated variances by 7 to 16 % while the kinship matrix is essentially unbiased (Figure S1b).

Both departures reduce the performance attributed to the design, so the accuracies reported in this study are a lower bound on what the same experiment would achieve with a kinship matrix. The two structures remain close here because the simulated genotypes are drawn from a single coalescent population with no family structure: scaled to a unit diagonal, the off-diagonal elements of **K** had a mean of −0.01, a standard deviation of 0.06, and never exceeded 0.29 in absolute value over the 150 simulated populations. In an experiment using related material this would no longer hold, and the kinship matrix should be preferred.

**Figure S1.**
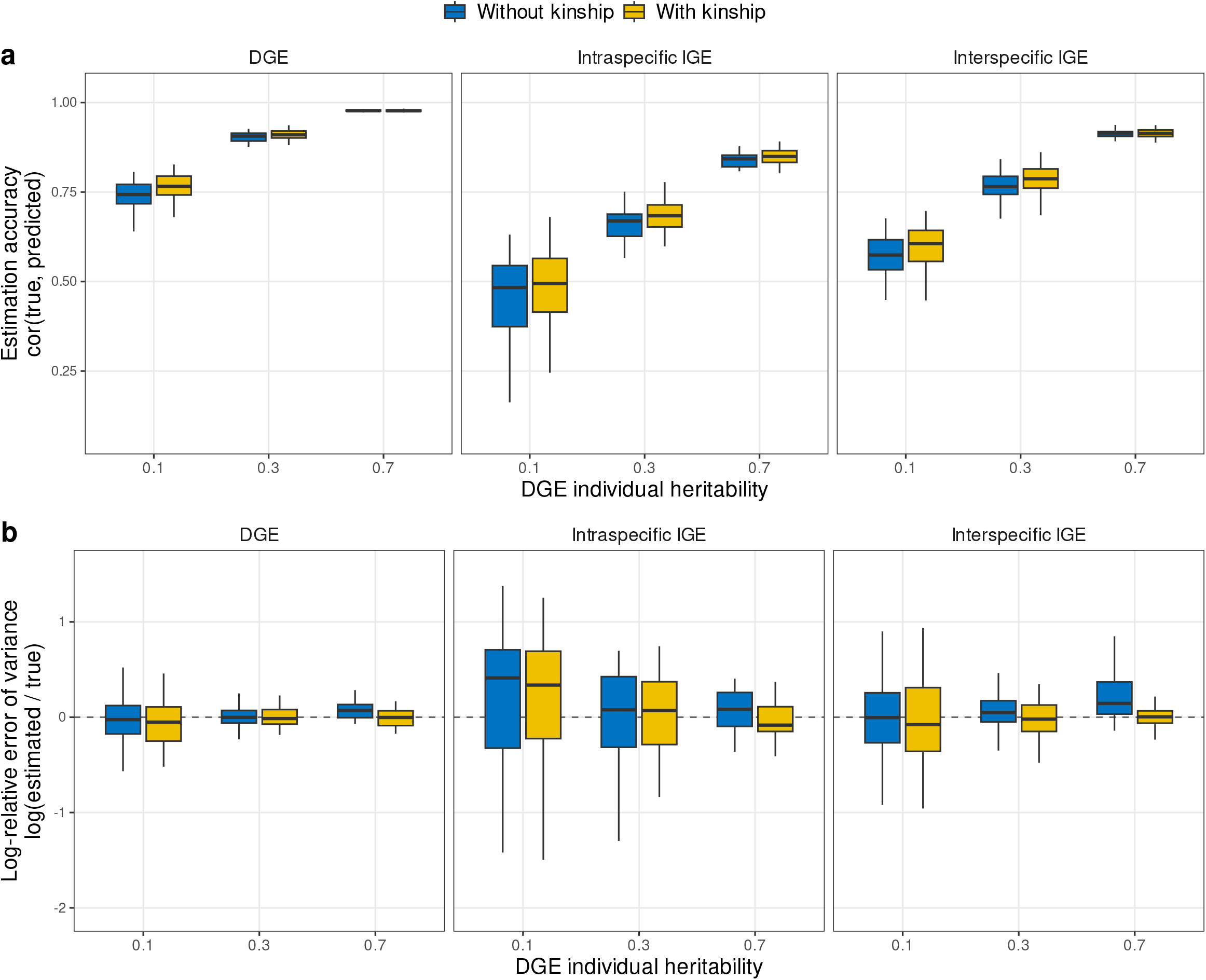
Estimation accuracies (a) and errors (b) obtained by fitting the same simulated datasets with the identity matrix and with the genomic relationship matrix as the genotype covariance structure **M** of Equation 13. DGE, direct genetic effect; IGE, indirect genetic effect. Each dataset was fitted twice, once with **M** = **I** and once with **M** = **K**, so that the two estimates are paired on the same data. **(a)** correlations between estimated and true genetic values of the different genetic effects DGE, IGE intra and IGE inter, according to the DGE individual heritability, **(b)** the logarithm of the ratio of the estimated over true variance of each genetic effect. Colors indicate the covariance structure used for inference. Variance components estimated under **K** were rescaled so that both parameterisations are on the same scale (see text). All the IGE over DGE variance ratios are fixed to 0.05. Correlations between all effects were fixed: r(DGE,IGEintra) = -0.6, r(DGE,IGEinter) = -0.6, r(IGEintra,IGEinter) = 0.6. No indirect environmental effect was simulated or fitted. Each boxplot represents 48 to 50 independent simulations with 100 genotypes per species and 10 replicates per genotype. For clarity, outlier points were removed from the graph.

### Validation of analytical expressions for phenotypic variance, total breeding value variance, and relative heritable variance ratio

The closed-form expressions derived for the phenotypic variance Var(*P*^*A*^ ) (Eq. 3), the total breeding value variance Var(**TBV**^*A*^) (Eq. 6), and the relative heritable variance ratios 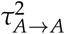 and 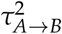 (Eq. 8) are population-level predictions that assume an infinitely large panmictic population. In finite field experiments, however, the empirical variance computed from the observed data is expected to differ from this population value because of finite-sample corrections and non-zero average pairwise relatedness among individuals. These corrections are formalized in the expected empirical variance expressions (Eqs. S12 and S17), which subtract the average pairwise covariance from the mean of the marginal variances.

To verify that these analytical expressions accurately predict the quantities observed in the simulated datasets, we conducted a dedicated simulation study. We generated a factorial grid of 128 parameter combinations by crossing four levels of the intraspecific IGE-to-DGE variance ratio (*α*_intra_ ∈ *{*0, 0.05, 0.1, 0.3*}*), four levels of the interspecific IGE-to-DGE variance ratio (*α*_inter_ ∈ *{*0, 0.05, 0.1, 0.3*}*), two levels of the DGE–intraspecific IGE correlation (*ρ*_intra_ ∈ *{*−0.6, 0*}*), two levels of the DGE–interspecific IGE correlation (*ρ*_inter_ ∈ *{*−0.6, 0*}*), and two levels of the intraspecific–interspecific IGE correlation (*ρ*_*II*_ ∈ *{*0, 0.6*}*). All other parameters were held constant: *H*^2^ = 0.3, *β*_intra_ = *β*_inter_ = 0.05, *ρ*_*E*,intra_ = *ρ*_*E*,inter_ = −0.3, and *ρ*_*E,II*_ = 0.3. For each combination, 50 independent simulation replicates were generated using *n*_geno_ = 100 genotypes per species and *n*_rep_ = 10 replicates per genotype (2000 plants per species) in the same alternating-row field layout as the main simulation study.

For the genetic effects, realistic additive kinship matrices were generated for each species using a synthetic genomic marker panel of 3000 SNPs at a minor allele frequency of 0.2, following the same vanRaden standardization as in the main simulation (Section Genome generation). The mean off-diagonal kinship value 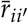 was computed for each replicate and used to correct the population-level theoretical predictions according to Eqs. S12 and S17.

For each replicate, we computed the empirical values of 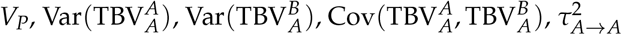, and 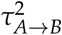 directly from the simulated phenotypes and per-individual TBV values, and compared them with the corresponding theoretical predictions. Figures S2, S3, and S4 show the agreement between the empirical scenario means and theoretical predictions across all 128 parameter combinations. All Pearson correlations exceeded 0.99, and the fitted regression lines (blue) closely followed the identity line (red) across the full range of explored values, including values of 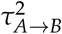 exceeding 1.

Taken together, these results confirm that the closed-form expressions for Var(*P*^*A*^ ), Var(**TBV**^*A*^), and *τ*^2^ are mathematically consistent with the simulation model and provide accurate predictions of the quantities observed in realistically sized experiments across the full range of genetic architectures explored in the study.

**Figure S2.**
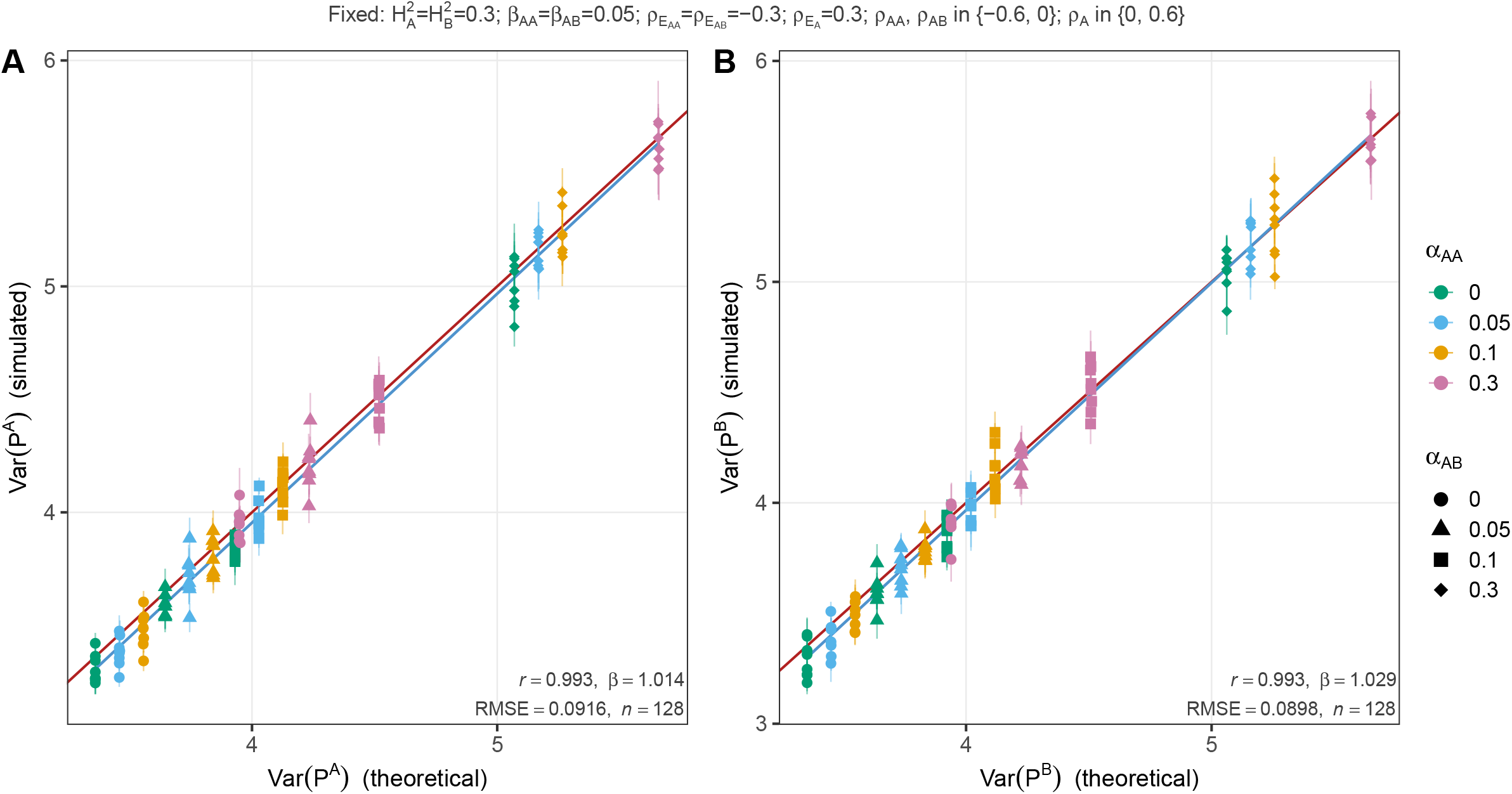
Validation of the analytical expression for phenotypic variance Var(*P*) (Eq. 3) across 128 parameter combinations. Each point represents the mean over 10 simulation replicates for one scenario, with standard error bars on the axes. The red line is the identity (*y* = *x*), and the blue line is the ordinary least-squares regression. Each panel reports the Pearson correlation coefficient *r*, the OLS regression slope *β* (expected to be 1 under perfect agreement), and the RMSE directly on the plot. Point colors indicate the four levels of *α*_intra_ (0, 0.05, 0.1, 0.3) and point shapes indicate the four levels of *α*_inter_ (0, 0.05, 0.1, 0.3); residual scatter within each color–shape group reflects variation in *ρ*_intra_, *ρ*_inter_, and *ρ*_*II*_. **(a)** Species *A*, Pearson *r* = 0.993. **(b)** Species *B*, Pearson *r* = 0.993. (*n* = 128 scenarios).

**Figure S3.**
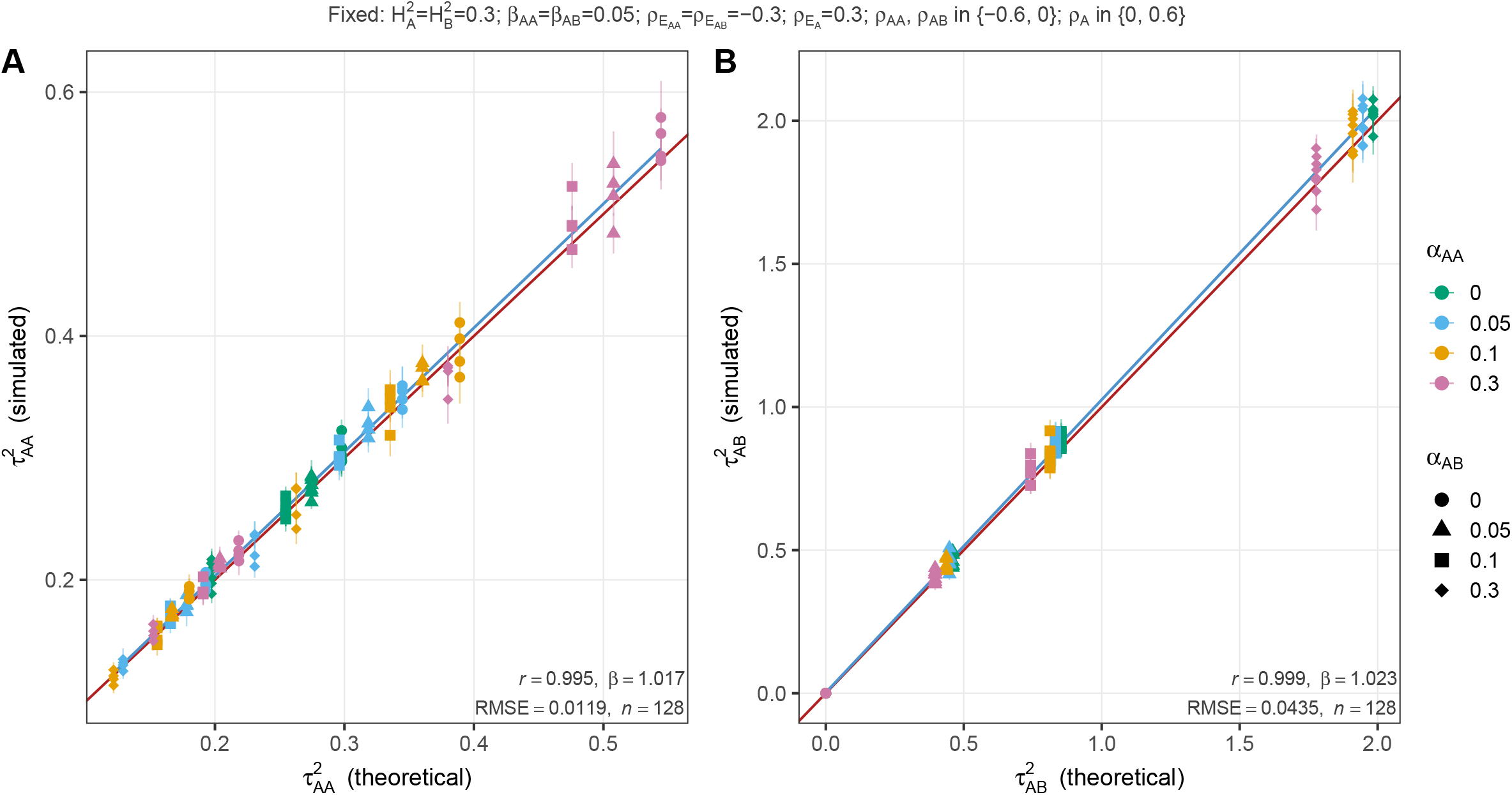
Validation of the analytical expression for the relative heritable variance ratios (Eq. 8) across 128 parameter combinations in the univariate analysis. Each point represents the mean over 10 simulation replicates for one scenario, with standard error bars on the axes. The red line is the identity (*y* = *x*), and the blue line is the ordinary least-squares regression. Point colors indicate the four levels of *α*_intra_ (0, 0.05, 0.1, 0.3) and point shapes indicate the four levels of *α*_inter_ (0, 0.05, 0.1, 0.3). **(a)** Intraspecific ratio 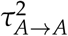, Pearson *r* = 0.995. **(b)** Interspecific ratio 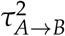, Pearson *r* = 0.999. Panel b covers values from 0 to above 2.0, confirming that the analytical expression remains accurate in the *τ >* 1 regime, where the heritable variance attributable to species *A* exceeds the total phenotypic variance of a population of isolated individuals of species *B* (*n* = 128 scenarios).

**Figure S4.**
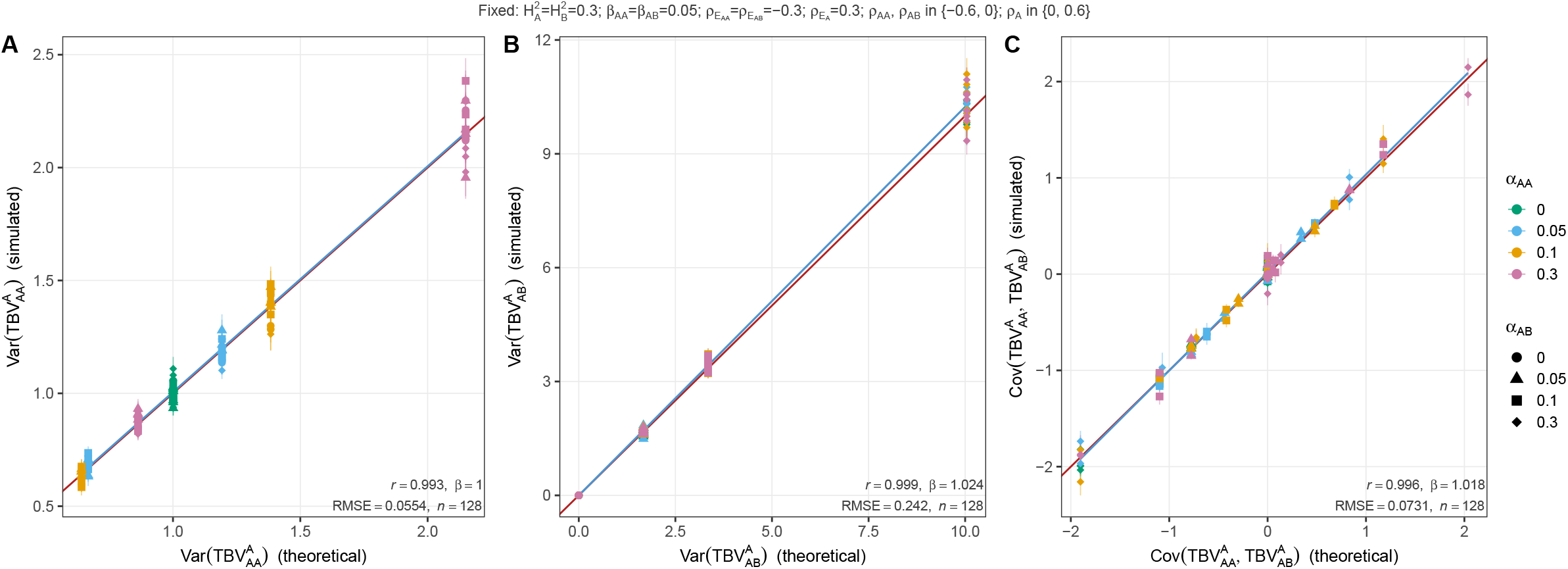
Validation of the analytical expressions for the total breeding value decomposition (Eq. 6) across 128 parameter combinations. Each point represents the mean over 10 simulation replicates for one scenario, with standard error bars on the axes. The red line is the identity (*y* = *x*), and the blue line is the ordinary least-squares regression. Point colors indicate the four levels of *α*_intra_ (0, 0.05, 0.1, 0.3) and point shapes indicate the four levels of *α*_inter_ (0, 0.05, 0.1, 0.3). **(a)** Intraspecific TBV variance 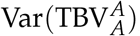, Pearson *r* = 0.993. **(b)** Interspecific TBV variance 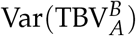, Pearson *r* = 0.999. **(c)** Covariance 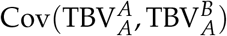, Pearson *r* = 0.996. Panel c spans negative values, confirming that the analytical expression correctly captures trade-off configurations (*n* = 128 scenarios).

**Figure S5.**
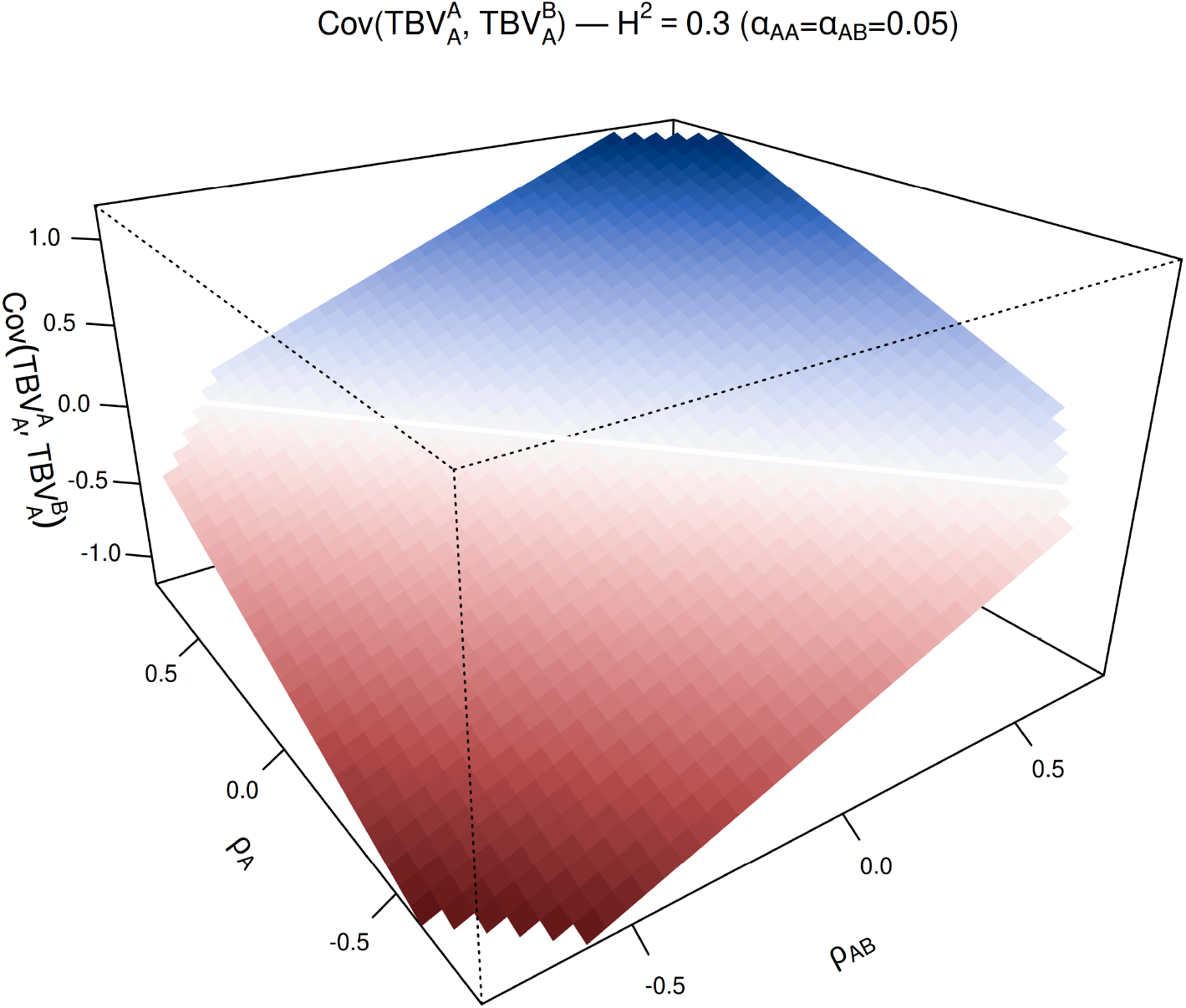
Covariance between the intraspecific 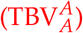 and interspecific 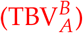 total breeding values of species *A*, as a function of the DGE-interspecific IGE correlation *ρ*_*AB*_ and the correlation between the two indirect genetic effects of species *A, ρ*_*A*_, with the variance ratios fixed at *α*_*AA*_ = *α*_*AB*_ = 0.05 and *n* = 4 neighbors of each type. The surface is computed from the analytical expression (Eq. 6), which was validated against simulations in Figure S4. It varies linearly with *ρ*_*AB*_, so the two total breeding values of species *A* become correlated as soon as *ρ*_*AB*_*≠* 0. This is in direct contrast with the interspecific relative heritable variance ratio 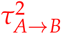 (Figure 3b), which does not depend on *ρ*_*AB*_. The surface does not depend on heritability, since the total breeding value is a purely genetic quantity, and the panel shown is for 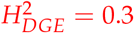. The white line marks the zero-covariance boundary, and combinations of *ρ*_*AB*_ and *ρ*_*A*_ that do not yield a positive-definite genetic correlation matrix are left blank.

### Upper bound on the accuracy of the estimated genetic effects

The accuracy reported in this study is the Pearson correlation *r*(*u, û*) between a true simulated effect and its BLUP. This correlation cannot reach one in a finite experiment, and the value it can reach depends on how much information the design carries on the effect concerned. This section derives that upper bound, so that the plateaus observed in the accuracy panels can be read against it.

#### The bound for a single effect

For any BLUP, the squared correlation with the true effect equals its reliability (Henderson 1975):

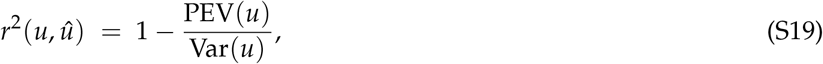

where PEV(*u*) = Var(*u* − *û*) is the prediction error variance. Consider one effect *u* with variance 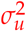, informed by *m* records, each of which also carries an independent disturbance of variance 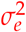 gathering everything else that acts on that record. The best predictor of *u* from the mean of its *m* records is the regression of *u* on that mean, whose reliability is

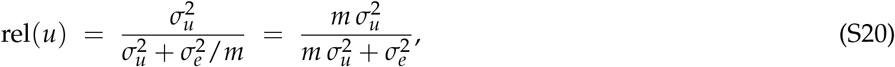

so that 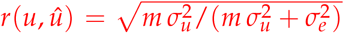. The bound therefore depends on two quantities only: how much variance the effect carries relative to the noise, and how many records inform it. It is strictly below one for any finite *m* and reaches one only as *m* → ∞ or 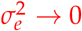.

#### Application to the direct genetic effect

For a direct genetic effect, the *m* records are the *n*_*rep*_ plants carrying the genotype. Writing the individual heritability as *h*^2^ = Var(*DGE*)/Var(*P*) and taking Var(*P*) as the unit, we have 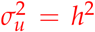 and 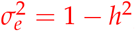, so that Equation S20 becomes

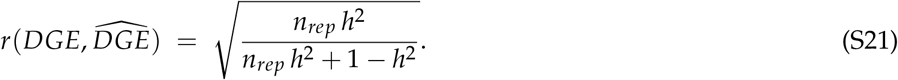

This is the classical accuracy of a genotype mean (**?**), and it is the reason why the accuracy panels plateau below one: with *n*_*rep*_ = 10 the bound is 0.73 at *h*^2^ = 0.1 and still only 0.98 at *h*^2^ = 0.7.

#### Application to the indirect genetic effects

An indirect genetic effect differs in both quantities. Its variance is smaller, by the ratio *α* = Var(*IGE*)/Var(*DGE*) explored in the simulations, so 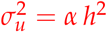. But it is informed by more records: each of the *n*_*rep*_ plants carrying the genotype transmits its indirect effect to each of its *n*_*nb*_ neighbors of the species concerned, so the effect appears in

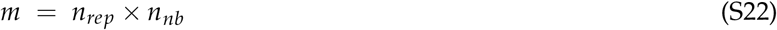

phenotypic records instead of *n*_*rep*_. In the design simulated here *n*_*nb*_ = 2 for the intraspecific effect and *n*_*nb*_ = 6 for the interspecific effect. Substituting into Equation S20 gives

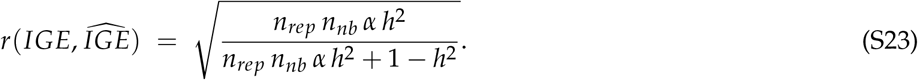

The two changes act in opposite directions. The factor *α*, of the order of 0.05 to 0.10, lowers the bound; the factor *n*_*nb*_ raises it. Their product decides, which is why the interspecific effect, seen by six neighbors, reaches a higher ceiling than the intraspecific effect, seen by two, even though both have the same variance ratio. This is the same asymmetry that appears in the estimation accuracies, and it is a property of the design rather than of the model.

**Table S1.** Upper bound of Equations S21 and S23 at *n*_*rep*_ = 10 and *α* = 0.05, compared with the accuracies obtained in the simulations. The bound is approached but never exceeded.

| $h^2$ | 0.1 | 0.3 | 0.5 | 0.7 |
| --- | --- | --- | --- | --- |
| DGE, bound | 0.73 | 0.90 | 0.95 | 0.98 |
| DGE, simulated | 0.73 | 0.90 | 0.95 | 0.97 |
| Interspecific IGE, bound | 0.50 | 0.75 | 0.87 | 0.94 |
| Interspecific IGE, simulated | 0.50 | 0.74 | 0.84 | 0.91 |

#### Why the bound is an upper bound

Equation S20 treats the *m* records as independent and everything other than the effect of interest as uncorrelated noise. Neither holds exactly. A plant is a neighbor to several focal plants, so the records informing one indirect effect share phenotypes and are not independent, and several indirect effects act on the same phenotype, so part of what the bound counts as noise is itself structured. Both make the real accuracy fall slightly below the bound, which is what Table S1 shows at high heritability. The joint model works in the other direction, since the DGE-IGE covariance lets the well-estimated direct effect carry information to the indirect ones, but this does not overturn the bound: it is why the simulated values stay close to it rather than well below.

### Effect of the genetic correlations on estimation

To examine the effects of the correlations between IGEs and DGE and between both IGEs, we fixed the following parameters: 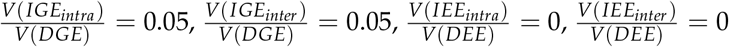.

**Figure S6.**
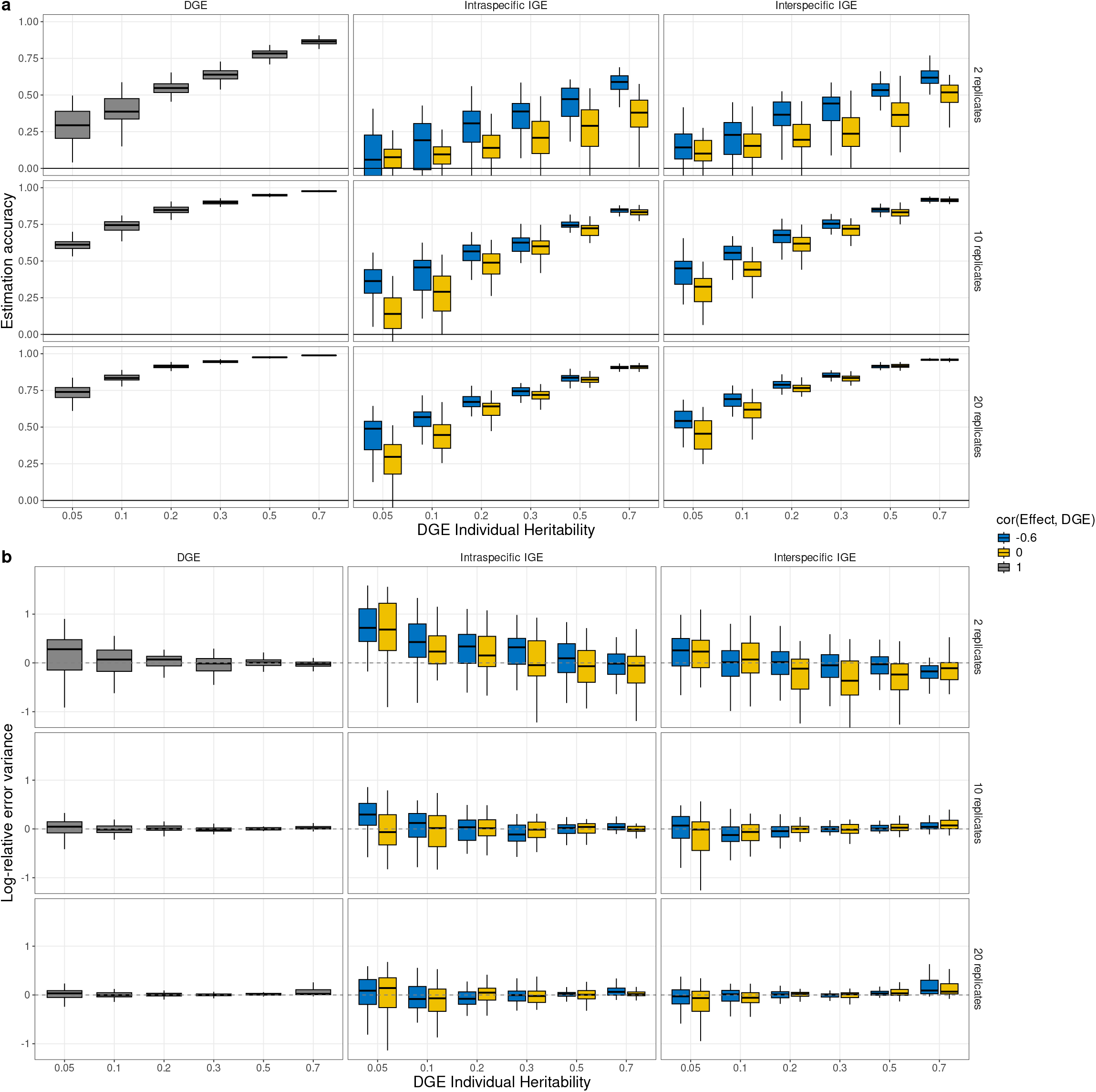
Estimation accuracies (a) and errors (b) obtained by simulating two interacting species in the case of varying correlations between direct and indirect genetic effects. DGE, direct genetic effect; IGE, indirect genetic effect; IEE: Indirect environmental effect. DGE heritability is defined as V(DGE)/V(Pheno). The simulated data follow the model given in Equation. 11 (see text). **(a)** correlations between estimated and true genetic values of the different genetic effects DGE, IGE intra and IGE Inter according to the heritability and the number of replicates per genotype, **(b)** the logarithm of the ratio of the estimated over true variance of each genetic effect. Colors indicate the correlation between IGE and DGE. In column DGE, both r(IGEinter,DGE) and r(IGEintra,DGE) are fixed to -0.6. In column IGE inter, r(IGEintra,DGE) is fixed to -0.6. Reciprocally, in column IGE, r(IGEinter,DGE) is fixed to -0.6. The correlation r(IGEinter,IGEintra) was always fixed at 0.6. All IGE over DGE variance ratios are fixed at 0.05. No interacting environmental effects were simulated (v(IEE)=0). Each boxplot represents up to 50 independent simulations with 100 genotypes per species, runs that did not return a complete genetic covariance matrix being excluded (Section Model fitting and Estimation accuracies). For clarity, outlier points were removed from the graph.

For both IGEs, the estimation accuracy increased with the absolute value of the DGE–IGE covariance (Figure S6 (a)), and this improvement was more pronounced for the intraspecific IGE than for the interspecific IGE. This pattern is a direct consequence of the joint multivariate estimation performed by the model: because DGE is the best-estimated effect in all scenarios, a non-null covariance between DGE and an IGE effectively transfers information from the well-identified DGE to the less-identified IGE, reducing its estimation error (Henderson and Quaas 1976). Intraspecific IGE benefits more from this information transfer than interspecific IGE because it is intrinsically harder to estimate, and its signal relies on fewer intraspecific neighbors 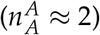 than interspecific ones 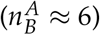; therefore, the marginal gain from exploiting the DGE–IGE covariance is larger. Importantly, *V*_*error*_ was not affected by changes in the DGE–IGE covariance (Figure S6 (b)), indicating that while the covariance structure improves the ranking of individuals (accuracy), it does not correct the overestimation of the variance components.

**Figure S7.**
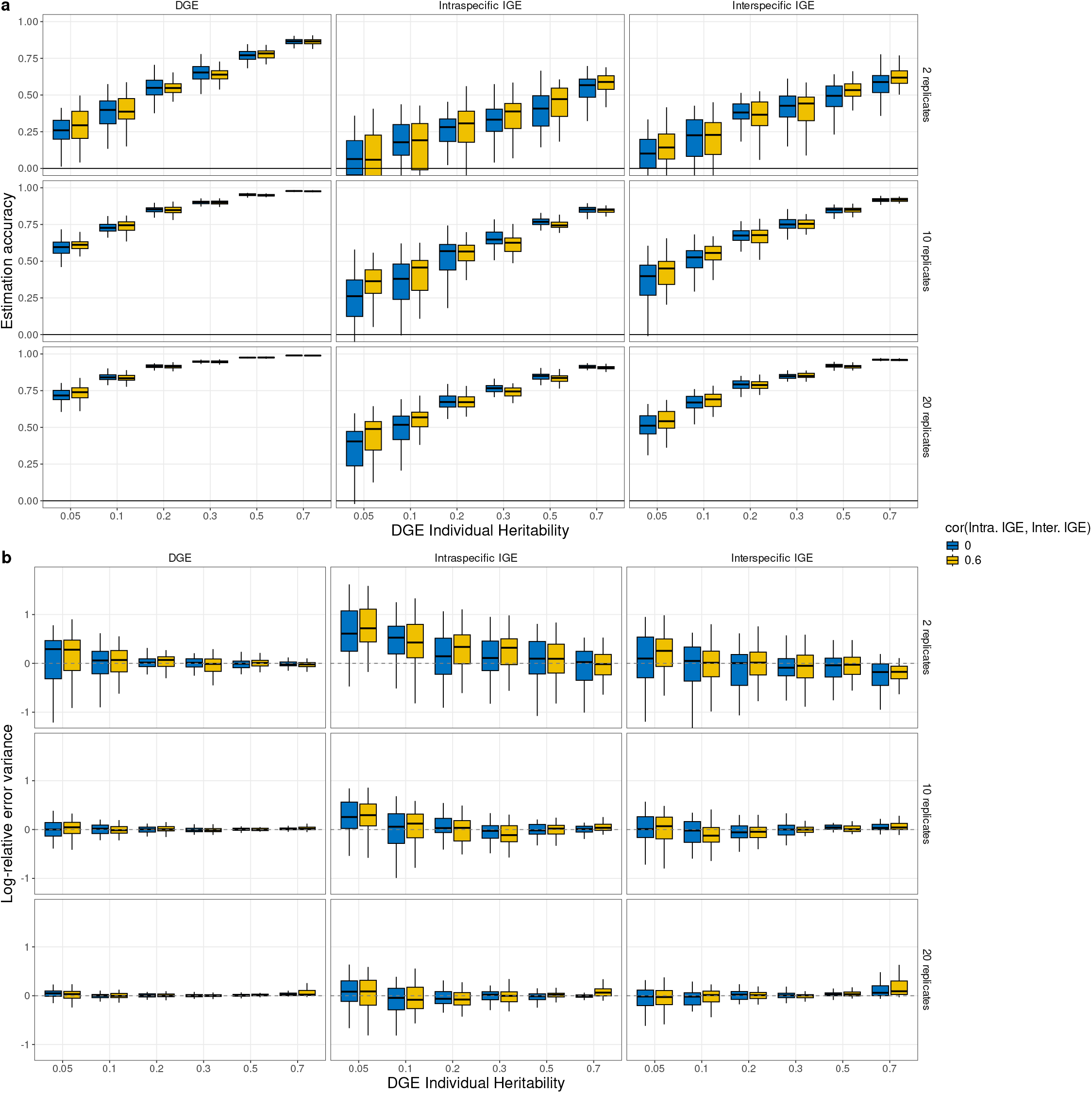
Estimation accuracies (a) and errors (b) obtained by simulating two interacting species in the case of varying correlations between both indirect genetic effects. DGE: direct genetic effect, IGE: indirect genetic effect DGE heritability is V(DGE)/V(Pheno), IEE: Indirect environmental effect. The simulated data follow the model given in Eq. 11 (see text). **(a)** correlations between estimated and true genetic values of the different genetic effects DGE, IGE intra and IGE Inter according to the heritability and the number of replicates per genotype, **(b)** the logarithm of the ratio of the estimated over true variance of each genetic effect. Colors indicate the correlation between intraspecific and interspecific IGEs. Both r(IGEinter,DGE) and r(IGEintra,DGE) were fixed at -0.6. All the IGE over DGE variances ratios are fixed to 0.05. No interacting environmental effect were simulated (v(IEE)=0). Each boxplot represents up to 50 independent simulations with 100 genotypes per species, runs that did not return a complete genetic covariance matrix being excluded (Section Model fitting and Estimation accuracies). For clarity, outlier points were removed from the graph.

The covariance between intraspecific and interspecific IGE did not have a meaningful impact on prediction accuracy (Figure S7 (a)). This more limited effect is consistent with the fact that neither IGE is as well-estimated as the DGE, so the mutual information gain between the two IGEs is smaller than that obtained through their covariance with the DGE. As with the DGE-IGE covariance, *V*_*error*_ remained unaffected by the IGE-IGE covariance (Figure S7 (b)), confirming that the covariance structure influences individual ranking but not the variance component estimation.

**Figure S8.**
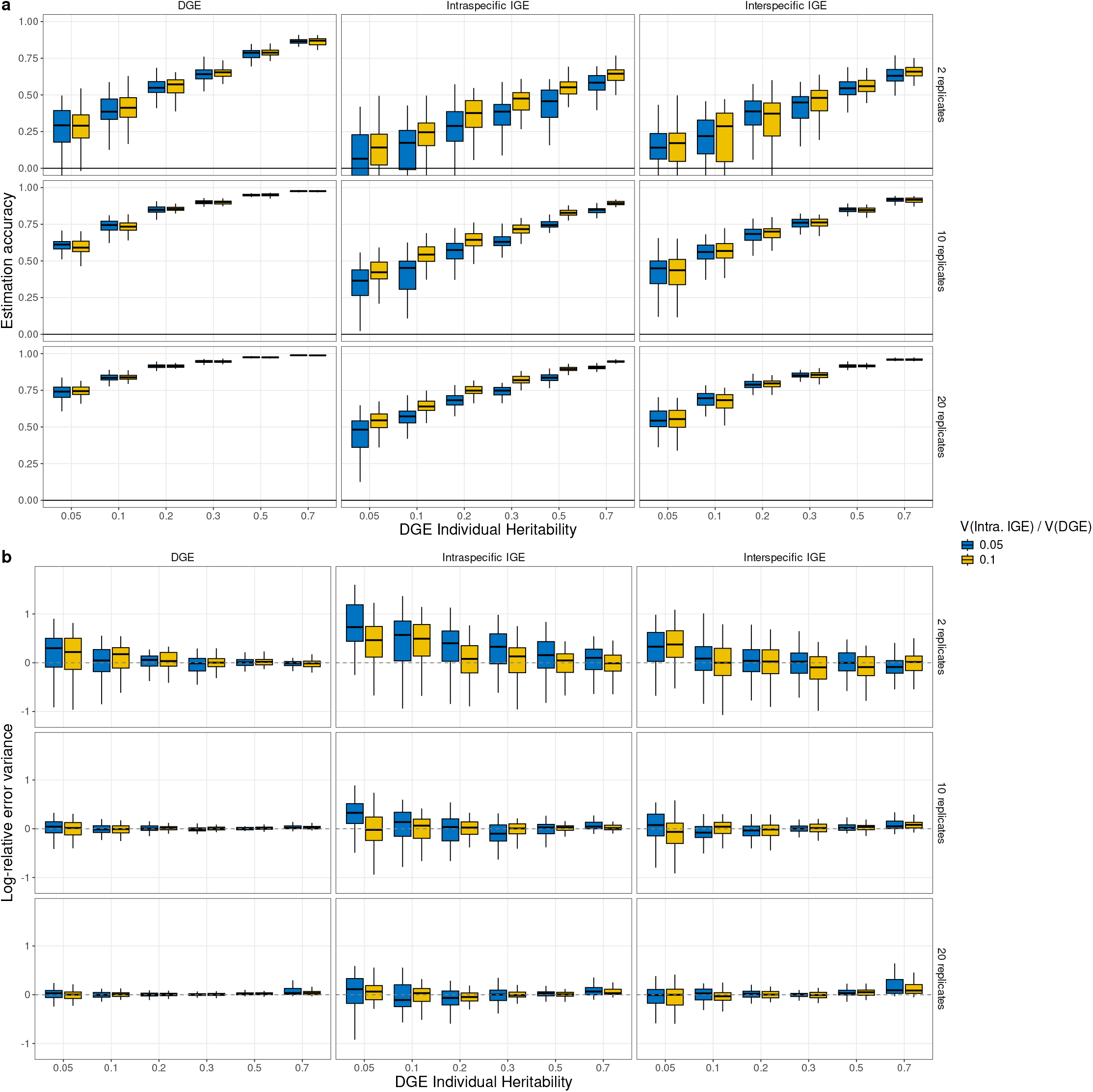
Estimation accuracies (a) and errors (b) obtained by simulating two interacting species in the case of varying levels of plant-plant intraspecific interactions. DGE : direct genetic effect, IGE : indirect genetic effect, IEE: Indirect environmental effect. DGE heritability is V(DGE)/V(Pheno). The simulated data follow the model given in Eq. 11 (see text). **(a)** correlations between estimated and true genetic values of the different genetic effects DGE, IGE intra and IGE Inter according to the heritability and the number of replicates per genotype, **(b)** the logarithm of the ratio of the estimated over true variance of each genetic effect. The colors indicate the relative ratio of V(IGEintra)/V(DGE). For all columns, V(IGEinter)/V(DGE) was fixed at 0.05. Correlations between all effects were fixed: r(DGE,IGEintra)=-0.6, r(DGE,IGEinter)=-0.6, r(IGEintra,IGEinter)=0.6. No interacting environmental effect were simulated (v(IEE)=0). Each boxplot represents up to 50 independent simulations with 100 genotypes per species, runs that did not return a complete genetic covariance matrix being excluded (Section Model fitting and Estimation accuracies). For clarity, outlier points were removed from the graph.

**Figure S9.**
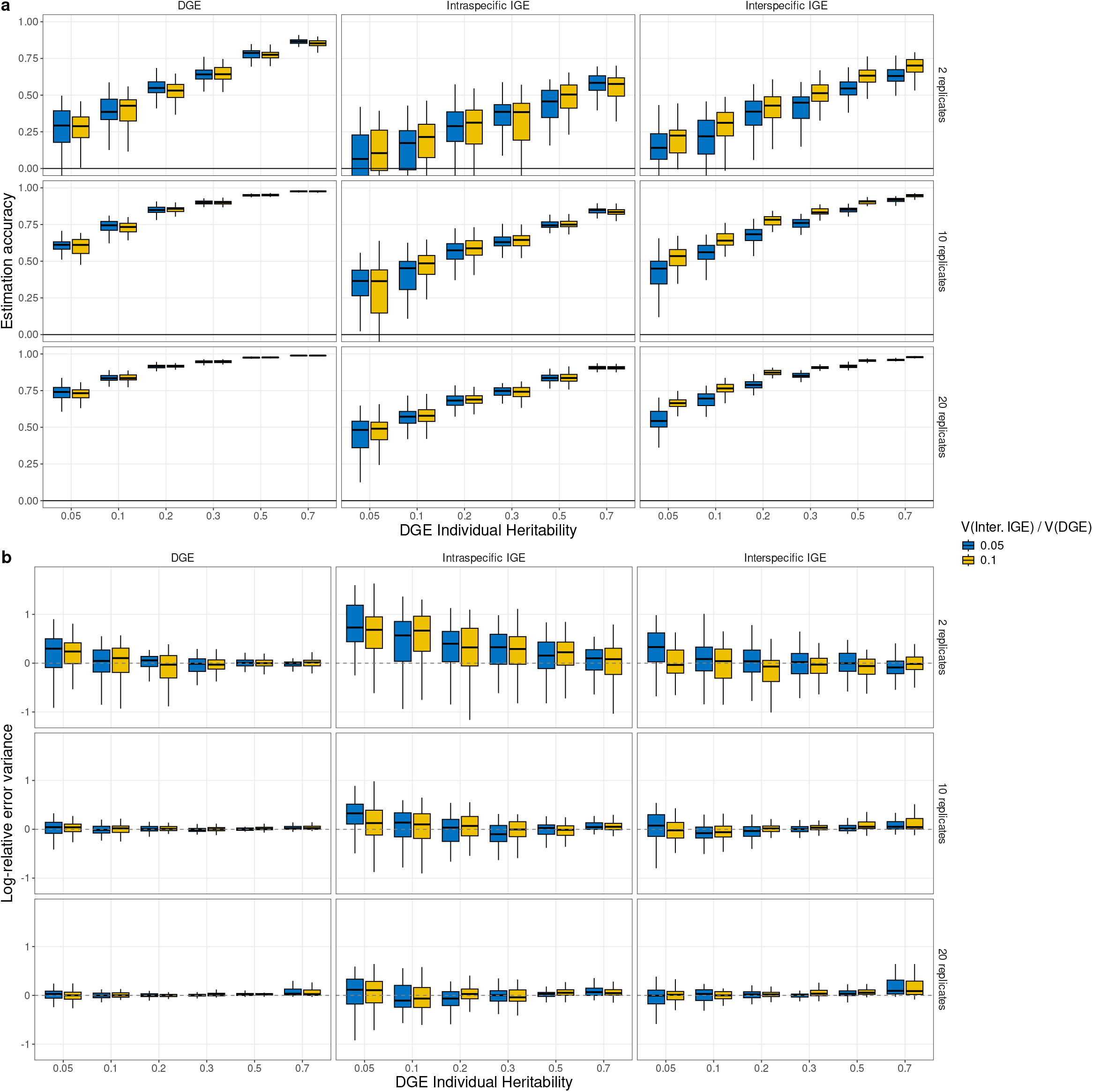
Estimation accuracies (a) and errors (b) obtained by simulating two interacting species in the case of varying levels of plant-plant intraspecific interactions. DGE : direct genetic effect, IGE : indirect genetic effect, IEE: Indirect environmental effect. DGE heritability is V(DGE)/V(Pheno). The simulated data follow the model given in Eq. 11 (see text). **(a)** correlations between estimated and true genetic values of the different genetic effects DGE, IGE intra and IGE Inter according to the heritability and the number of replicates per genotype, **(b)** the logarithm of the ratio of the estimated over true variance of each genetic effect. Colors indicate the relative ratio of V(IGEinter)/V(DGE). For all columns, V(IGEintra)/V(DGE) was fixed at 0.05. Correlations between all effects were fixed: r(DGE,IGEintra)=-0.6, r(DGE,IGEinter)=-0.6, r(IGEintra,IGEinter)=0.6. No interacting environmental effect were simulated (v(IEE)=0). Each boxplot represents up to 50 independent simulations with 100 genotypes per species, runs that did not return a complete genetic covariance matrix being excluded (Section Model fitting and Estimation accuracies). For clarity, outlier points were removed from the graph.

**Figure S10.**
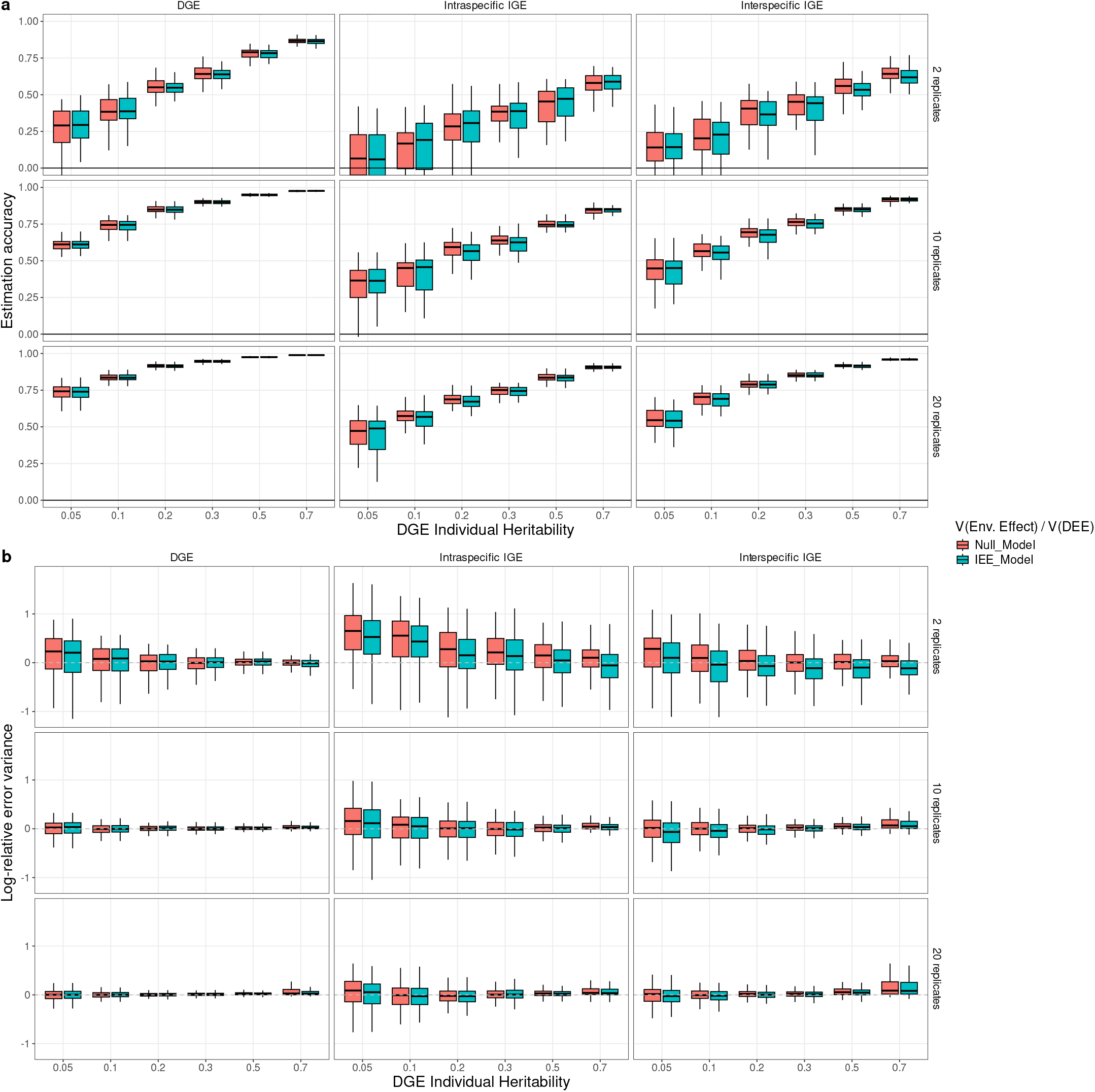
Estimation accuracies (a) and errors (b) obtained by two different models accounting explicitly or not for IEE on simulation of two interacting species. DGE, direct genetic effect; IGE, indirect genetic effect; DGE heritability is V(DGE)/V(Pheno); IEE: Indirect environmental effect. The simulated data follow the model given in Eq. 11 (see text). **(a)** correlations between estimated and true genetic values of the different genetic effects DGE, IGE intra and IGE Inter according to the heritability and the number of replicates per genotype, **(b)** the logarithm of the ratio of the estimated over true variance of each genetic effect. The colors indicate the model used for inference. The IEE model corresponds to the model presented in 13 and the null model corresponds to the same model without the IEE-related terms. All pairwise correlations between IGE and DGE are fixed at -0.6. The correlation between both IGE is fixed at 0.6. All the IGE over DGE variances ratios are fixed to 0.05. No indirect environmental effects were simulated (V(IEE)=0). Each boxplot represents up to 50 independent simulations with 100 genotypes per species, runs that did not return a complete genetic covariance matrix being excluded (Section Model fitting and Estimation accuracies). For clarity, outlier points were removed from the graph. The effect of the choice of the estimation model on the accuracy and the log-relative error of the variance in absence of IEEs is represented in S10

### Variance ratios cross effects

### Design comparison at normalized exposure

At a dilution factor of zero a plant of the continuous grid receives the sum of eight neighbor contributions where a potted plant receives one (Figure 8), so the advantage of the continuous neighborhood combines two mechanisms, the number of distinct partners a genotype grow in the vicinity of and the amount of neighborhood signal each plant receives. To separate them we repeated the comparison at a dilution factor of 1/2, which rescales the neighborhood weights so that every plant is exposed to the same total indirect variance whatever the number of neighbors it has, the two designs being then on an equal footing in that respect (Figure S11).

The direct genetic effect is unchanged, as expected, being informed by the phenotype of the plant itself rather than by its neighborhood. The two indirect effects lose accuracy in the continuous neighborhood, which is the part of the advantage that came from exposure, and the pairwise design is unaffected, its plants having one neighbor under either setting. The ranking nonetheless holds at every size: the continuous neighborhood reaches a given accuracy on either indirect effect with about half as many plants as the pairwise design, and the gap persists at a hundred thousand plants. What a continuous neighborhood buys is therefore not only that its plants are more exposed, but that the same plants realize many more genotype combinations, and only the second of these survives normalization.

**Figure S11.**
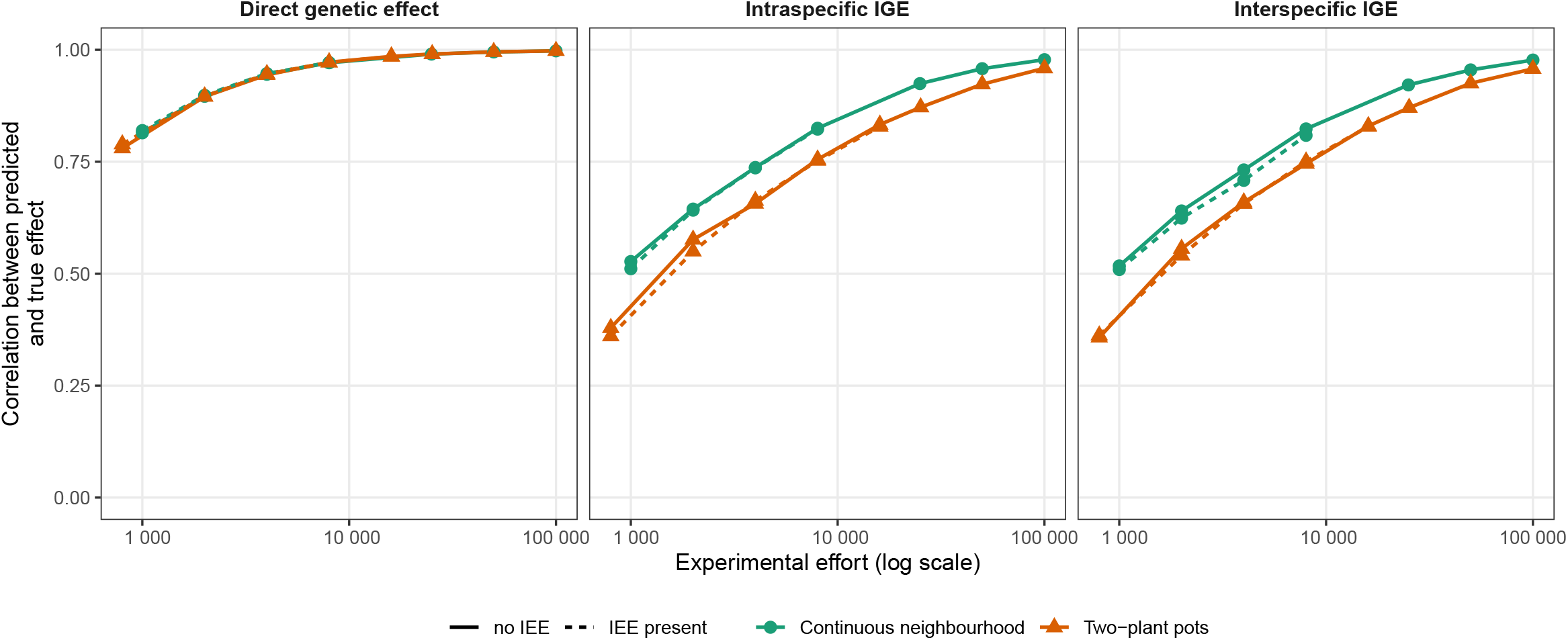
Accuracy of the predicted genetic effects for the two designs at a dilution factor of 1/2, at which every plant is exposed to the same total indirect variance whatever its number of neighbors. Accuracy is the correlation between the predicted and the true genetic effects, plotted against the number of plants grown, which is also the number of phenotypic measurements, every plant being measured in both designs. Solid lines: no indirect environmental effect. Dashed lines: indirect environmental effects present in the data, and fitted in the continuous neighborhood, the only design in which they are estimable. Simulation parameters, identical for both species: 100 genotypes per species, 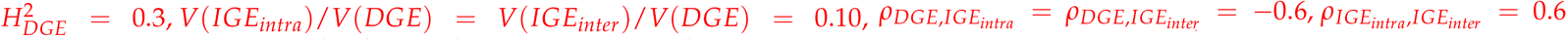, and, where indirect environmental effects are present, *V*(*IEE*_*intra*_)/*V*(*DEE*) = *V*(*IEE*_*inter*_)/*V*(*DEE*) = 0.1. No genotype-by-genotype interaction is simulated. The genetic effects are fitted with an unstructured covariance matrix. Each point is the median of 40 independent simulations.

### Neighborhood order in the semi controlled experiment

The order of the neighborhood used to analyse the semi-controlled experiment was chosen on the data. Models that differ only in *Zn* estimate the same number of parameters, so they are comparable by AIC. Ten ranks were fitted, each of them reporting the same parameterisation of Equation 13, so that the series of variance components can be read from one order to the next. The AIC falls steeply up to order 6 and only drifts afterwards: rank 6 already captures 97 % of the total decrease, and the four higher ranks add 2.3 points in all, for a total range of 68 points (Figure S12a). This is the counterpart of the geometric saturation of the exposure: an rank-5 neighborhood extends over six ranks of plants but amounts to between 5.8 and 7.7 adjacent-contact equivalents, depending on the species and the interaction type, against 1.9 and 3.7 at rank 1. We therefore retained rank 6 rather than the rank of lowest AIC, which lies 2.3 points below it at rank 10.

The variance components are stable over the whole range (Figure S12b), so the conclusions drawn on the semi-controlled experiment do not rest on that choice. The direct genetic effect stays between 8.3 and 8.7 % of the phenotypic variance in wheat and between 7.6 and 8.5 % in alfalfa, and the effect of alfalfa on wheat between 2.6 and 4.4 % from rank 3 onwards.

**Figure S12.**
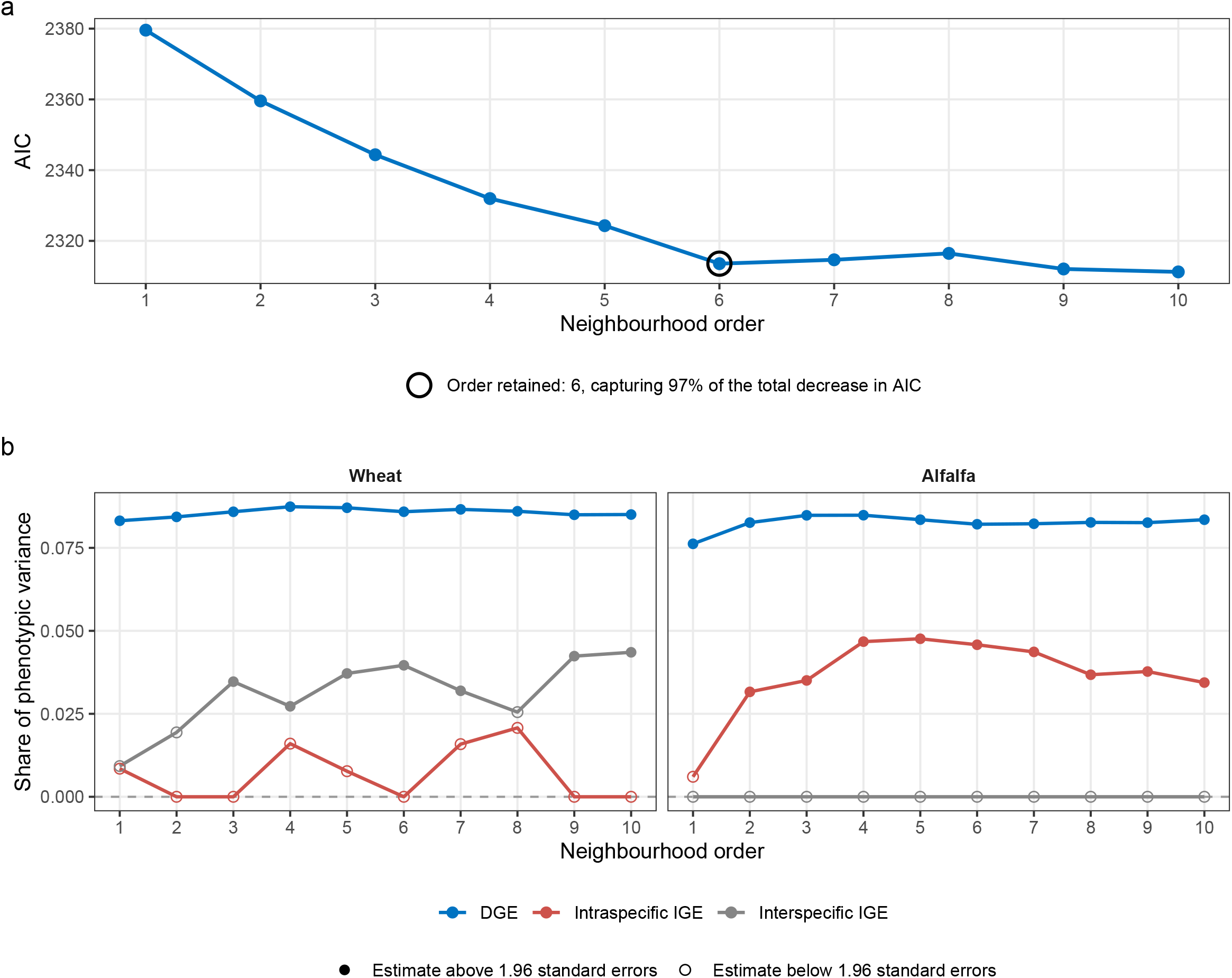
Effect of the neighborhood rank on the analysis of the wheat-alfalfa semi-controlled experiment. **(a)** AIC of the reported model against the rank of the neighborhood, the circled point being the rank retained. **(b)** variance components of the same model, as a share of the phenotypic variance of each species, against the rank of the neighborhood. DGE, direct genetic effect; IGE, indirect genetic effect. Filled points mark the estimates that exceed 1.96 standard errors.

